# Contrasting impacts of invasive *Opuntia* cacti on mammal habitat use

**DOI:** 10.1101/2024.12.12.627951

**Authors:** Peter S. Stewart, Russell A. Hill, Ayub M. O. Oduor, Philip A. Stephens, Mark J. Whittingham, Wayne Dawson

## Abstract

Biological invasions impact ecosystems worldwide, including through changing the behaviour of native species. Here, we used camera traps to investigate the effects of invasive *Opuntia* cacti on the habitat use of twelve mammal species in Laikipia County, Kenya, an internationally important region of mammalian biodiversity. We found that *Opuntia* impacted mammal occupancy and activity, but the strength and direction of the effects varied among species and between seasons, and depended on the spatial scale at which *Opuntia* was considered. Notably, we observed consistent positive effects for olive baboons and elephants, two major consumers of *Opuntia* fruit. We also observed seasonally varying effects on the occupancy of two key grazers: Grevy’s zebra and plains zebra. As well as having important implications for mammal conservation, ecosystem functioning, and the future spread of *Opuntia*, our findings highlight behavioural changes in large mammals as a potentially important pathway through which invasive species impact ecosystems.

## Introduction

Biological invasions are a rapidly expanding threat to ecosystems worldwide (Roy *et al*. 2024). Understanding the impacts of invasive species is vital if we are to mitigate them effectively but, until recently, attention has focused on impacts on biodiversity and native species’ abundance (Crystal-Ornelas & Lockwood 2020). However, a growing body of evidence now indicates that invasive species can cause profound ecological impacts by altering the behaviour of native animals (Langkilde *et al*. 2017; Stewart *et al*. 2021). Developing a mechanistic understanding of these behavioural impacts is an important focus of research in invasion ecology (Stewart *et al*. 2021).

Large mammals can be an important component of terrestrial ecosystems (Pringle *et al*. 2023; Ripple *et al*. 2014). Large mammalian herbivores structure the composition and trait distributions of plant communities (Boulanger *et al*. 2018; Dantas & Pausas 2020; Jia *et al*. 2018), disperse nutrients (le Roux *et al*. 2020) and seeds (Campos-Arceiz & Blake 2011), and regulate fire regimes (Karp *et al*. 2024), while large mammalian carnivores play a key role in controlling herbivore and mesopredator populations, with indirect effects on a range of ecosystem processes (Ripple *et al*. 2014; Ritchie & Johnson 2009). Importantly, behaviour can moderate the ecological effects of large mammals (Pringle *et al*. 2023). Consequently, changes to large mammal behaviour caused by invasive species may constitute an important, yet underappreciated, impact pathway for biological invasions.

Laikipia County, Kenya, is a key stronghold for mammalian biodiversity and hosts vital populations of endangered mammals including Grevy’s zebra (*Equus grevyi*; Rubenstein *et al*. 2016) and reticulated giraffe (*Giraffa reticulata*; Muneza *et al*. 2018). However, the region is also undergoing invasion by prickly pear cacti (*Opuntia spp*.). Native to the Americas, several species of *Opuntia* were introduced to Laikipia County, Kenya, in the latter half of the 20^th^ century to serve as live fences and ornamental plants (Loisaba Conservancy 2019; Strum *et al*. 2015; Witt 2017). Since their introduction, three of these species (*O. stricta*, *O. engelmannii*, and *O. ficus-indica*) have become invasive, spreading rapidly to cover large areas of the landscape (Githae 2019; Strum *et al*. 2015; Witt 2017).

*Opuntia* invasions could alter mammal behaviour through two mutually non-exclusive pathways. First, *Opuntia* substantially alters the physical structure of the habitat by forming dense, impenetrable stands (Witt 2017). These stands may disrupt sightlines and restrict movement, altering patterns of actual or perceived predation risk. *Opuntia* stands may also impede herbivores’ access to forage (Oduor *et al*. 2018); this effect is likely to be especially pronounced in drought conditions, when the overall availability of forage decreases (Boutton *et al*. 1988). Second, mature *Opuntia* stands provide a year-round supply of fruit which is consumed by species including elephants (*Loxodonta africana*), olive baboons, (*Papio anubis*), and vervet monkeys (*Chlorocebus pygerythrus*; Githae 2019; Strum *et al*. 2015; Witt 2017). Consequently, these species may be attracted to invaded areas (Shackleton *et al*. 2017), which could increase seed dispersal and accelerate the invasion.

Given the ecological importance of large mammals in Laikipia and the surrounding counties (Pringle *et al*. 2010), understanding the behavioural impacts of *Opuntia* on these species is of particular importance. Furthermore, as *Opuntia* species are also invasive in other mammal-rich regions across the world (Foxcroft *et al*. 2004, Pasiecznik & Rojas-Sandoval 2007, Pasiecznik 2015), understanding how *Opuntia* affects mammal behaviour is of global significance. Here, we addressed this problem by investigating the behavioural impacts of *Opuntia* on the habitat use of mammals in Laikipia County. Using camera traps and drawing on insights from the field of causal inference, we quantified the total effects of *Opuntia* on the occupancy, total activity, and temporal activity patterns of twelve mammal species which represent a range of trophic guilds and body sizes. As the behavioural impacts of invasive plants can depend on ecological context and spatial scale (Stewart *et al*. 2021), we explored how the effects varied across two seasons and two different scales of *Opuntia* measurement. Finally, to gain further insights into the mechanisms underlying *Opuntia*’s behavioural impacts, we examined whether *Opuntia* exerts indirect effects on habitat use through changing the cover of native plant species.

## Methods

### Study system

We conducted our study at Mpala Research Centre and Loisaba Conservancy, in Laikipia County, Kenya (Fig. 1). The study area is predominantly unfenced (Crego *et al*. 2021) semi-arid savanna, with varying densities of woodland and shrubland dominated by *Vachellia* and *Senegalia* (formerly *Acacia*) species, including *V. etbaica, S. brevispica, S. mellifera, and V. gerrardii*, alongside other species including *Boscia augustifolia*, *Croton dichogamus,* and *Grewia spp.* (Augustine 2003; Augustine *et al*. 2011; Mutuku & Kenfack 2019; Young *et al*. 1995). Grasses belonging to the genera *Cynodon*, *Pennisetum*, *Digitaria* and *Sporobolus* are common in the understory, and a variety of forbs including *Plecranthus spp*., *Pollichia campestris*, *Portulaca spp*. and *Blepharis spp*. are also present (Young *et al*. 1995). *Euphorbia nyikae* and other succulents also occur in some areas (Augustine 2003).

**Figure 1.**
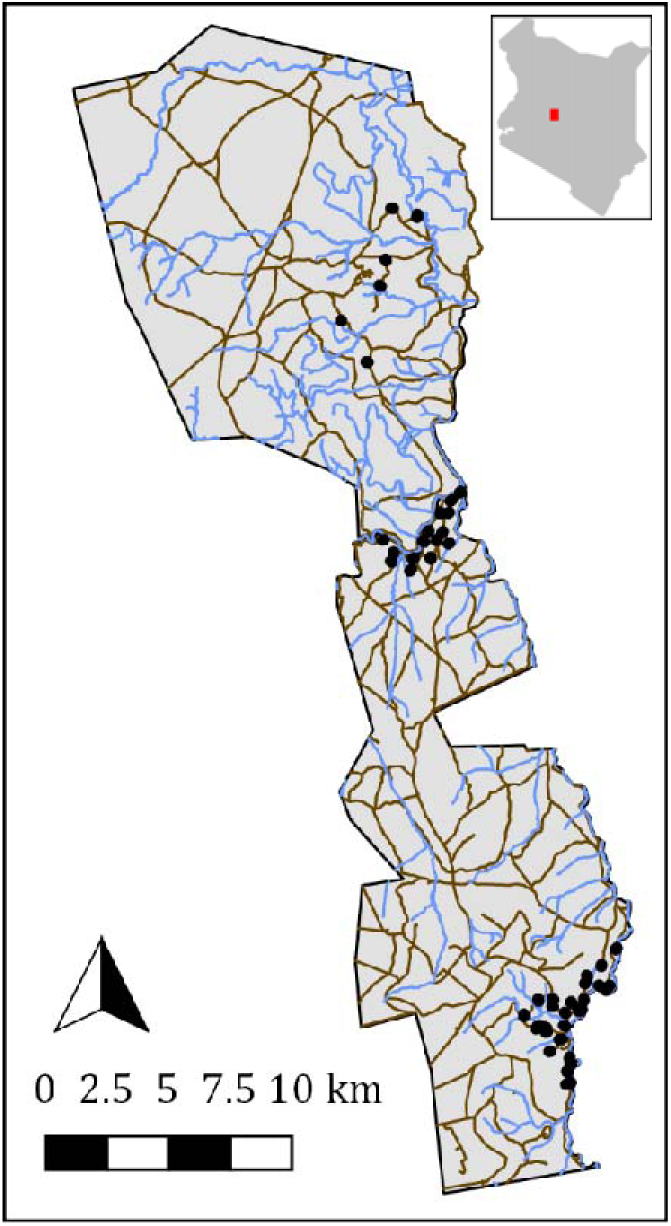
Map of the study area. Black points represent camera trap sites, light blue lines are rivers, and brown lines are roads. Camera trap sites are clustered within three regions: Loisaba (top), Mpala north (middle) and Mpala south (bottom). Canvas extends from longitude 244165 to 271026, latitude 28814 to 78171. Inset shows location of the study region within Kenya.

We focused on twelve mammal species. Olive baboons (*P. anubis*), vervet monkeys (*C. pygerythrus*), and elephants (*L. africana*) were selected because they commonly feed on *Opuntia* fruit and are presumed to be important dispersal agents (Githae 2019; Strum *et al*. 2015; Witt 2017). Buffalo (*Syncerus caffer*), dik-dik (*Madoqua spp.*), impala (*A. melampus*), greater kudu (*Tragelaphus strepsiceros*), reticulated giraffe (*G. reticulata*), Grevy’s zebra (*E. grevyi*), and plains zebra (*E. quagga*) were selected to represent a range of body sizes and diet compositions (Kartzinel *et al*. 2015) across herbivore species which are not known to feed on *Opuntia* fruit. Finally, we included two carnivore species: spotted hyena (*Crocuta crocuta*) and leopard (*Panthera pardus*).

### Camera trap deployment

To explore the effects of *Opuntia* on mammals’ occupancy and activity, we deployed camera traps in three sub-regions of the study area (Fig. 1) from January-April and October-November 2021. We selected the sub-regions because they contained varying densities of *Opuntia*, from scattered individual plants to heavily invaded areas where *Opuntia* covered the majority of the ground. To maximise variation in *Opuntia* while minimising variation in confounding variables, we employed a paired-site design, where a site is defined as the area immediately adjacent to one camera trap. Each sub-region was divided into 500 × 500m grid squares, with a subset of squares randomly selected for sampling. We placed two cameras within each square; the first camera was deployed in an area visually identified as high *Opuntia* density, and the second was deployed in a random direction 50-70m away. If the second site was found to have an equal or higher *Opuntia* density than the first site, we generated a new random direction until the density at the second site was lower.

We deployed 30 cameras (20 Browning Dark Ops Pro, 5 Browning Recon Force Extreme, and 5 Reconyx Hyperfire 2); three were lost to damage, leaving 27 operational at the end of the study. Due to these losses, and with some grid squares sampled more than once, our total sample comprised 101 sites within 46 squares for January-April, and 27 sites within 14 squares for October-November. The sites sampled in October-November were a subset of those sampled in January-April. Cameras remained operational at each location for between 3 and 56 days in January-April (Q1 = 18, median = 23, Q3 = 26 days) before being moved to a new location; we aimed to leave cameras in place for at least one week before they were moved, but on a few occasions (n = 5) cameras were moved earlier for logistical reasons. In October-November the cameras were not moved, and instead remained in place for between 33 and 46 days (Q1 = 36, median = 37, Q3 = 37 days).

We mounted the cameras on tree trunks/stumps (average ground-to-lens height = 81cm) and positioned them to ensure good visibility 10m in front of the camera. We set the cameras to take images with a five second delay between captures. For the Browning cameras we used the “long range” infrared flash setting, and used the default “optimised” infrared flash for the Reconyx cameras.

### Habitat surveys

To collect information on site-level variables that could affect occupancy and activity, we conducted habitat surveys in a circular area with 10m radius, centred on the camera. We divided this area into the field of view (FOV), defined as the area in which we could see the camera’s lens, and the rest of the area located beside and behind the camera. Within each zone we estimated the percentage of ground covered by *Opuntia*, grasses, shrubs, forbs, succulents, trees, bare ground, and other cover (*e.g.,* rocks) using a cover estimator chart (Anderson 1986). These percentages were not required to sum to 100%, as vegetation types could grow under/over one another. We then averaged the values for the FOV and non-visible area to obtain a site-level value for each ground cover type. In addition, we counted the number of standing trees (woody plants taller than 2m; shorter woody plants were classed as shrubs). To quantify the use of each site by livestock, we calculated the proportion of days in which livestock were detected by the camera. Finally, we calculated the straight-line distances from each site to the nearest river and road using QGIS (v2.28.25; QGIS Development Team, 2018).

### Estimating grid square-level Opuntia

To estimate the quantity of *Opuntia* in each grid square, we performed distance sampling (Kéry & Royle, 2015). Due to constraints imposed by the COVID-19 pandemic, we only sampled 41 of the 46 squares in which camera traps were deployed. The sampling was conducted on foot along a transect; we looked for *Opuntia* visible from the transect in either direction. When an *Opuntia* stand was observed we recorded the size category (small = <1m, medium = 1-2m, large = >2m height) and measured the distance to the stand. Distances were measured using a tape measure for stands <10m away, and using a laser range finder (Leica Rangemaster CRF 2400-R, accurate to ±1m) for stands 10-80m from the transect. We did not count stands further than 80m away to avoid including stands situated outside the square.

To obtain *Opuntia* volume estimates for each of the 41 squares, we first used a Poisson-binomial multinomial distance sampling model with half-normal detection function (Kéry & Royle, 2015; code adapted from Joseph, 2021) to estimate abundance for each *Opuntia* size class. We then combined the median abundance estimates for different size classes into a single volume estimate by assuming (based on the volume of a hemisphere) that the volume of a large stand (h = 2.5m) was 32.725m^3^, a medium stand (h = 1.5m) was 7.070m^3^, and a small stand (h = 0.5m) was 0.260m^3^. Finally, we divided each square’s volume estimate by the respective transect length (measured using QGIS).

### Processing camera trap images

We used Megadetector (v.4.1.0, Beery *et al*. 2019) to classify images as containing an animal (any species), human, or vehicle. We manually screened all images with probability of ≥0.10 of containing a human or vehicle, discarding all images which contained a human or vehicle and retaining images which contained animals. We also retained all images classed as containing at least one animal with probability ≥0.98.

We uploaded all retained images to the Zooniverse platform (Prickly Pear Project Kenya: https://www.zooniverse.org/projects/peter-dot-stewart/prickly-pear-project-kenya), where members of the public were able to classify the images. Each image was classified by at least 12 volunteers before retirement from the active image pool, except for images classified as “human”, which were immediately retired. We generated consensus classifications for each image using a threshold-based approach, in which at least 66% of the volunteers had to classify the species as being present. We also quantified volunteer agreement by calculating the Shannon entropy (Shannon 1948) of the classification distribution for each image; images with entropy values greater than one were discarded. In cases where an image was classified by an expert (either P.S.S. or the Prickly Pear Project Kenya moderator), the expert classification was accepted regardless of volunteer disagreement.

### Statistical models

We fitted models to explore the effects of *Opuntia* on three aspects of habitat use. First, we used an occupancy model (MacKenzie *et al*. 2002) to explore the effects on occupancy (*i.e.,* the probability that a site is used at least once). Second, we modelled the total number of daily detections using a negative-binomial model. Finally, we used a binomial model to explore temporal shifts in species’ activity by modelling the proportion of detections occurring at night (after dusk and before dawn; times obtained using the *suncalc* package; v.0.5.1; Thieurmel & Elmarhraoui, 2022).

We estimated the total effect (*i.e.,* including both the direct and indirect pathways; Fig. 2A) of *Opuntia* measured at two spatial scales: the site-level *Opuntia* cover, and the grid square-level *Opuntia* volume. In both cases, we estimated separate effects for the January-April and October-November seasons. We selected additional covariates by encoding our causal assumptions about the data-generating process in a directed acyclic graph (Fig. 2A), and selecting variables to satisfy the back-door criterion (Arif & MacNeil 2022; Pearl 1995; Stewart *et al*. 2023). As with the *Opuntia* effect, we estimated separate covariate effects for each season. Furthermore, we fitted two sets of models (Fig. 2B) to examine whether *Opuntia* indirectly affects occupancy and activity through its effects on the native plant community.

**Figure 2.**
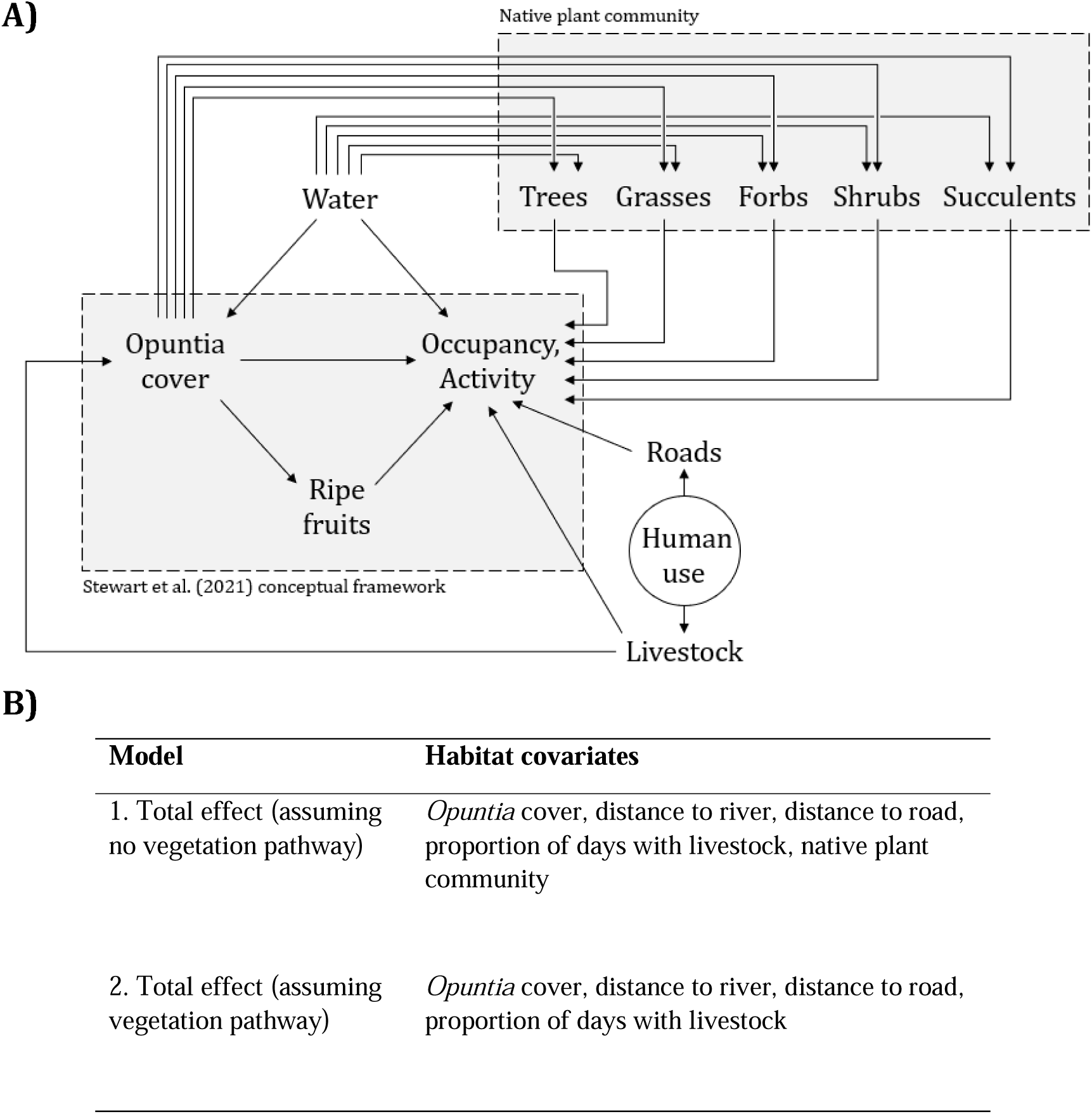
A) Directed acyclic graph representing assumptions about the ways in which *Opuntia* cover and other environmental variables might affect mammal occupancy and activity. Nodes represent variables, while arrows represent possible mechanistic links between variables. “Human use” is a latent variable and is therefore displayed in a circle. **B)** Choice of habitat covariates in our models. “Native plant community” is shorthand for the total percentage cover covariates for grass, shrubs, forbs and succulents, and the total number of trees.

In our occupancy model, we included daily mean temperature in the detection sub-model, as changes to the thermal environment can affect the performance of the camera’s passive infrared sensor (Welbourne *et al*. 2016). We also included an offset intercept for each model of camera trap. In our binomial model, we included an interaction between *Opuntia* and lunar illumination (obtained using the *suncalc* package), to investigate whether moonlight moderated *Opuntia*’s effects (e.g., by altering predation risk; Prugh & Golden 2014). We standardised all covariates, excluding lunar illumination, by subtracting the mean and dividing by the standard deviation. In all models, we incorporated a Gaussian process to model spatial autocorrelation.

We fitted all models in a Bayesian framework using Stan, implemented with CmdStan (v2.33.1; Stan Development Team 2023); all data processing and visualisation was performed in R (v.4.4.2; R Core Team, 2024). We used weakly regularising prior distributions (McElreath 2021, p.214). For each model we used four chains, each with 2000 sampling iterations. We ran 4000 warmup iterations for the occupancy and binomial models, and 7000 for the negative-binomial model. We assessed convergence using the 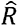 diagnosti (Vehtari *et al*. 2021), inspected the effective sample size for each parameter, and checked for divergent transitions.

## Results

We collected a total of 1,726,954 camera trap images. After retaining images which Megadetector classified as containing at least one animal, and discarding images which contained humans or vehicles, the remaining 186,861 images were uploaded to Zooniverse. We were able to obtain at least one consensus classification for 174,336 of these images; of these, 22,714 images were consensus classified as empty (*i.e.,* Megadetector false positives). For 12,525 of the images uploaded to Zooniverse we were unable to obtain a consensus classification, generally because the animal was too poorly photographed to be identified to species level. As some images contained multiple species, we ultimately obtained 155,630 species detections across 151,622 images. Based on the analysis of 26,952 images which were classified by an expert in addition to the volunteers, our approach resulted in accurate classifications, with sensitivity ≥ 0.973 and specificity ≥ 0.998 for all focal species (Table S1).

### Occupancy

In our occupancy models for olive baboons and elephants, we observed positive effects of *Opuntia* for both seasons and spatial scales (Figs. 3A,C, 4A,C). By contrast, the effect for vervet monkeys was only positive for site-level *Opuntia* in January-April; in October-November we observed no clear effect, and for grid square *Opuntia* the estimated effect was negative in both seasons.

**Figure 3.**
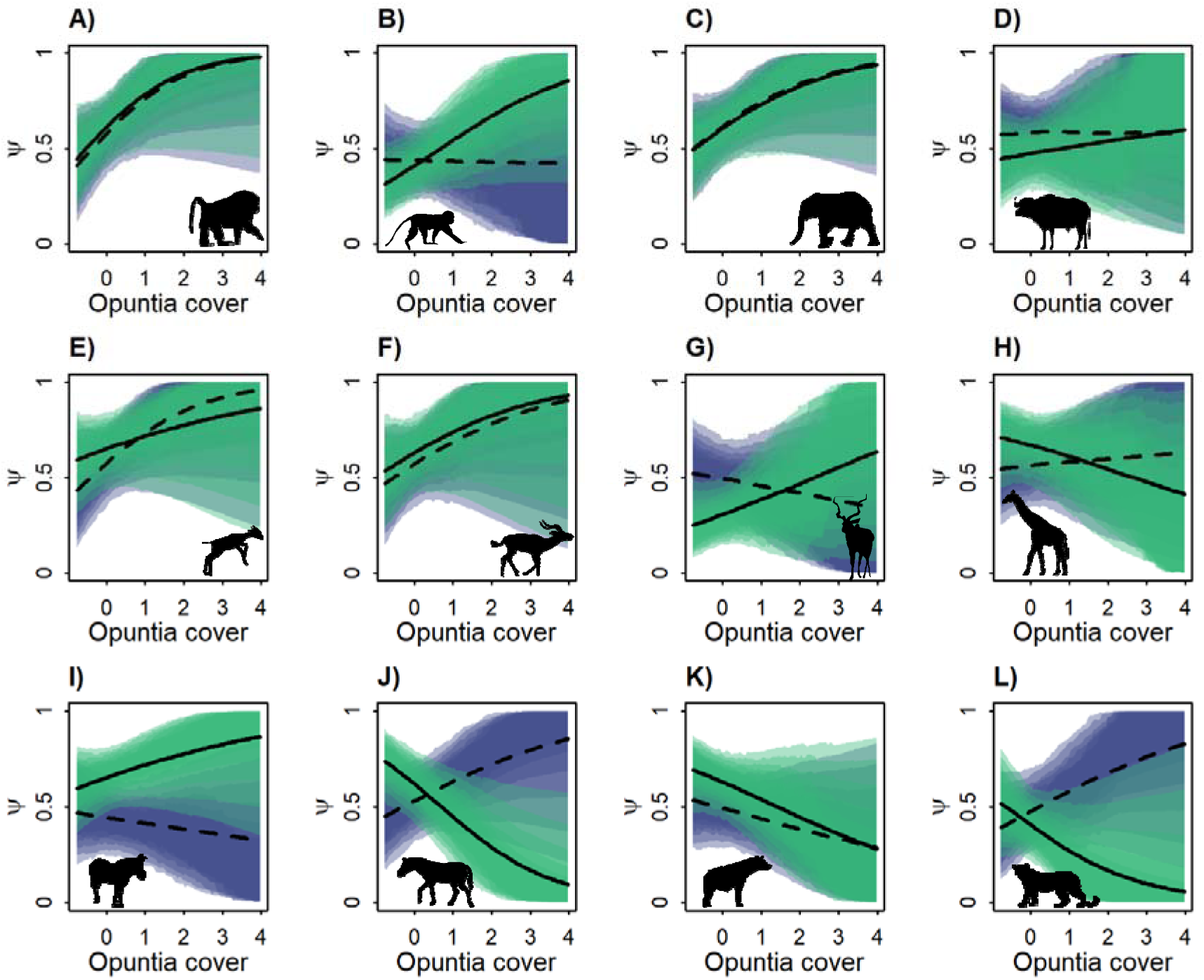
Marginal total effect of site-level *Opuntia* percentage cover (standardised) on occupancy probability (ψ) for: **A)** olive baboon, **B)** vervet monkey, **C)** elephant, **D)** buffalo, **E)** dik-dik, **F)** impala, **G)** kudu, **H)** giraffe, **I)** Grevy’s zebra, **J)** plains zebra, **K)** spotted hyena, and **L)** leopard. The models assume that *Opuntia* does not indirectly affect occupancy through altering the composition of the native plant community; for model structure, see Figure 2. Shaded areas represent (from outside) 95%, 89%, 80%, 70%, 60%, and 50% compatibility intervals for the January-April (light green) and October-November (purple) seasons. Black lines indicate the posterior median marginal effect for the January-April (solid) and October-November (dashed) seasons.

For the non-frugivorous herbivores, we observed a variety of effects of *Opuntia* on occupancy. While buffalo occupancy did not exhibit a clear relationship with site-level *Opuntia* in either season (Fig. 3D), their occupancy was positively related to grid square *Opuntia* in January-April, and negatively related in October-November (Fig. 4D). For both dik-dik and impala, the posterior median effect of *Opuntia* was positive at both spatial scales, but there was considerable uncertainty (Figs. 3E,F, 4E,F). By contrast, the estimated effects for kudu were positive for site-level *Opuntia* in January-April and negative for grid square *Opuntia* in October-November; we did not observe clear effects for kudu in the other two cases (Figs. 3G, 4G). For reticulated giraffe, the posterior median effect of site-level *Opuntia* was negative in January-April, although the effect was highly uncertain (Fig. 3H). We also observed a slight positive effect of grid square *Opuntia* in the same season (Fig. 4H). Grevy’s zebra occupancy exhibited a weak positive relationship with site-level *Opuntia*, and a stronger negative relationship with grid square *Opuntia*, in January-April (Figs. 3I, 4I). We did not observe any clear effects for October-November. By contrast, the effect of *Opuntia* on plains zebra occupancy was negative at both spatial scales in January-April, but positive at both scales in October-November (Figs. 3J, 4J).

**Figure 4.**
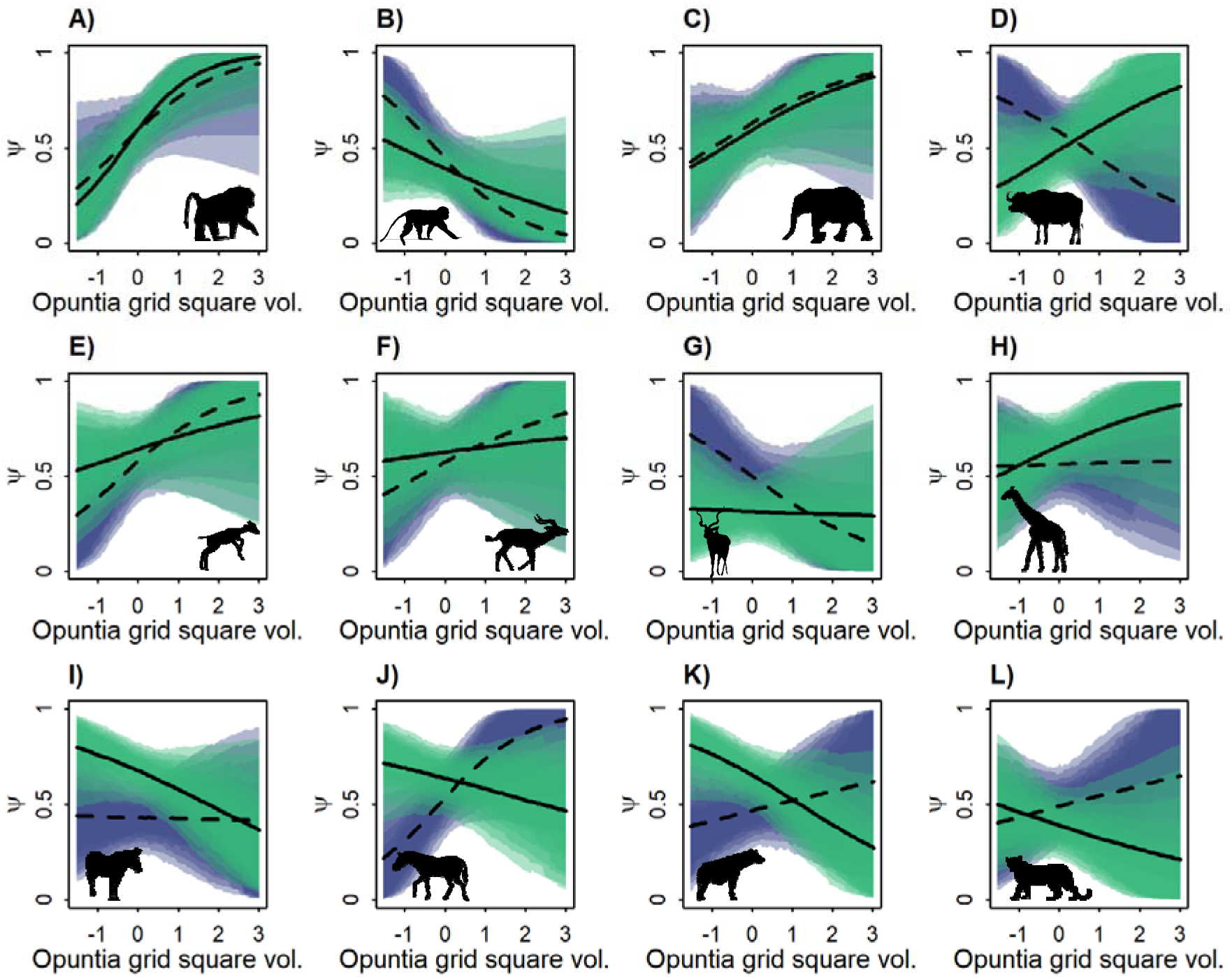
Marginal total effects of grid square-level *Opuntia* volume (standardised) on occupancy probability (ψ) for: **A)** olive baboon, **B)** vervet monkey, **C)** elephant, **D)** buffalo, **E)** dik-dik, **F)** impala, **G)** kudu, **H)** giraffe, **I)** Grevy’s zebra, **J)** plains zebra, **K)** spotted hyena, and **L)** leopard. The models assume that *Opuntia* does not indirectly affect occupancy through altering the composition of the native plant community; for model structure, see Figure 2. Shaded areas represent (from outside) 95%, 89%, 80%, 70%, 60%, and 50% compatibility intervals for the January-April (light green) and October-November (purple) seasons. Black lines indicate the posterior median marginal effect for the January-April (solid) and October-November (dashed) seasons.

Spotted hyena and leopard occupancies were negatively related to *Opuntia* at both spatial scales in January-April (Figs. 3K,L, 4K,L). However, the effects in October-November were positive (albeit uncertain) for both species in the grid square model, and for leopard in the fine-scale model. The effect for hyena in the site-level model was weakly negative.

In all cases, we observed qualitatively similar results for models which included (Figs. 3, 4) and omitted (Figs. S1, S2) measures of the native plant community.

### Total Number of Detections

In our models for the total number of daily detections, the estimated effects of *Opuntia* were generally small and uncertain (Figs. 5, 6). Nevertheless, we did observe clear effects for some species. Olive baboon and elephant detections increased with *Opuntia* at both scales (Figs. 5A,C, 6A,C), except for elephants and site-level *Opuntia* in October-November. Dik-dik and impala exhibited negative effects of site-level *Opuntia* in January-April, but not October-November (Figs. 5E,F). For dik-dik, the posterior median effect in the latter season was positive, while for impala the effect was weakly negative but highly uncertain. The effects of grid square *Opuntia* on these species were less certain, but the posterior median effects were positive for dik-dik (Fig. 6E) and negative for impala (Fig. 6F). Reticulated giraffe detections showed a weak negative relationship with site-level *Opuntia* in January-April (Fig. 5G). Grevy’s zebra detections were also negatively related to *Opuntia* at both scales, but only in October-November (Figs. 5I, 6I). Finally, we observed negative effects at both scales for spotted hyena in January-April.

**Figure 5.**
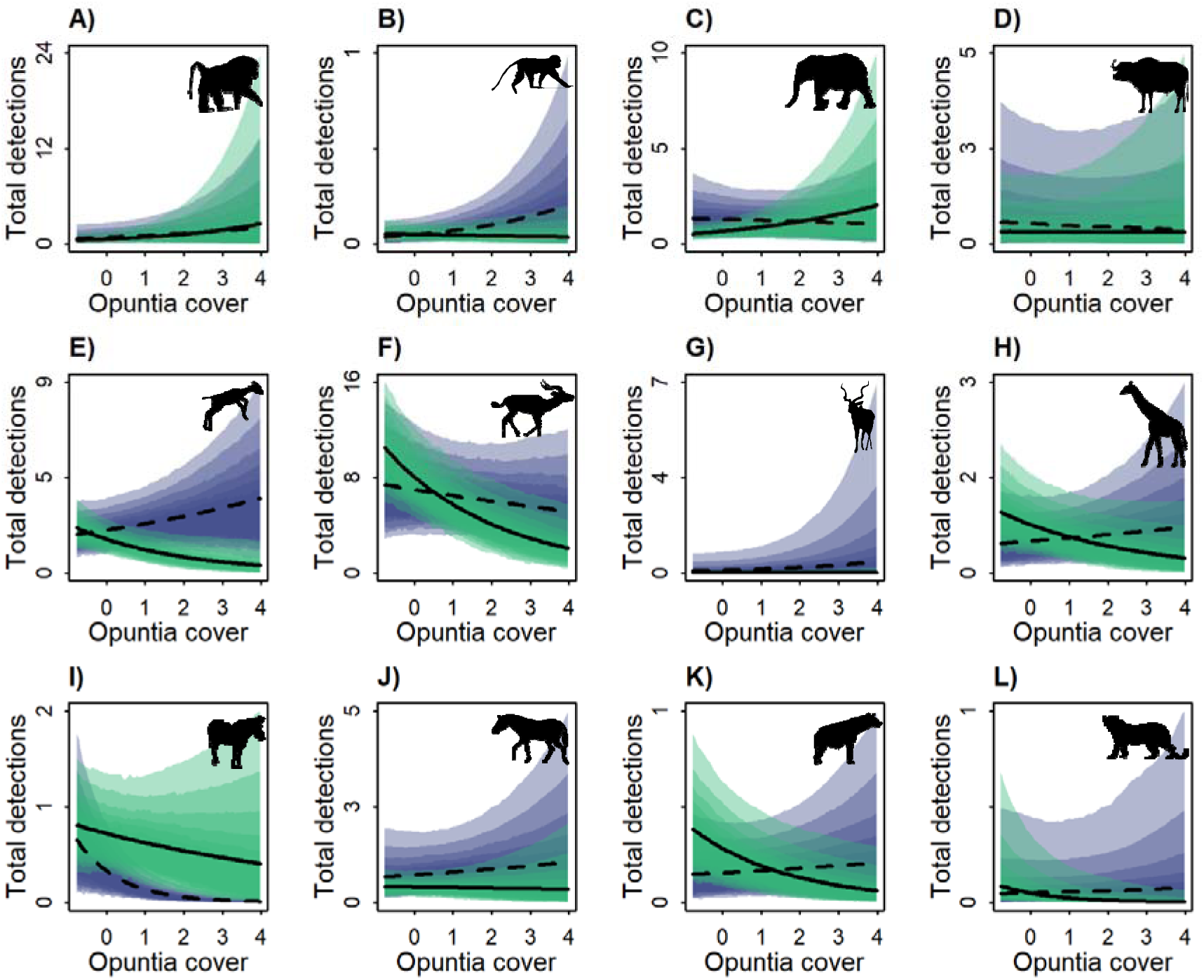
Marginal total effect of site-level *Opuntia* percentage cover (standardised) on the total number of detections per day for: **A)** olive baboon, **B)** vervet monkey, **C)** elephant, **D)** buffalo, **E)** dik-dik, **F)** impala, **G)** kudu, **H)** giraffe, **I)** Grevy’s zebra, **J)** plains zebra, **K)** spotted hyena, and **L)** leopard. The models assume that *Opuntia* does not indirectly affect occupancy through altering the composition of the native plant community; for model structure, see Figure 2. Shaded areas represent (from outside) 95%, 89%, 80%, 70%, 60%, and 50% compatibility intervals for the January-April (light green) and October-November (purple) seasons. Black lines indicate the posterior median marginal effect for the January-April (solid) and October-November (dashed) seasons.

**Figure 6.**
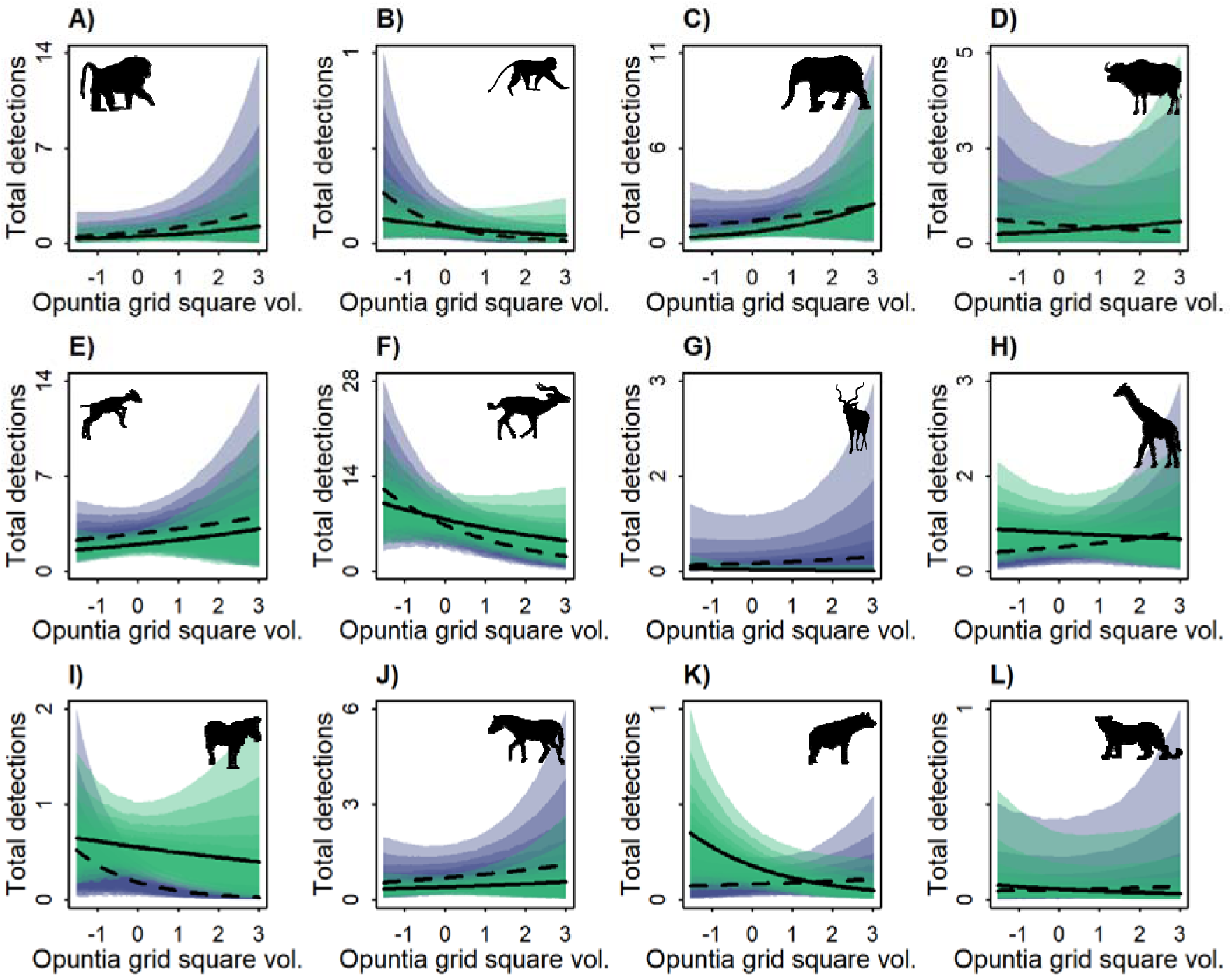
Marginal total effect of grid square-level *Opuntia* volume (standardised) on the total number of detections per day for: **A)** olive baboon, **B)** vervet monkey, **C)** elephant, **D)** buffalo, **E)** dik-dik, **F)** impala, **G)** kudu, **H)** giraffe, **I)** Grevy’s zebra, **J)** plains zebra, **K)** spotted hyena, and **L)** leopard. The models assume that *Opuntia* does not indirectly affect occupancy through altering the composition of the native plant community; for model structure, see Figure 2. Shaded areas represent (from outside) 95%, 89%, 80%, 70%, 60%, and 50% compatibility intervals for the January-April (light green) and October-November (purple) seasons. Black lines indicate the posterior median marginal effect for the January-April (solid) and October-November (dashed) seasons.

As observed for our occupancy models, models which included measures of the native plant community (Figs. 5, 6) and those which omitted them (Figs. S3, S4) generally gave similar results. However, for dik-dik, the effect of site-level *Opuntia* in January-April switched from negative (Fig. 5E) to positive (Fig. S4E).

### Diurnal and Nocturnal Detections

The timing of elephant activity was strongly influenced by site-level *Opuntia*, with a higher proportion of nocturnal detections occurring at high-*Opuntia* sites in both seasons (Fig. S5C). We also observed positive relationships between elephant nocturnal detections and grid square *Opuntia* in October-November (Fig. S6C). During January-April the proportion of night-time buffalo detections was negatively related to site-level *Opuntia* when lunar illumination was high, but positively related to grid square *Opuntia* regardless of lunar illumination (Fig. S6D). Dik-dik were detected less often at night as both site-level and grid square level *Opuntia* increased in October-November, and these effects were stronger under a full moon (Figs. S5E, S6E). Impala were observed more often at night as site-level *Opuntia* increased during October-November, but only under high lunar illumination (Fig. S5F). Conversely, the proportion of night-time detections decreased with grid square *Opuntia*, again only when the moon was full. For kudu, we observed a positive effect of grid square *Opuntia* which was strongest under high lunar illumination (Fig S6G). For giraffe, the proportion of night-time detections increased with grid square *Opuntia* in January-April under high lunar illumination, and decreased in October-November (Fig S6H). For Grevy’s zebra, the effects of grid-level *Opuntia* were negative in January-April under high lunar illumination and in October-November under low lunar illumination, but positive in October-November when lunar illumination was high (Fig. S6I).

Again, we observed qualitatively similar results regardless of whether we included measures of the native plant community (Figs. S7, S8).

## Discussion

We assessed the impacts of invasive *Opuntia* cacti on the occupancy and activity of twelve mammal species. We found that *Opuntia* exerted strong and consistent positive effects on the occupancy, and to a lesser extent the daily number of detections, of two key *Opuntia* fruit consumers: olive baboons and elephants. We also observed seasonally varying effects on the occupancy of Grevy’s zebra and plains zebra, two of the region’s key grazer species. For the other five herbivore species in our sample, we observed a variety of responses to increasing *Opuntia*. We also observed seasonally varying effects for two predator species: spotted hyena and leopard. Across all species, our results were similar regardless of whether we included measures of the native plant community in our models, suggesting that *Opuntia*’s effects are not mediated by changes to native vegetation cover.

The positive effects of *Opuntia* on olive baboon and elephant occupancy were strong and consistent across seasons and scales of *Opuntia* covariate. Furthermore, we generally observed positive effects of *Opuntia* on the daily number of detections for both species. These results are unsurprising; both olive baboons and elephants regularly consume *Opuntia* fruit, and are thought to be key dispersal agents for *Opuntia* (Githae 2019; Strum *et al*. 2015; Witt 2017). The findings for elephants are also consistent with local communities’ observation that elephants are attracted to *Opuntia* fruit, resulting in elephants encroaching with increased frequency on grazing areas near *Opuntia* stands (Shackleton *et al*. 2017).

Interestingly, we observed a shift towards nocturnal activity for elephants as site-level *Opuntia* increased. This suggests that *Opuntia* invasion around agricultural areas could exacerbate crop foraging, which occurs at night (Graham *et al*. 2010). For vervet monkeys, which are also frugivorous, occupancy and activity did not generally increase with increasing *Opuntia*; only the effects of site-level *Opuntia* on occupancy in January-April and total detections in October-November were clearly positive. This result may be attributable to competition with and the risk of predation from baboons (Willems & Hill 2009).

The positive effects of *Opuntia* on the occurrence of elephants and olive baboons may generate a positive feedback loop which facilitates the spread of the invasion: these frugivores are drawn to invaded areas, where they consume and subsequently disperse *Opuntia seeds*. This further increases the extent of the invaded area, in turn attracting more frugivores. However, understanding how elephant and olive baboon habitat use feeds back to influence the *Opuntia* invasion requires further information on how far seeds are carried before they are deposited and the proportion of deposition which occurs in currently invaded areas versus adjacent uninvaded habitat.

A second key finding was the seasonally-varying effect of *Opuntia* on the occupancy of Grevy’s zebra and plains zebra – two grazers which are not known to consume *Opuntia* fruit. In the predominantly dry January-April season, the occupancies of both species declined as the quantity of *Opuntia* in the grid square increased, while in the wetter October-November season the effect of *Opuntia* was neutral for Grevy’s zebra and positive for plains zebra. These results may be explained by changes in forage availability; in wet conditions, when grass biomass is high (Boutton *et al*. 1988), there is ample forage for zebra to occupy heavily invaded areas. However, as grass biomass declines in dry conditions (Boutton *et al*. 1988), the areas between *Opuntia* stands no longer contain enough forage for zebra to persist, and they must instead concentrate on less invaded areas. This explanation is compatible with the fact that our models displayed similar results regardless of whether we included native vegetation (including grass) cover because grasses commonly grow among *Opuntia* stands, and hence contribute to percentage cover despite potentially being inaccessible to zebra. We also observed that the average number of Grevy’s zebra detections declined with increasing *Opuntia* in both seasons; this may suggest that even though high-*Opuntia* areas remain occupied, they are used less frequently. We did not observe similar effects for plains zebra; this difference could be explained by a number of factors, including subtle dietary differences between the two species (Kartzinel *et al*. 2015).

The reduction in Grevy’s zebra occupancy and detections in high-*Opuntia* areas under dry conditions is especially significant for conservation, because it implies that the ongoing *Opuntia* invasion may erode the zebras’ ability to withstand the frequent – and often severe – droughts which the region experiences (Ndiritu 2021). As Laikipia County hosts approximately half of the world’s Grevy’s zebra population (Rubenstein *et al*. 2016), the synergistic effects of *Opuntia* invasion and drought may pose a threat to the species. Consequently, further investigation into the effects of *Opuntia* on Grevy’s zebra presents a key avenue for future research.

We also examined the effects of *Opuntia* on five non-equid species of non-frugivorous herbivore: buffalo, dik-dik, impala, greater kudu, and reticulated giraffe. We observed a range of responses to increasing *Opuntia* among these species, which is perhaps unsurprising because the species differ substantially in diet (Kartzinel *et al*. 2015) and in their response to predation risk (Thaker *et al*. 2011). Dik-dik occupancy and daily detections were positively related to *Opuntia*, while impala daily detections decreased in invaded areas. These findings align with the species’ preferences for open *versus* closed habitats; dik-dik prefer closed habitats because they perceive open areas as risky, while impala tend to avoid closed habitats as they are less able to detect and escape from predators (Epperly *et al*. 2021; Ford *et al*. 2014; Otieno *et al*. 2019). Although the positive effect of *Opuntia* on impala occupancy initially appears inconsistent with this explanation, the discrepancy could be explained by invaded areas being occupied by poor-condition individuals (McNamara & Houston 1987; Winnie & Creel 2007) or bachelor herds. For the larger-bodied herbivores, particularly giraffe and buffalo, the habitat use of adult individuals is less likely to be structured by predation risk (Thaker *et al*. 2011). However, we note that buffalo occupancy is inversely related to grid square *Opuntia* in the wetter October-November season; one possible explanation is that buffalo avoid invaded areas close to the birthing season, which coincides with the wet season (Ryan *et al*. 2007), due to the risk of calves being predated.

The heterogeneous effects we observed across herbivore species suggest that *Opuntia* invasion can substantially alter the composition of the local herbivore community. Long-term herbivore exclusion experiments – conducted in the same region as our study (Goheen *et al*. 2013; Young *et al*. 1997) – illustrate that these compositional changes could result in powerful indirect effects on a number of ecosystem properties. For instance, mesoherbivores – including impala, kudu, buffalo and zebra – strongly reduce total understory density (Goheen *et al*. 2013), while megaherbivores – especially elephants – regulate shrub and tree dynamics, reducing the cover of taller shrubs (Augustine & Mcnaughton 2004; Goheen *et al*. 2013) and influencing species composition (Augustine & Mcnaughton 2004). Smaller herbivores also play a key role in shrub and tree dynamics; dik-dik suppress the growth rate and biomass of shrubs close to the ground, and reduce the recruitment of shrubs into larger size classes (Augustine & Mcnaughton 2004), while impala exert effects which manifest as shifts in the relative abundance of thorny *versus* less-thorny tree species (Ford *et al*. 2014).

Exclusion experiments also illustrate that herbivores’ effects extend beyond the plant community. For example, herbivores affect the nitrogen cycle, and these effects appear to differ between browsers and grazers (Coetsee *et al*. 2023). Finally, the herbivore exclusion experiments suggest that the effects of *Opuntia* on mammalian herbivores may feed back to influence the future dynamics of the *Opuntia* invasion; herbivore-accessible plots have significantly fewer *Opuntia* plants than plots which exclude herbivores, which may indicate that herbivores inhibit *Opuntia* establishment via herbivory or trampling (Wells *et al*. 2023). Further investigation of *Opuntia*’s herbivore-mediated effects on the native plant community, nutrient cycles, and future invasion dynamics represent important avenues for future research.

Finally, we explored the effects of *Opuntia* on two predators. For hyena, the effects on occupancy and activity were generally negative in January-April but neutral or positive in October-November; the only exception was that the effect of site-level *Opuntia* on occupancy was negative regardless of season. By contrast, the effects on leopard occupancy were positive in January-April and negative in October-November at both spatial scales of *Opuntia* covariate; we did not observe any clear effects of *Opuntia* on the number of leopard detections. These results could be partly driven by the hunting strategies of the two predator species; hyena are cursorial predators (Périquet *et al*. 2015) while leopards are ambush hunters (Kruuk & Turner 1967), so it is likely that the former will be impeded by high levels of site-level *Opuntia* while the latter may benefit. The seasonal differences which we observed could be explained by changes to prey availability; at sites with high *Opuntia* cover, the number of dik-dik and impala detections was higher in October-November than in January-April. However, we note that the fission-fusion dynamics of spotted hyena groups, which are also influenced by prey availability (Smith *et al*. 2008), may manifest as changes in occupancy or the number of detections. Furthermore, as both species are territorial (Boydston *et al*. 2001; Fattebert *et al*. 2016), changes to territory size or location may also influence the results.

We have shown that the *Opuntia* invasion in Laikipia County, Kenya, affects mammal occupancy and activity; the strength and direction of these effects varies among species and between seasons, and depends on the scale at which *Opuntia* was considered. These results have important implications for the conservation of endangered mammal species, native plant community composition and functioning, and the future spread of *Opuntia* in the region.

More broadly, our findings underscore the role of animal behaviour – and the behaviour of large mammals specifically – in mediating the ecological impacts of biological invasions. As invasive species continue to spread worldwide, obtaining a stronger understanding of these behavioural impacts will become increasingly important.

## Acknowledgements

We thank Mpala Research Centre and Loisaba Conservancy for facilitating and supporting the research. Particular thanks is extended to Peter Leidura, Ibrahim Adan, Boniface Lowoi, and Cate Lonyangaita for their assistance in data collection, and to Tom Silvester, Dino Martins, Fardosa Hassan, Cosmas Nzomo, Beatrice Wanjohi, David Hewett, Tony Maina, Thomas Koitimet, Julius Nakolonyo, Hannah Campbell, and Susan Lentaam for their logistical support and advice. The authors would also like to thank NACOSTI and Dr. Vincent Obanda from WRTI for permitting the research.

## Statement of Authorship

P.S.S and W.D. designed the data collection protocol. P.S.S collected and analysed the data, and wrote the first manuscript draft. All authors contributed substantially to revisions of the framework and manuscript.

## Data accessibility statement

All data and code required to reproduce our analyses are available on Zenodo (code: https://doi.org/10.5281/zenodo.14388983 data: https://doi.org/10.5281/zenodo.14389490). Our camera trap images are available at: https://www.zooniverse.org/projects/peter-dot-stewart/prickly-pear-project-kenya

## Notes

https://doi.org/10.5281/zenodo.14388983

https://doi.org/10.5281/zenodo.14389490

### Competing Interest Statement

The authors have declared no competing interest.

