## Supplementary figures and tables for "Contrasting impacts of invasive *Opuntia* cacti on mammal habitat use"

**Supplementary Material**

**Appendix S1 – Supplementary Figures and Tables**

**Table S1.** Classification accuracy for volunteer classifications, based on 26952 images which were classified by both an expert and volunteers. Results are shown for both consensus classifications (see main text) as well as the raw non-consensus volunteer classifications. Sensitivity and specificity are defined as Pr(classified present | truly present) and Pr(classified absent | truly absent) respectively. Species marked with an asterisk (*) were not present in the expert-classified set of images, and therefore do not have sensitivity values as they were never truly present.

|  | **Raw** | | **Consensus** | |
| --- | --- | --- | --- | --- |
|  | **Sensitivity** | **Specificity** | **Sensitivity** | **Specificity** |
| **Focal species** | |  |  |  |
| Buffalo *(Syncerus caffer)* | 0.785 | 0.999 | 1.000 | 1.000 |
| Dik-dik *(Madoqua spp.)* | 0.848 | 0.985 | 0.993 | 1.000 |
| Elephant *(Loxodonta africana)* | 0.948 | 0.999 | 0.995 | 1.000 |
| Greater kudu *(Tragelaphus strepsiceros)* | 0.638 | 0.999 | 1.000 | 1.000 |
| Grevy’s zebra *(Equus grevyi)* | 0.762 | 0.995 | 0.973 | 1.000 |
| Impala *(Aepyceros melampus)* | 0.834 | 0.991 | 0.996 | 0.999 |
| Leopard *(Panthera pardus)* | 0.650 | 1.000 | 1.000 | 1.000 |
| Olive baboon *(Papio anubis)* | 0.899 | 0.999 | 0.997 | 1.000 |
| Plains zebra *(Equus quagga)* | 0.913 | 0.991 | 0.990 | 0.998 |
| Reticulated giraffe *(Giraffa reticulata)* | 0.908 | 1.000 | 0.995 | 1.000 |
| Spotted hyena *(Crocuta crocuta)* | 0.737 | 1.000 | 0.991 | 1.000 |
| Vervet monkey *(Chlorocebus pygerythrus)* | 0.879 | 0.999 | 1.000 | 1.000 |
| **Other mammals** | |  |  |  |
| Aardvark *(Orycteropus afer)** | NA | 0.999 | NA | 1.000 |
| Aardwolf *(Proteles cristata)** | NA | 1.000 | NA | 1.000 |
| African wild dog *(Lycaon pictus)** | 0.636 | 1.000 | 1.000 | 1.000 |
| African wildcat *(Felis lybica)** | NA | 1.000 | NA | 1.000 |
| Bat | 0.500 | 1.000 | 0.941 | 1.000 |
| Black-backed jackal *(Canis mesomelas)* | 0.479 | 0.999 | 1.000 | 1.000 |
|  | 0.872 | 0.999 | 0.993 | 0.999 |
| Bushbuck *(Tragelaphus sylvaticus)* | NA | 1.000 | NA | 1.000 |
| Camel *(Camelus dromedarius)* | NA | 1.000 | NA | 1.000 |
| Caracal *(Caracal caracal)** | 0.083 | 1.000 | 0.500 | 1.000 |
| Cheetah *(Acinonyx jubatus)** | NA | 1.000 | NA | 1.000 |
| Civet *(Civettictis civetta)** | 0.083 | 0.994 | 0.500 | 1.000 |
| Dog (domestic)* | 0.402 | 0.999 | 0.962 | 1.000 |
| Duiker (tribe Cephalophini)* | 0.722 | 1.000 | 1.000 | 1.000 |
| Eland *(Taurotragus oryx)* | NA | 0.995 | NA | 1.000 |
| Genet *(Genetta spp.)* | 0.388 | 0.984 | 0.857 | 1.000 |
| Gerenuk *(Lotocranius walleri)** | 0.762 | 0.999 | 0.990 | 1.000 |
| Grant’s gazelle *(Nanger granti)* | NA | 0.999 | NA | 1.000 |
|  | 0.916 | 1.000 | 1.000 | 1.000 |
| Hare *(Lepus victoriae)* | 0.745 | 1.000 | 1.000 | 1.000 |
| Hartebeest *(Alcelaphus buselaphus)** | NA | 1.000 | NA | 1.000 |
| Hippopotamus *(Hippopotamus amphibius)* | 0.848 | 1.000 | 1.000 | 1.000 |
| Honey badger *(Mellivora capensis)* | 0.692 | 1.000 | 1.000 | 1.000 |
| Hyrax (family Procaviidae)* | 0.822 | 0.996 | 0.927 | 1.000 |
| Lion *(Panthera leo)* | 0.762 | 0.999 | 1.000 | 1.000 |
| Livestock (non-camel) | 0.575 | 1.000 | 1.000 | 1.000 |
| Mongoose (family Herpestidae) | 0.750 | 0.999 | 1.000 | 1.000 |
| Mouse/rat | 0.441 | 1.000 | 1.000 | 1.000 |
| Oryx *(Oryx beisa)* | 0.769 | 0.999 | 0.978 | 1.000 |
| Porcupine *(Hystrix cristata)* | NA | 0.997 | NA | 1.000 |
| Squirrel (tribe Xerini) | 0.563 | 1.000 | 0.933 | 1.000 |
| Steenbok *(Raphicerus campestris)** | NA | 0.998 | NA | 1.000 |
| Striped hyena *(Hyaena hyaena)* | 0.844 | 0.999 | 0.988 | 1.000 |
| Thompson’s gazelle *(Nanger granti)** | 0.548 | 0.999 | 0.989 | 1.000 |
| Warthog *(Phacochoerus spp.)* | 0.520 | 1.000 | 1.000 | 1.000 |
| Waterbuck *(Kobus defassa)* | NA | 0.999 | NA | 1.000 |
| Zorilla *(Ictonyx striatus)* | NA | 1.000 | NA | 1.000 |
| **Birds and other taxa** |  |  |  |  |
| Bird (other) | 0.806 | 0.995 | 0.975 | 0.999 |
| Helmeted guineafowl *(Numida meleagris)* | 0.738 | 0.996 | 0.977 | 1.000 |
| Insect/spider | 0.083 | 0.999 | 0.500 | 1.000 |
| Kori bustard *(Ardeotis kori)* | 0.783 | 0.999 | 1.000 | 1.000 |
| Ostrich *(Struthio camelus)** | NA | 1.000 | NA | 1.000 |
| Reptile/amphibian | 0.815 | 1.000 | 1.000 | 1.000 |
| Secretary bird *(Sagittarius serpentarius)* | 0.848 | 1.000 | 1.000 | 1.000 |
| Vulturine guineafowl *(Acryllium vulturinum)* | 0.901 | 0.995 | 0.997 | 1.000 |
| **Special categories** |  |  |  |  |
| Human | 0.214 | 1.000 | 1.000 | 1.000 |
| Other | 0.267 | 0.990 | 0.667 | 1.000 |
| No animals present | 0.954 | 0.969 | 0.994 | 0.996 |

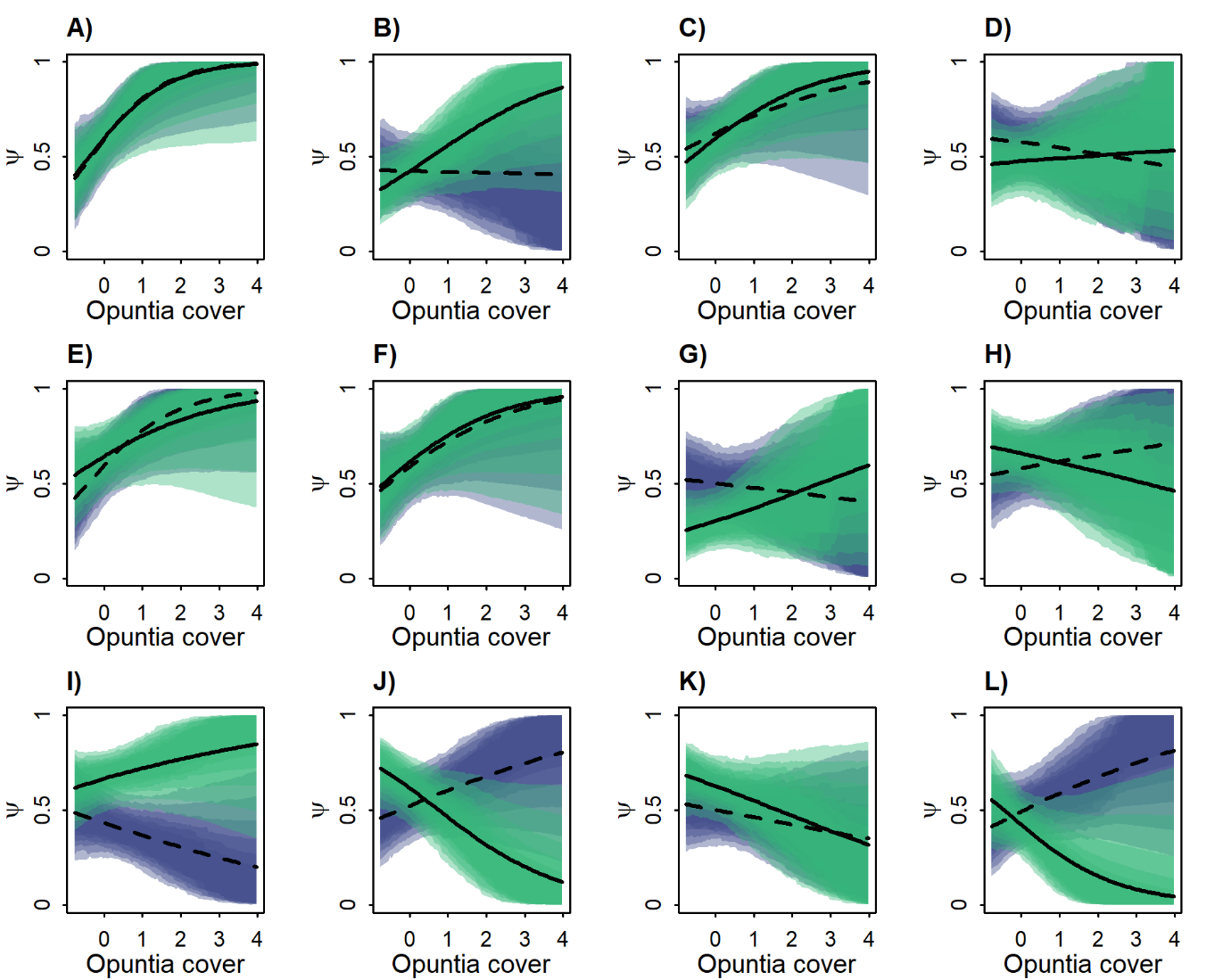

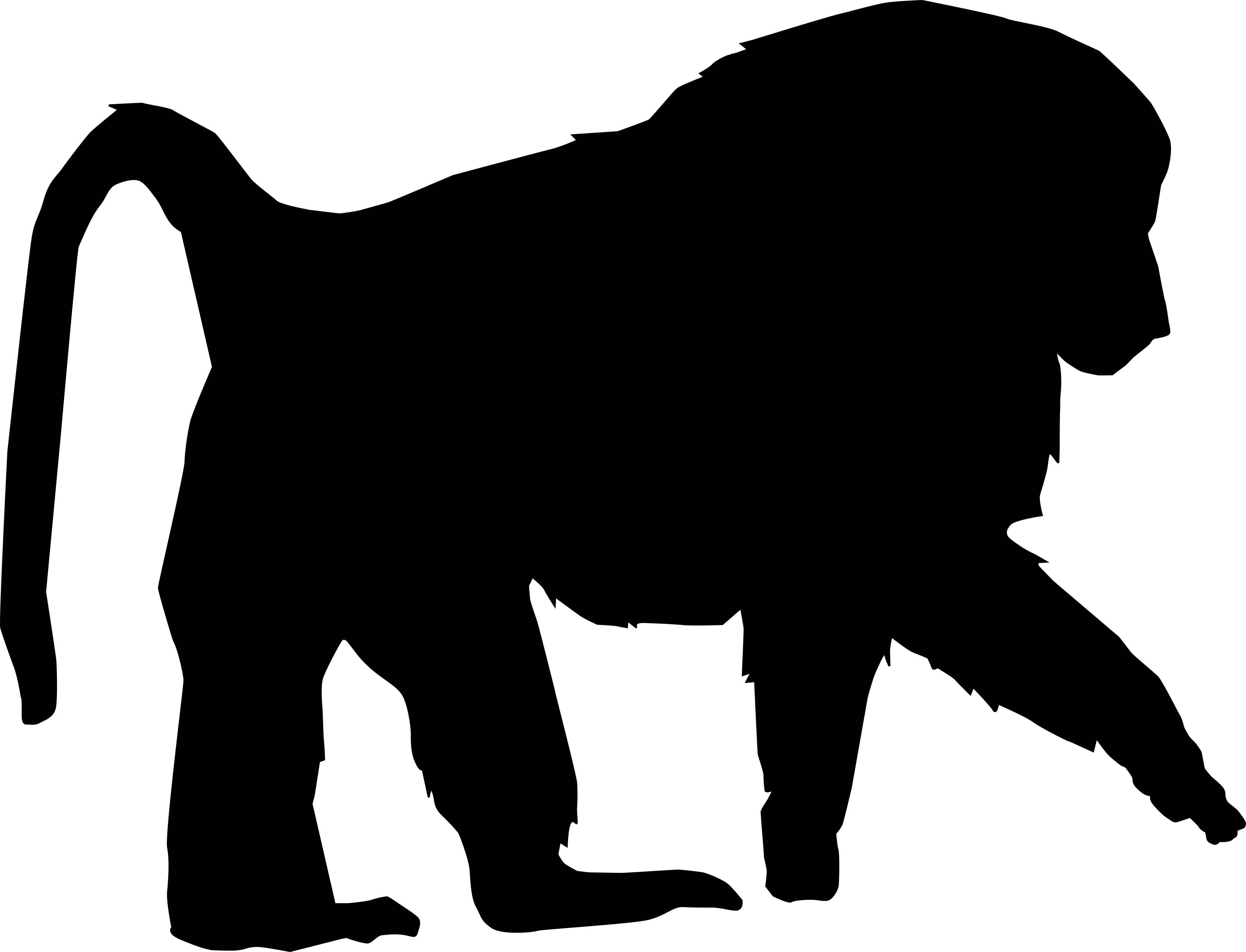

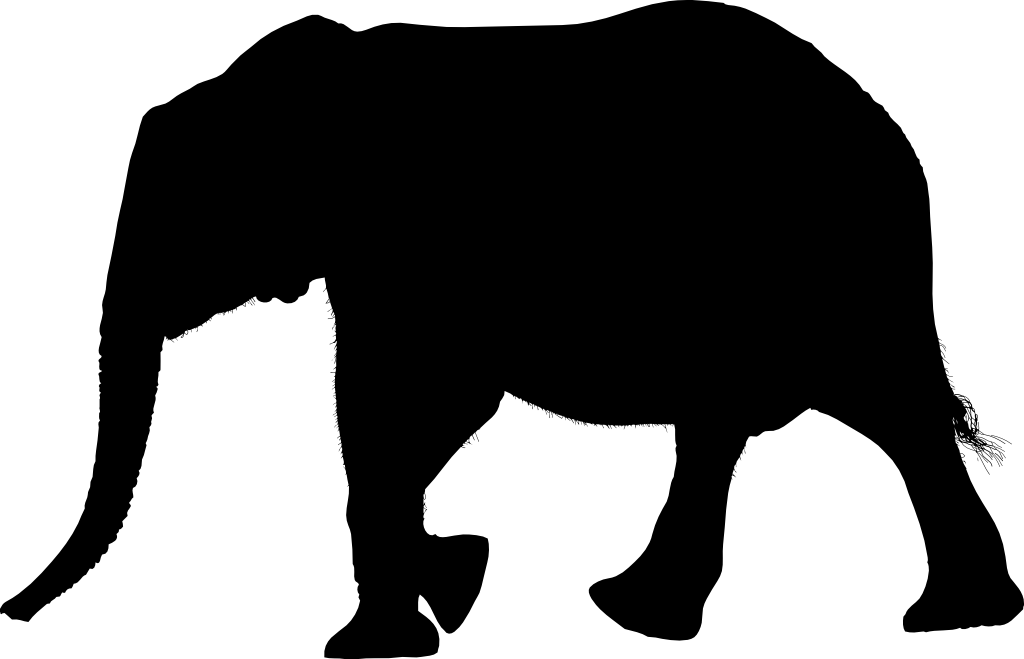

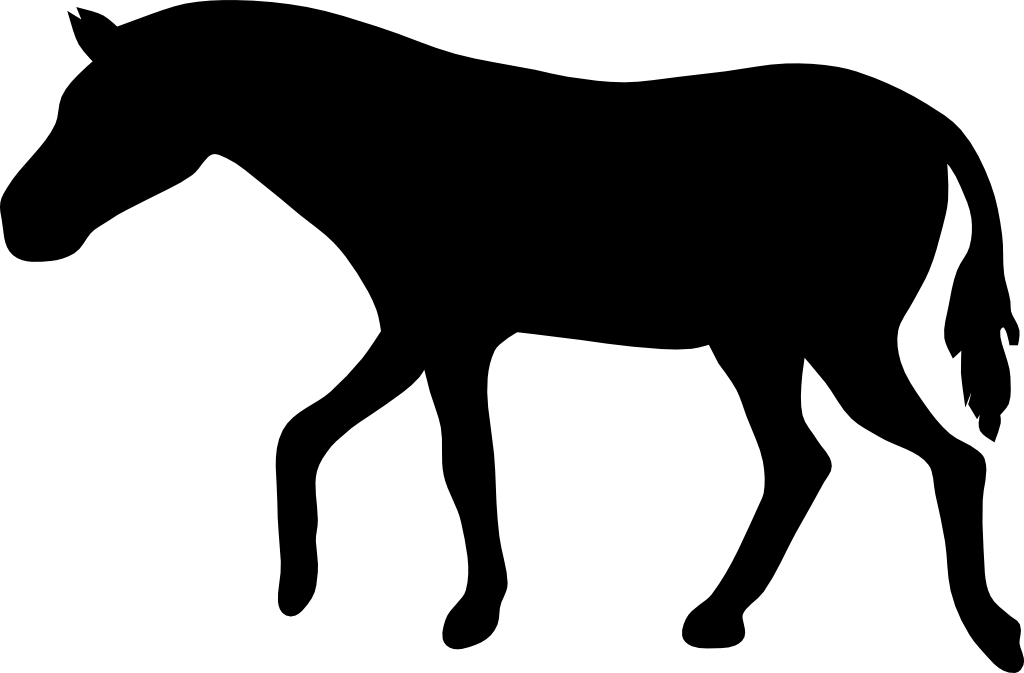

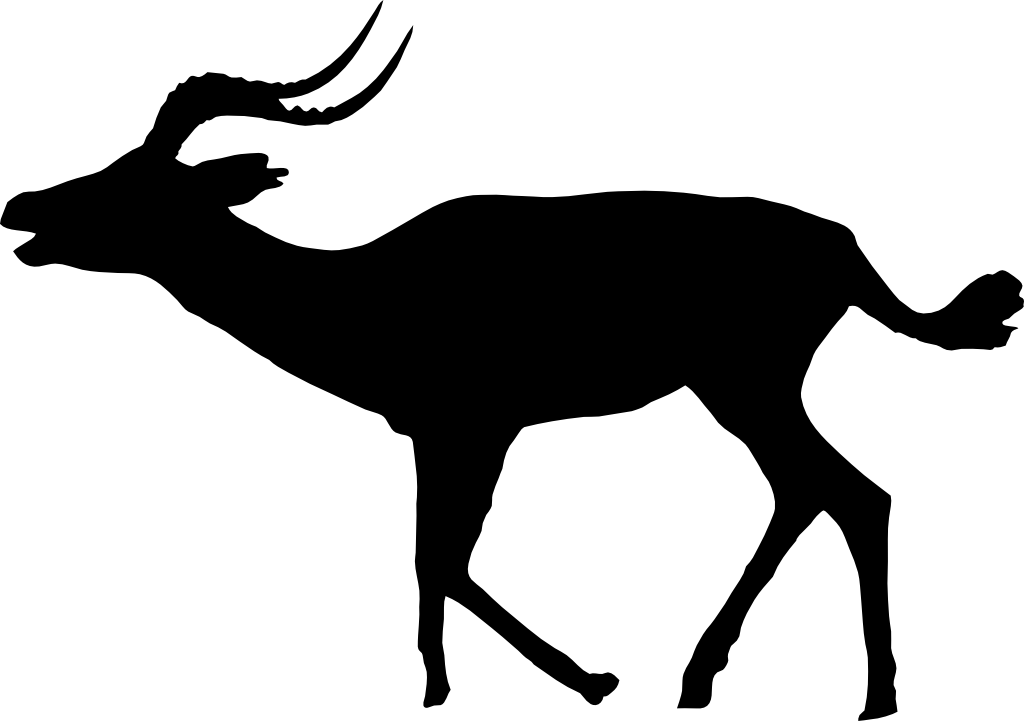

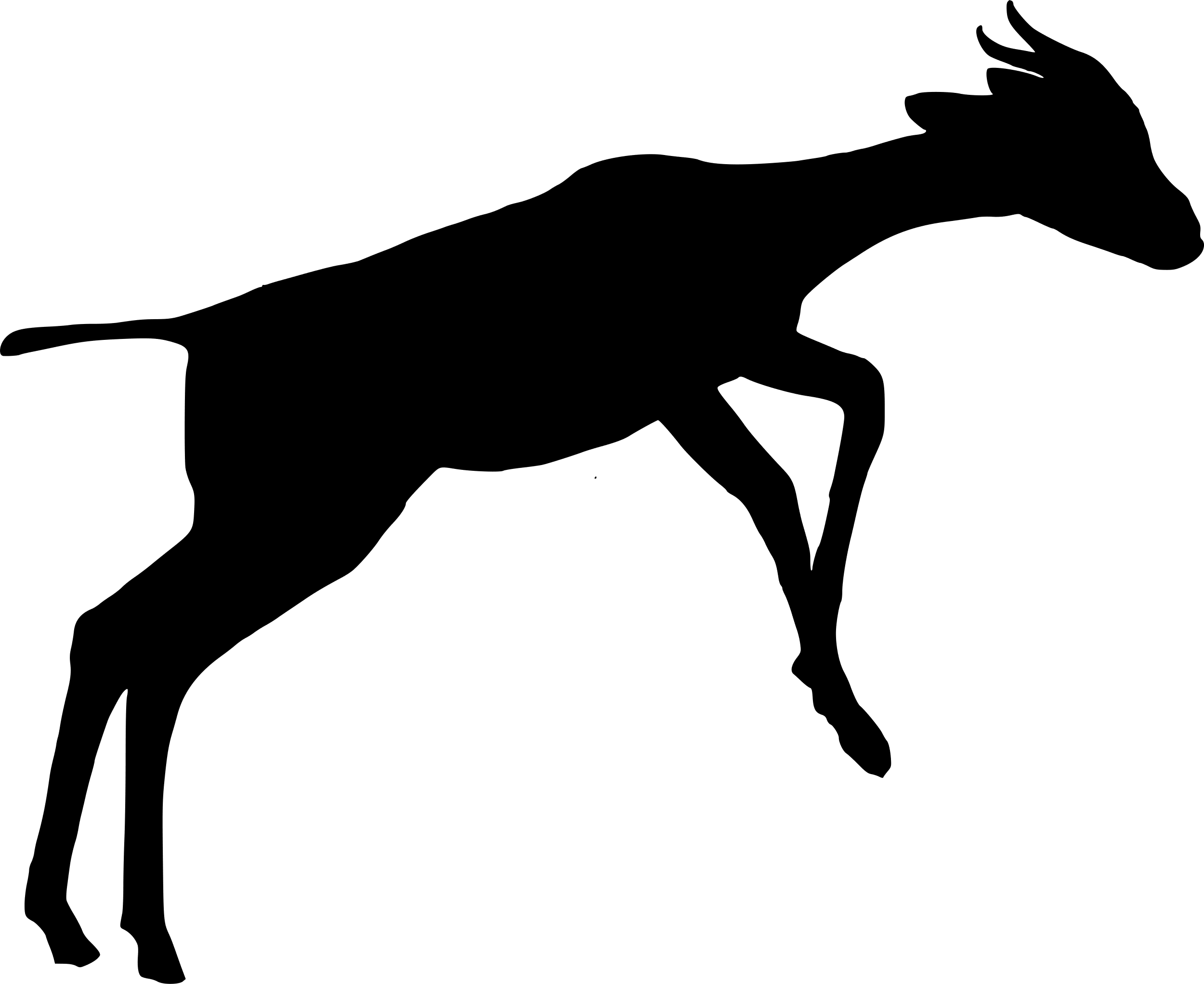

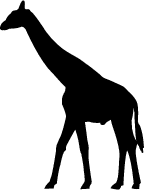

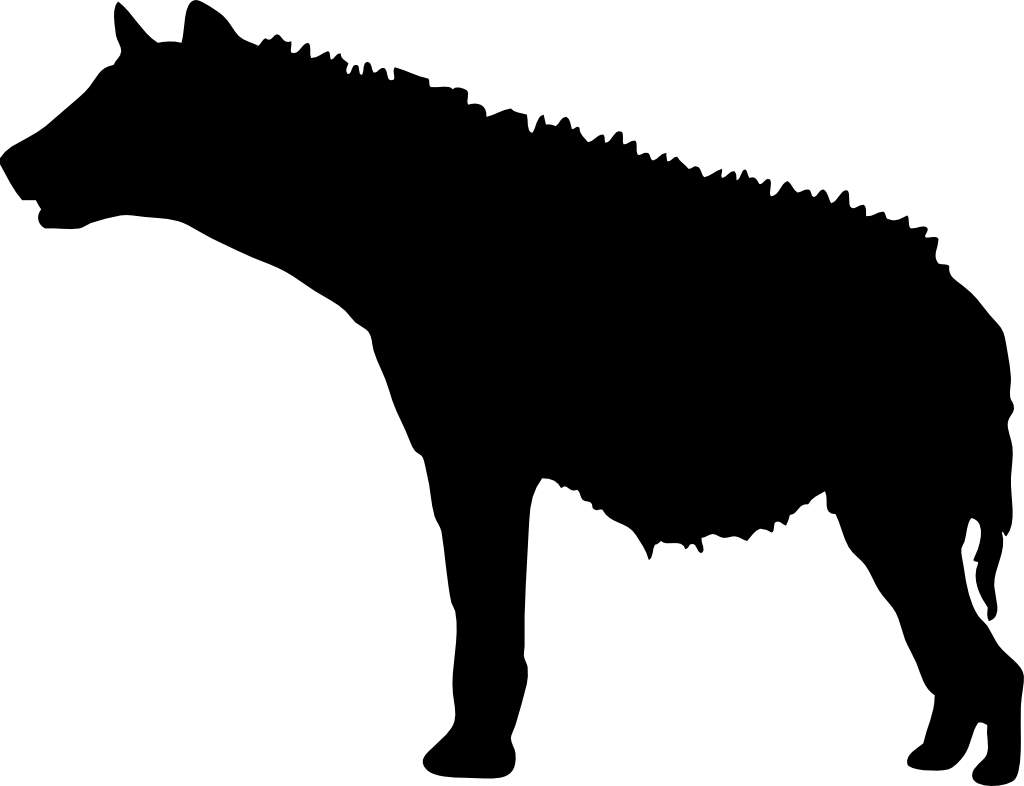

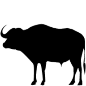

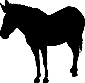

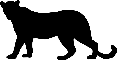

**Figure S1.** Marginal total effect of site-level *Opuntia* percentage cover (standardised) on occupancy probability (ψ) for: **A)** olive baboon, **B)** vervet monkey, **C)** elephant, **D)** buffalo, **E)** dik-dik, **F)** impala, **G)** kudu, **H)** giraffe, **I)** Grevy’s zebra, **J)** plains zebra, **K)** spotted hyena, and **L)** leopard. The models assume that *Opuntia* indirectly affects occupancy through altering the composition of the native plant community; for model structure, see Figure 2 in main text. Shaded areas represent (from outside) 95%, 89%, 80%, 70%, 60%, and 50% compatibility intervals for the January-April (light green) and October-November (purple) seasons. Black lines indicate the posterior median marginal effect for the January-April (solid) and October-November (dashed) seasons.

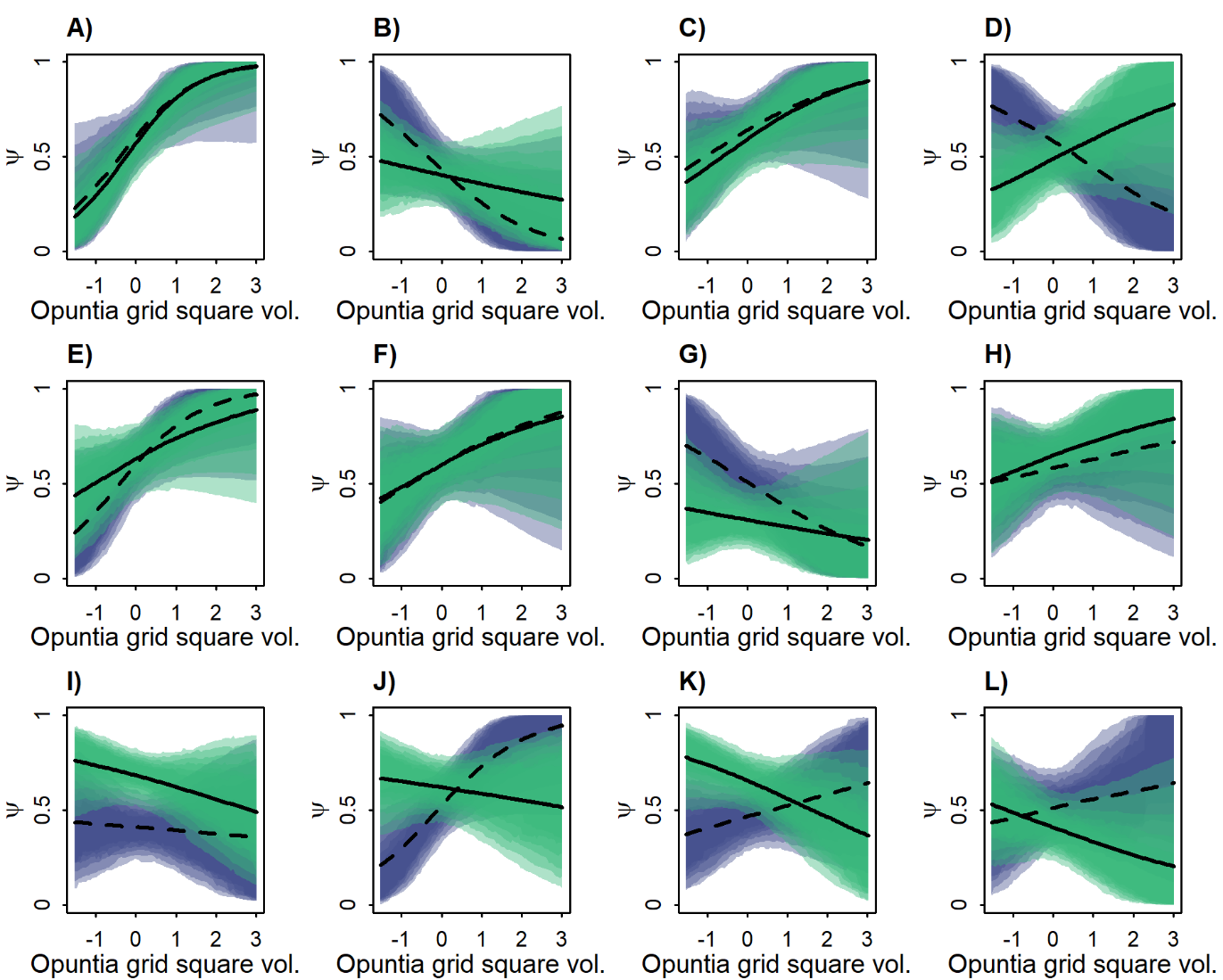

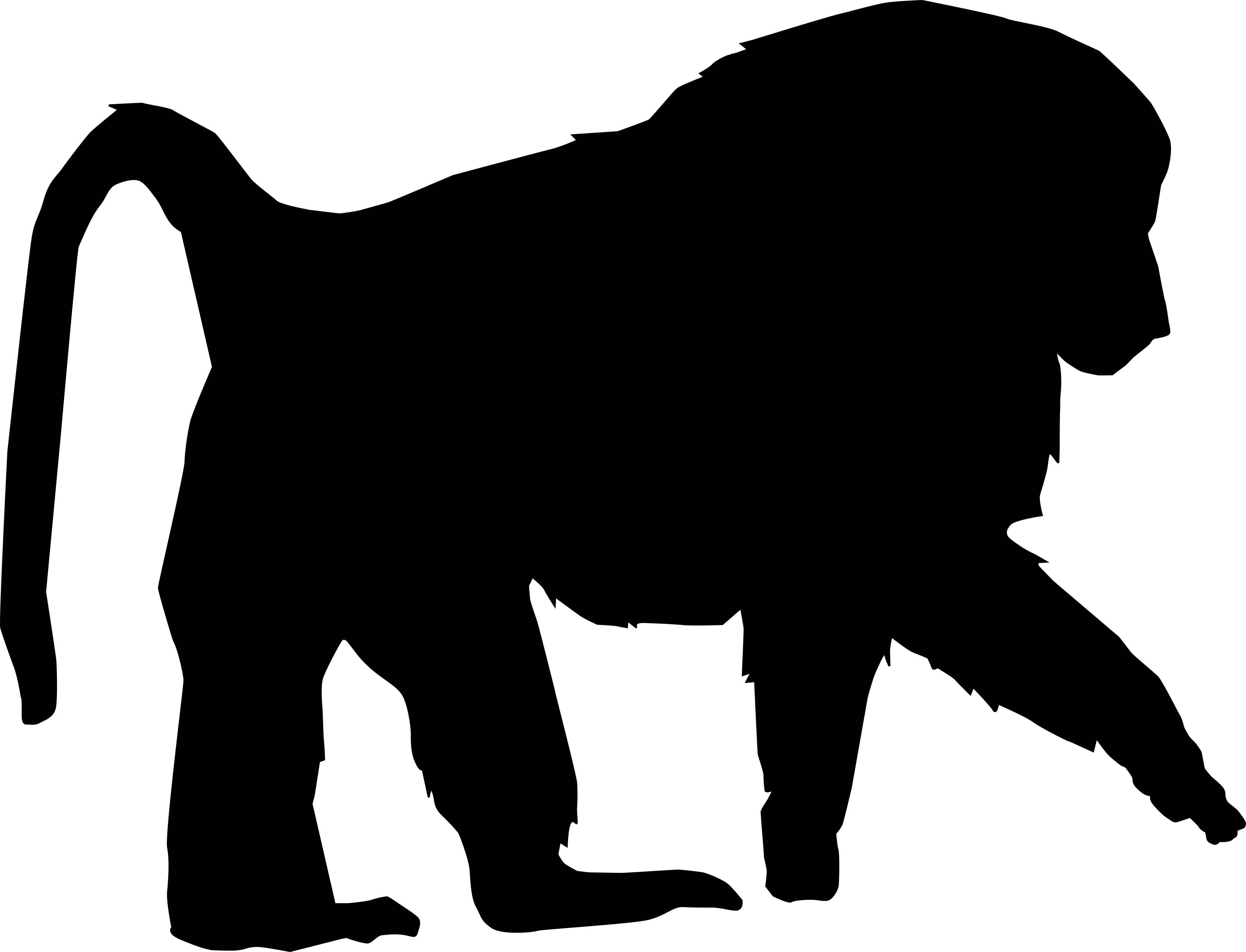

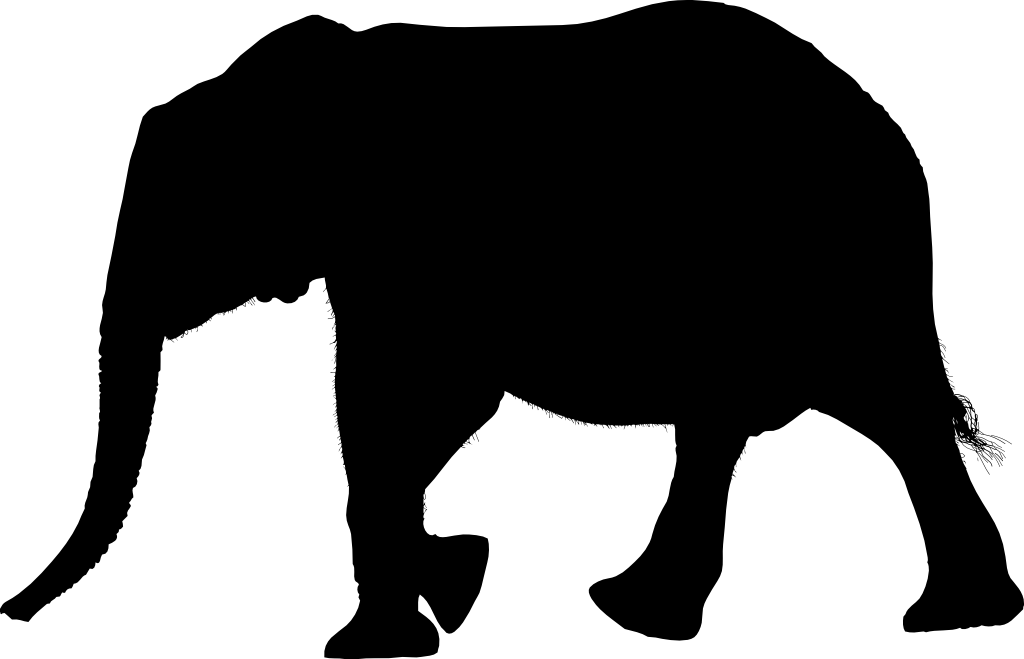

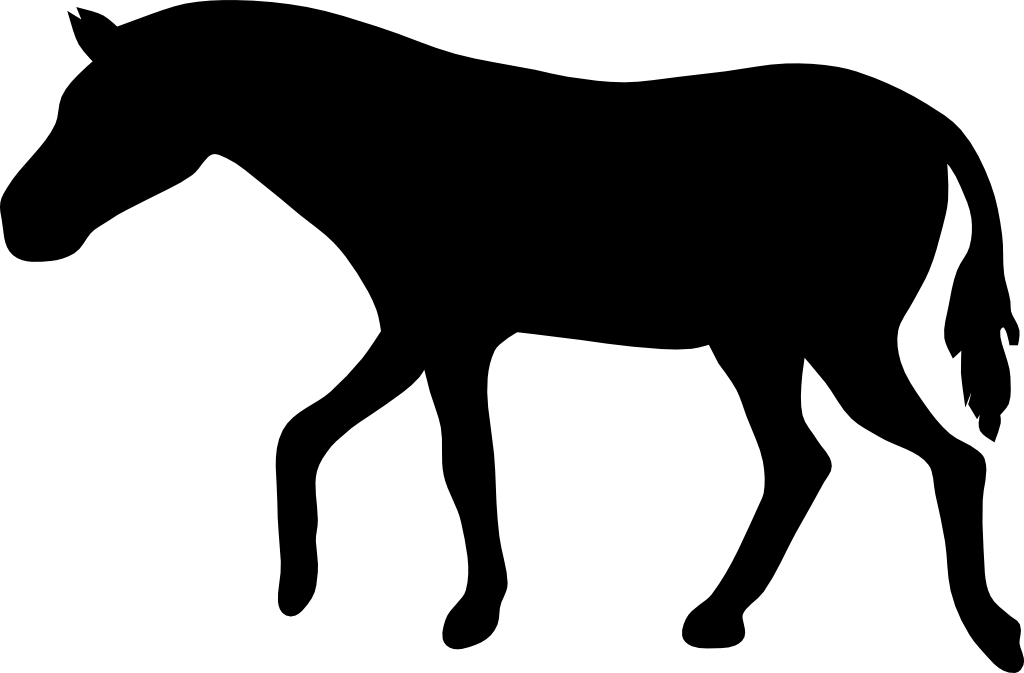

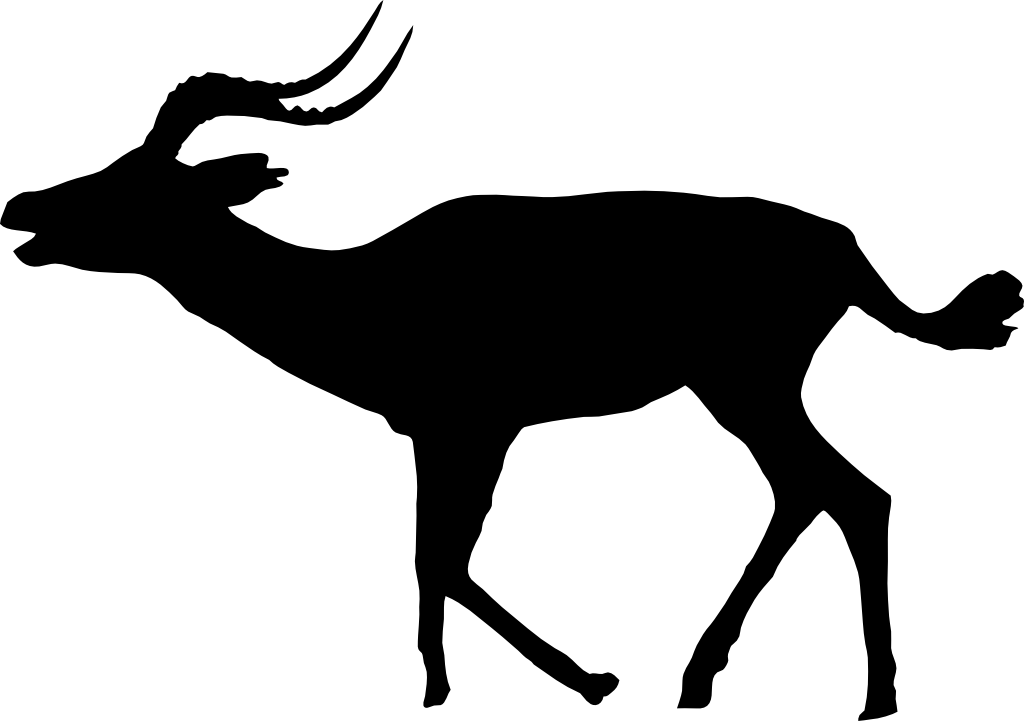

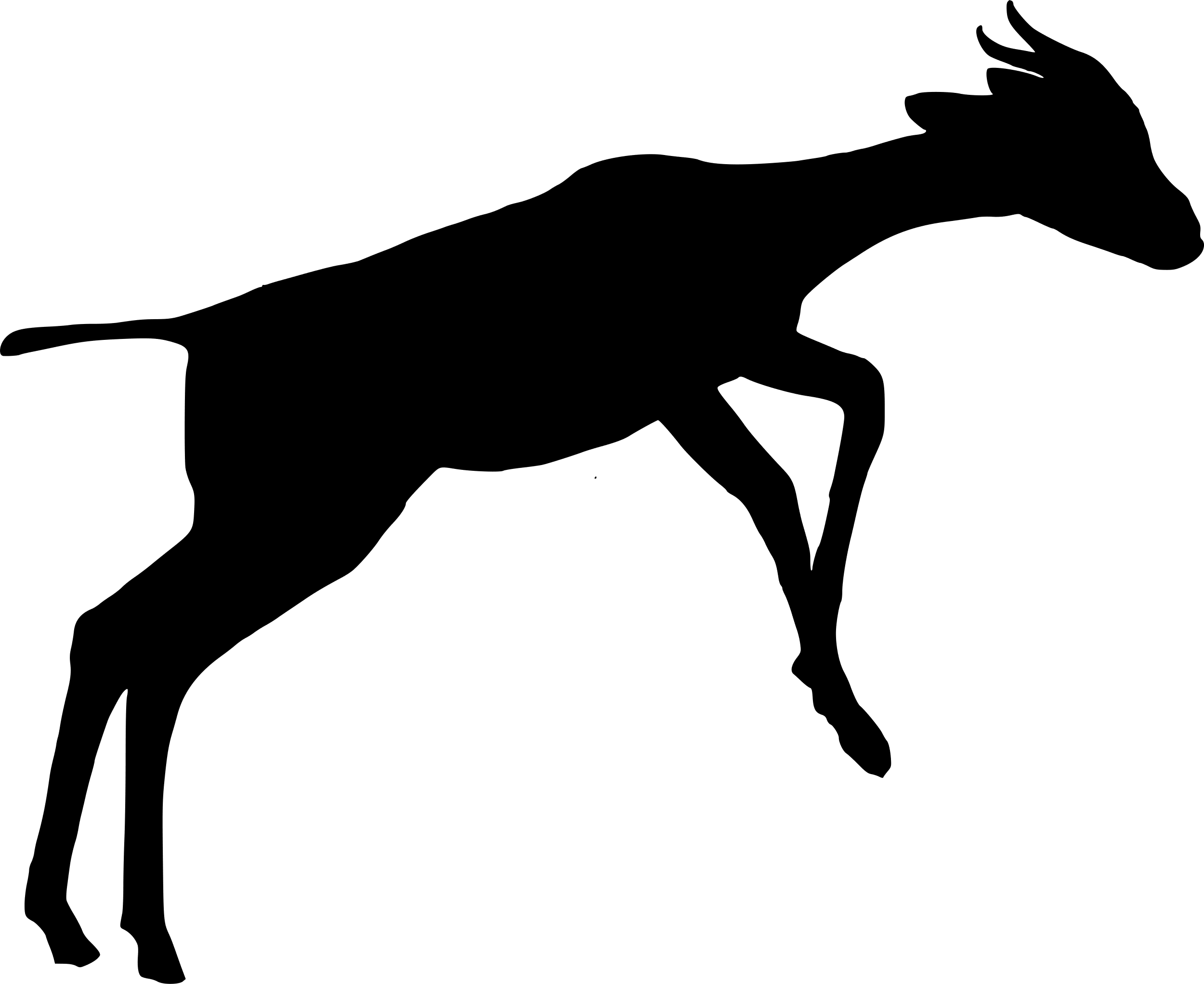

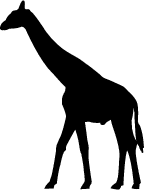

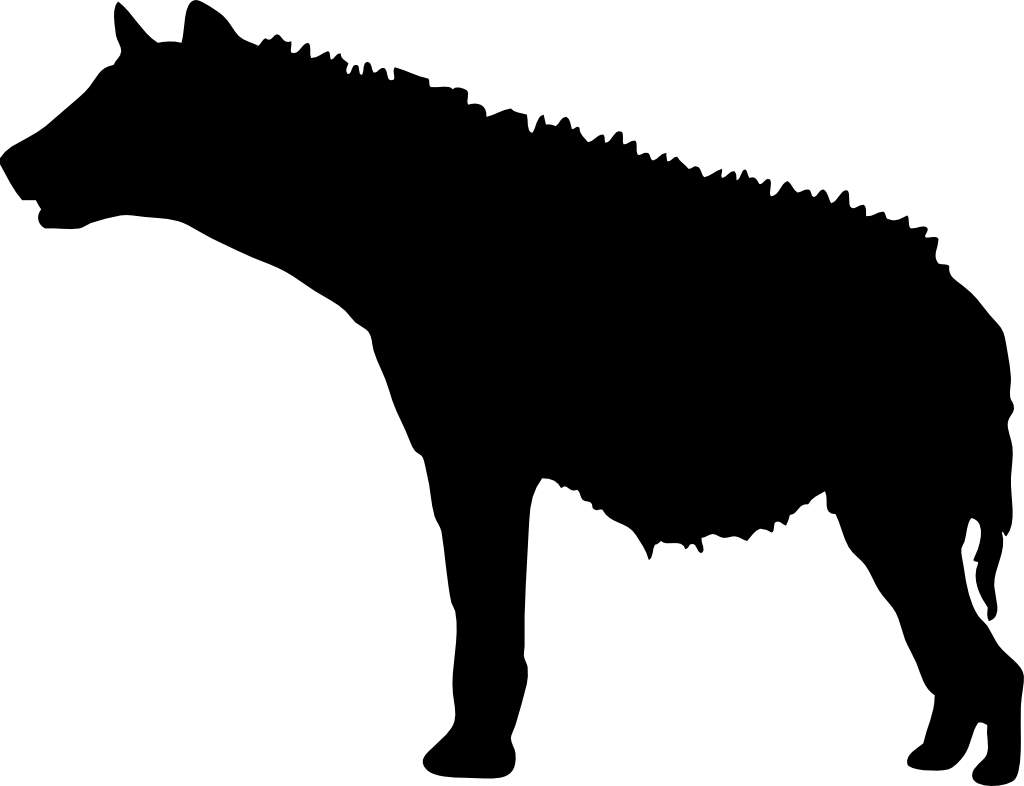

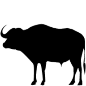

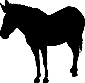

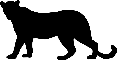

**Figure S2.** Marginal total effects of grid square-level *Opuntia* volume (standardised) on occupancy probability (ψ) for: **A)** olive baboon, **B)** vervet monkey, **C)** elephant, **D)** buffalo, **E)** dik-dik, **F)** impala, **G)** kudu, **H)** giraffe, **I)** Grevy’s zebra, **J)** plains zebra, **K)** spotted hyena, and **L)** leopard. The models assume that *Opuntia* indirectly affects occupancy through altering the composition of the native plant community; for model structure, see Figure 2 in main text. Shaded areas represent (from outside) 95%, 89%, 80%, 70%, 60%, and 50% compatibility intervals for the January-April (light green) and October-November (purple) seasons. Black lines indicate the posterior median marginal effect for the January-April (solid) and October-November (dashed) seasons.

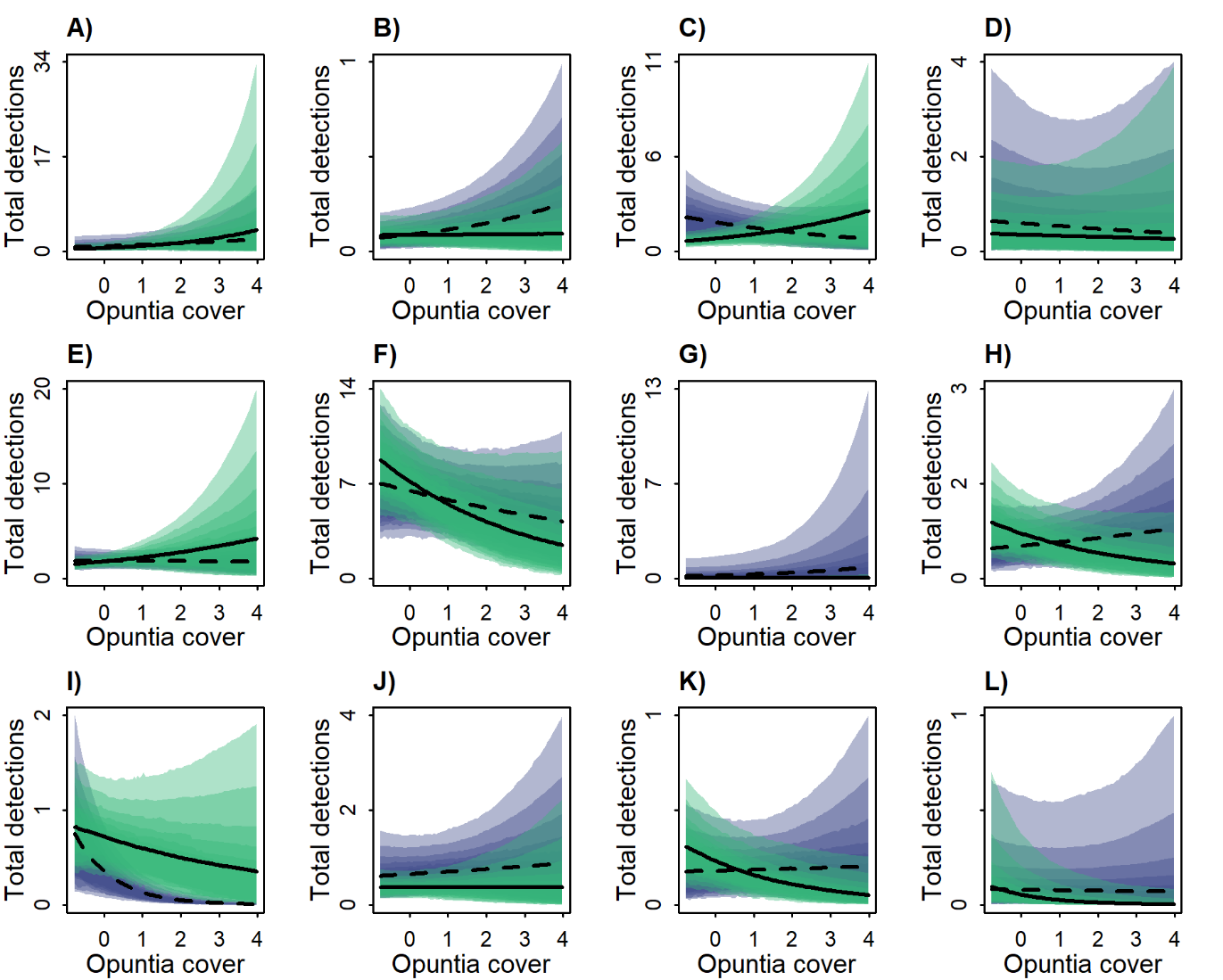

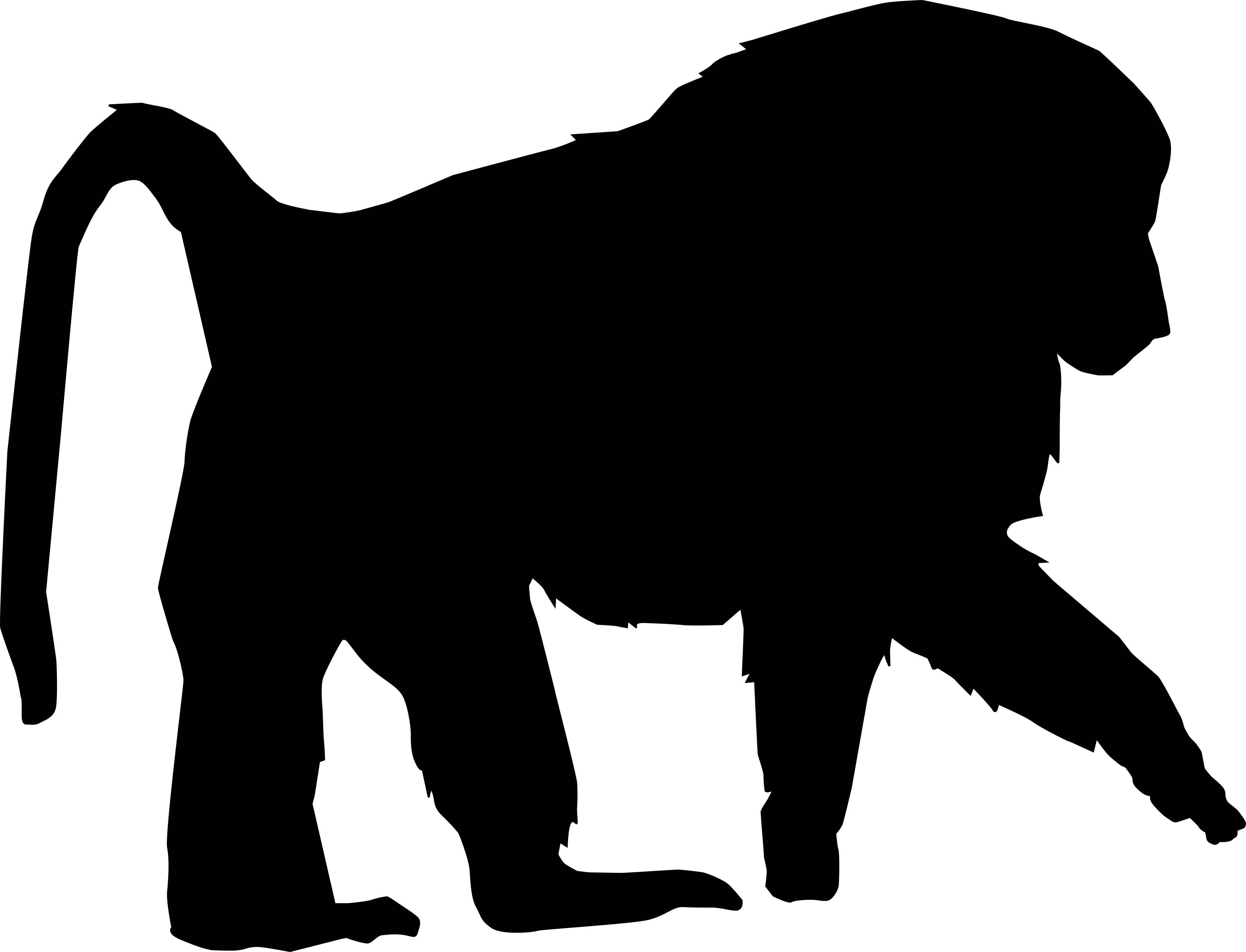

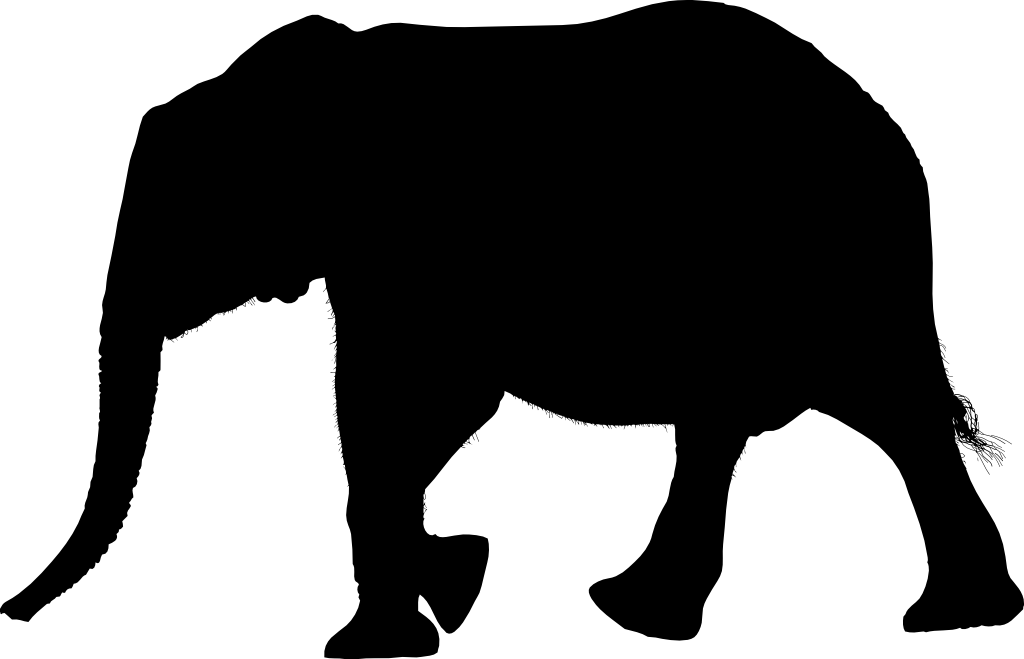

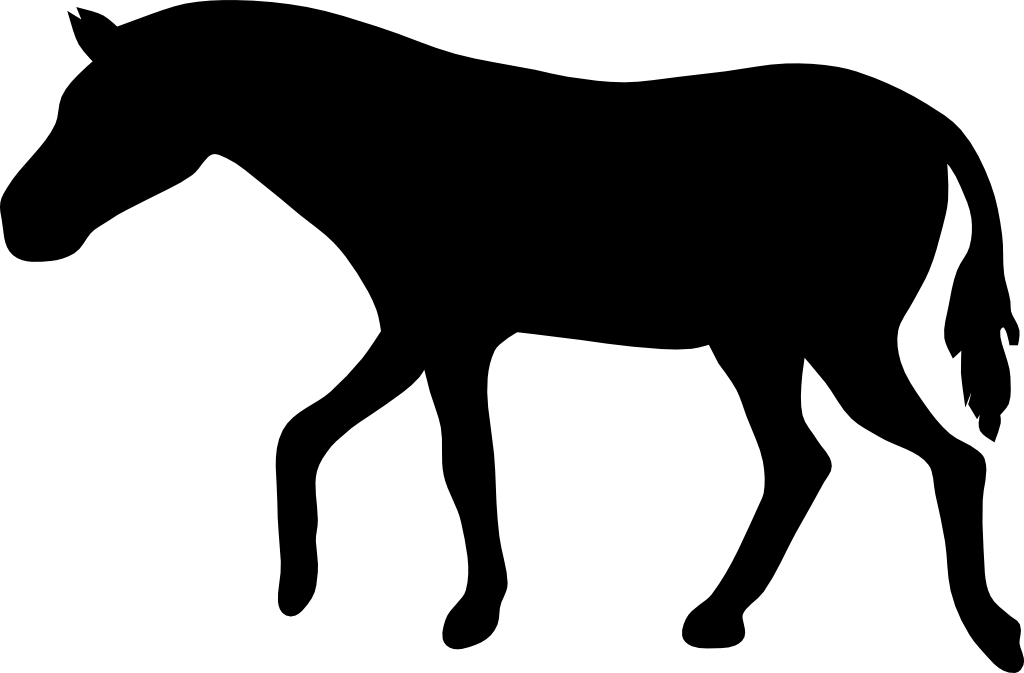

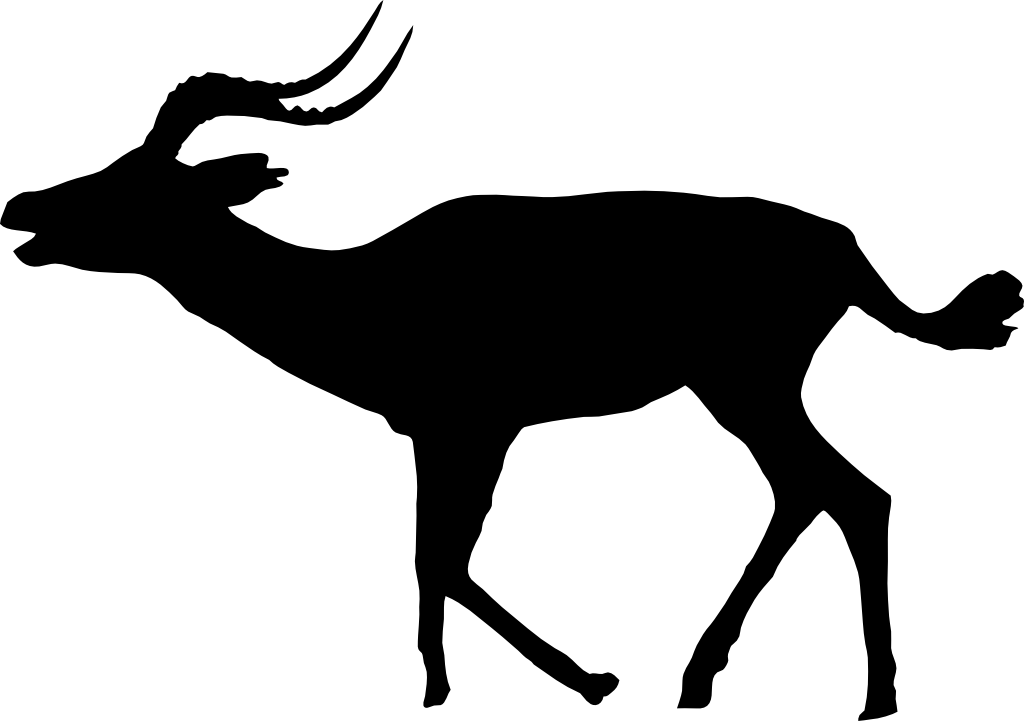

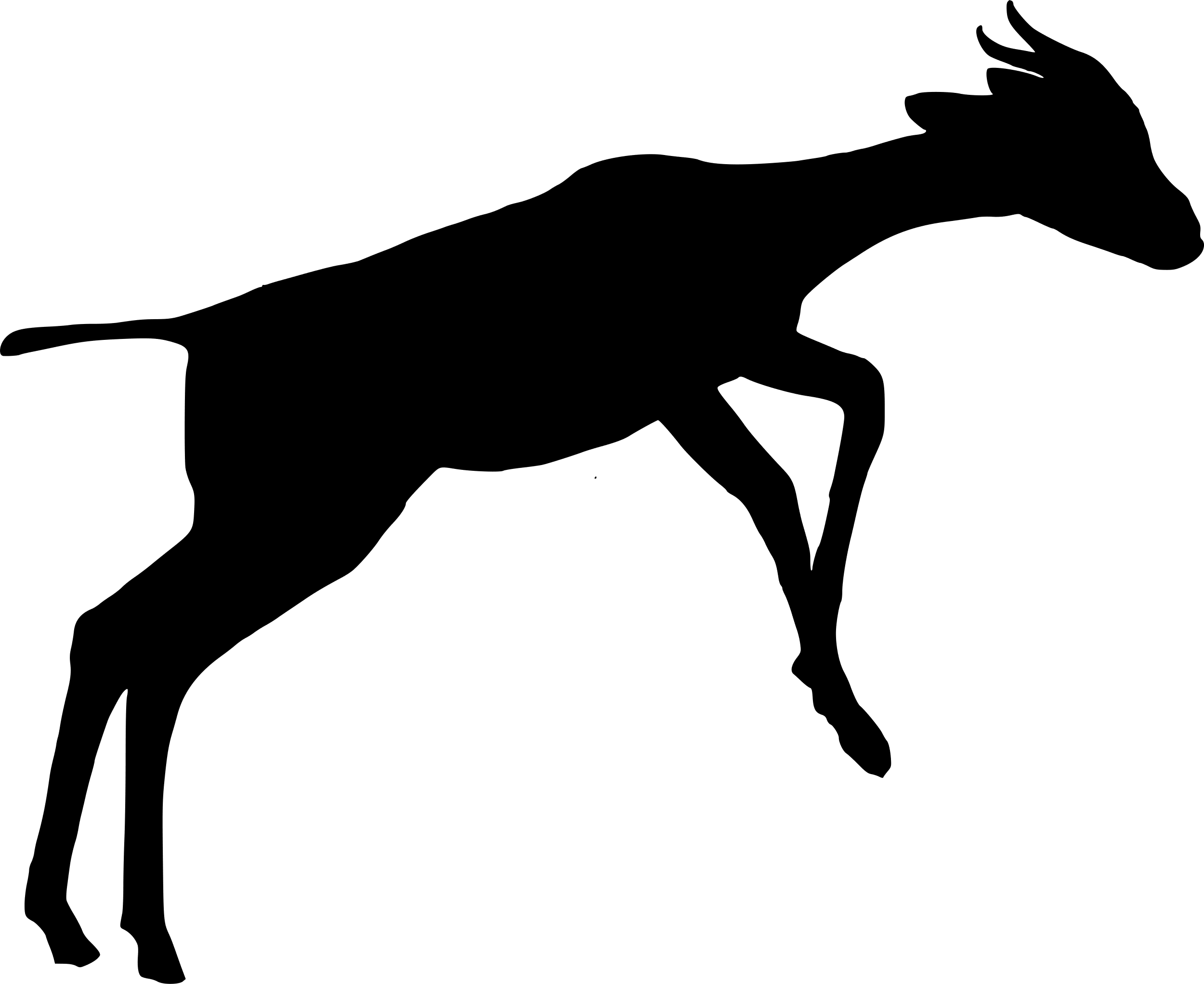

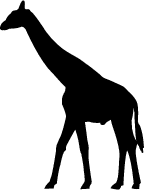

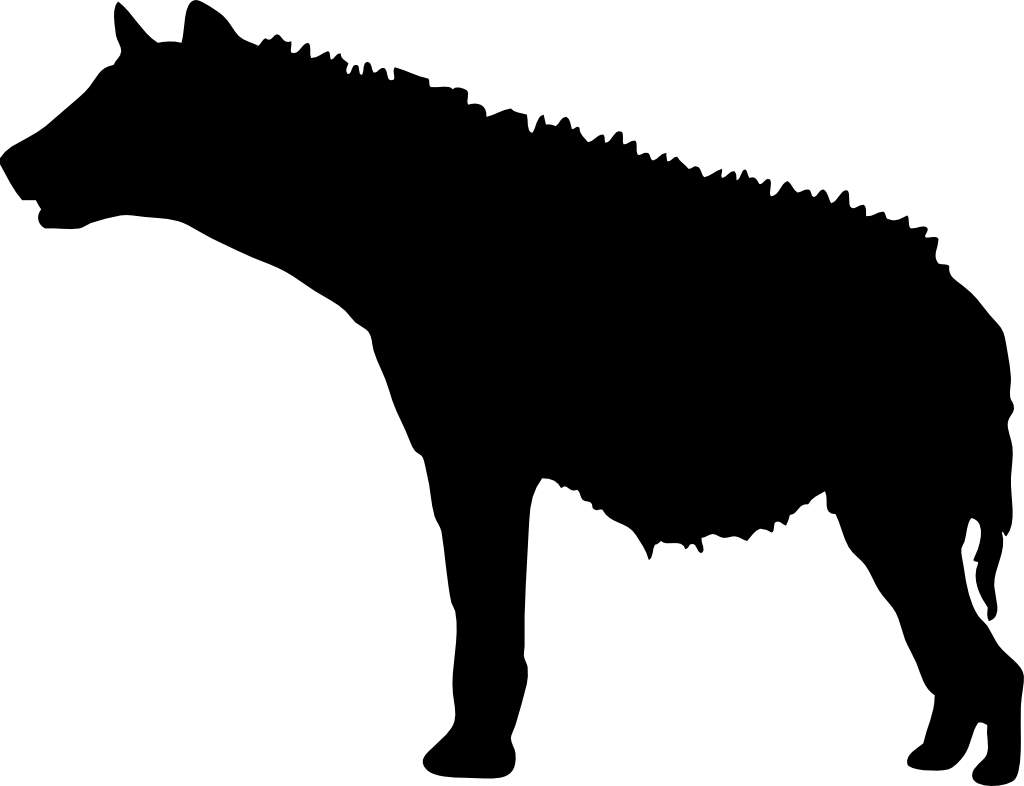

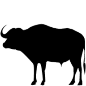

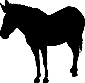

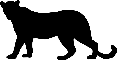

**Figure S3.** Marginal total effect of site-level *Opuntia* percentage cover (standardised) on the total number of detections per day for: **A)** olive baboon, **B)** vervet monkey, **C)** elephant, **D)** buffalo, **E)** dik-dik, **F)** impala, **G)** kudu, **H)** giraffe, **I)** Grevy’s zebra, **J)** plains zebra, **K)** spotted hyena, and **L)** leopard. The models assume that *Opuntia* indirectly affects occupancy through altering the composition of the native plant community; for model structure, see Figure 2 in main text. Shaded areas represent (from outside) 95%, 89%, 80%, 70%, 60%, and 50% compatibility intervals for the January-April (light green) and October-November (purple) seasons. Black lines indicate the posterior median marginal effect for the January-April (solid) and October-November (dashed) seasons.

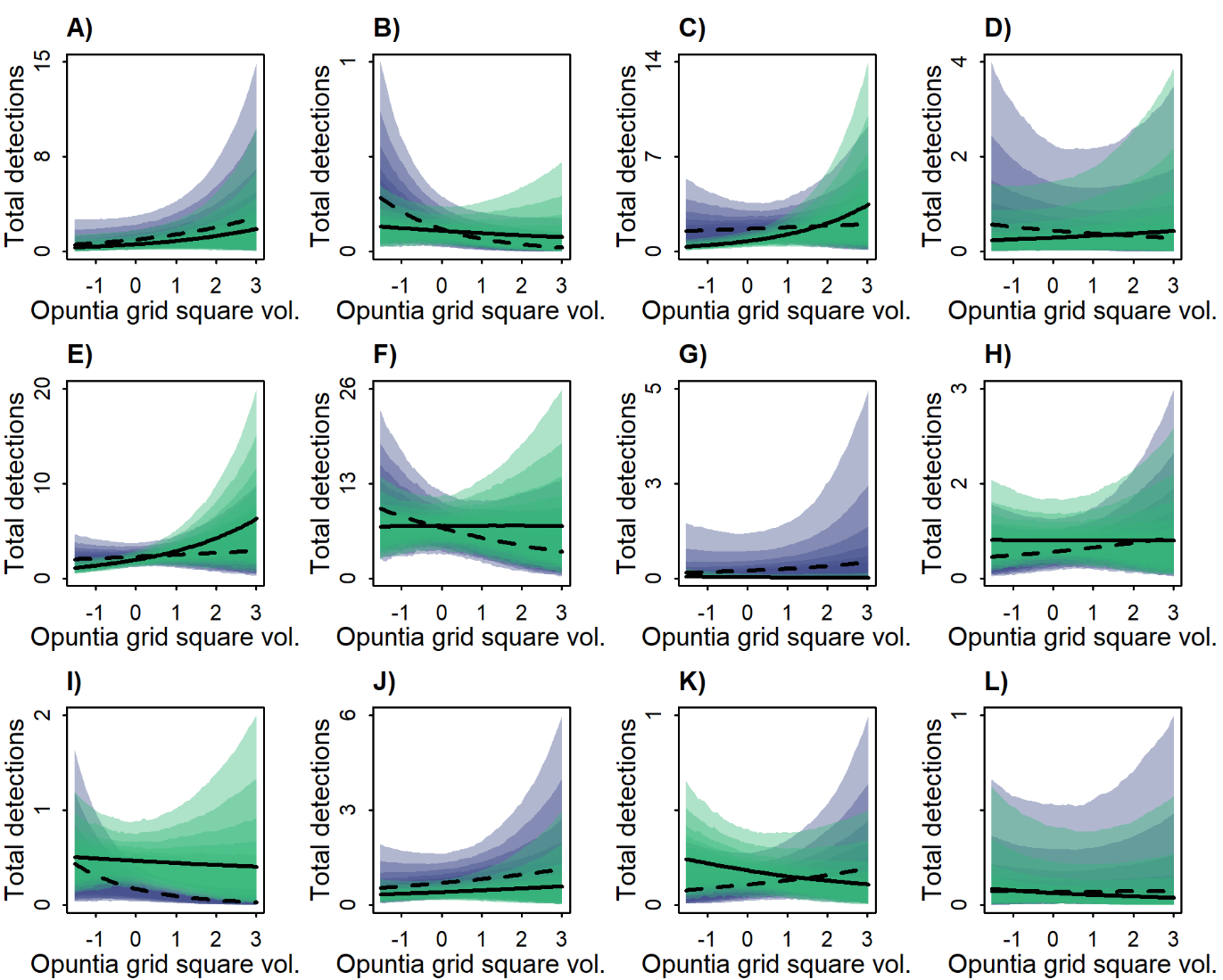

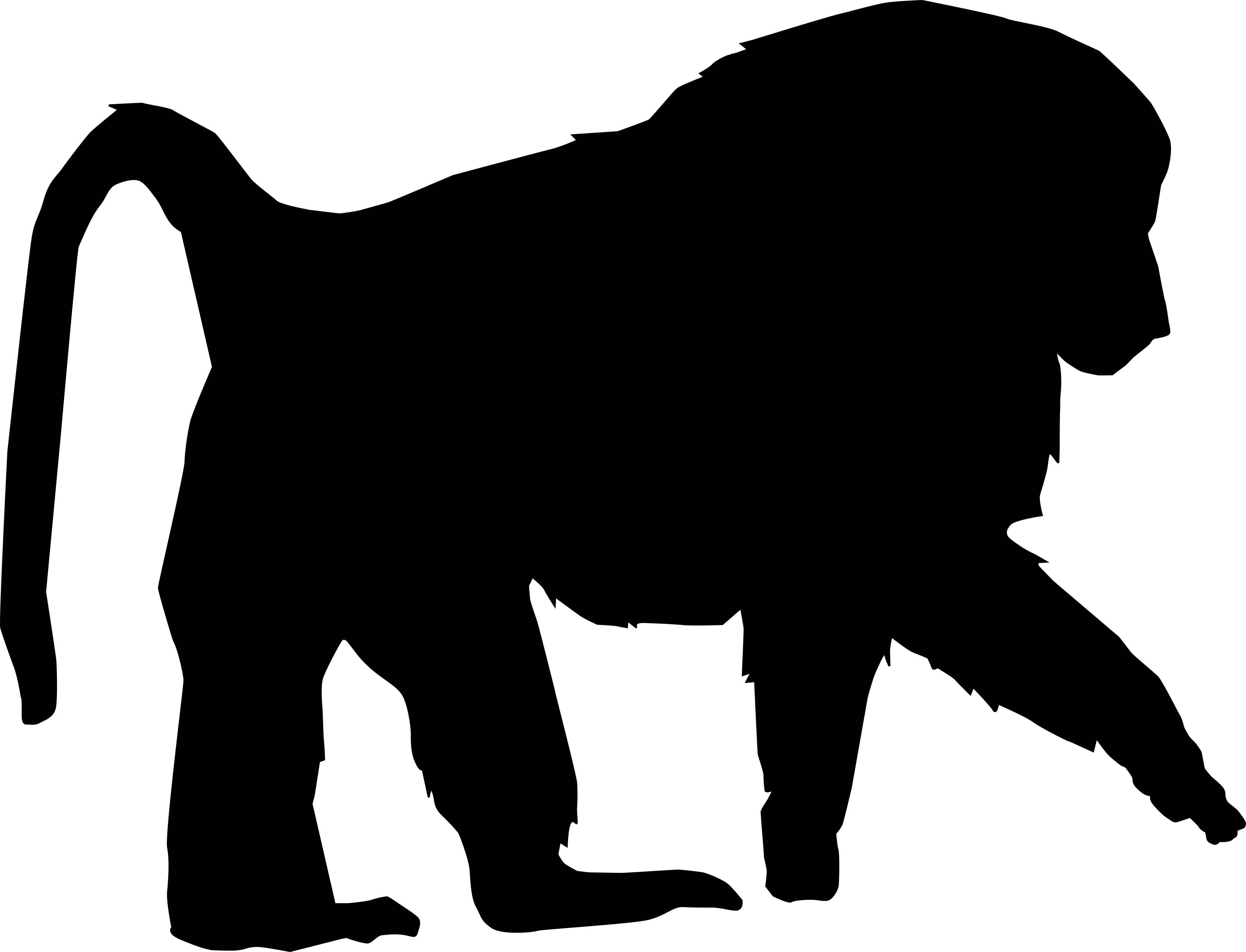

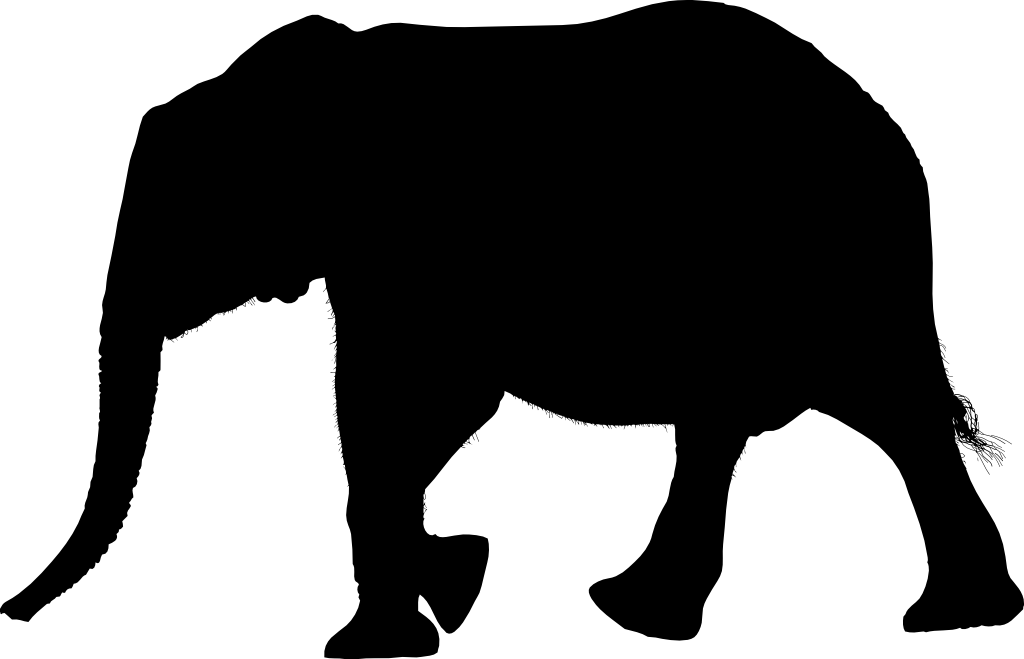

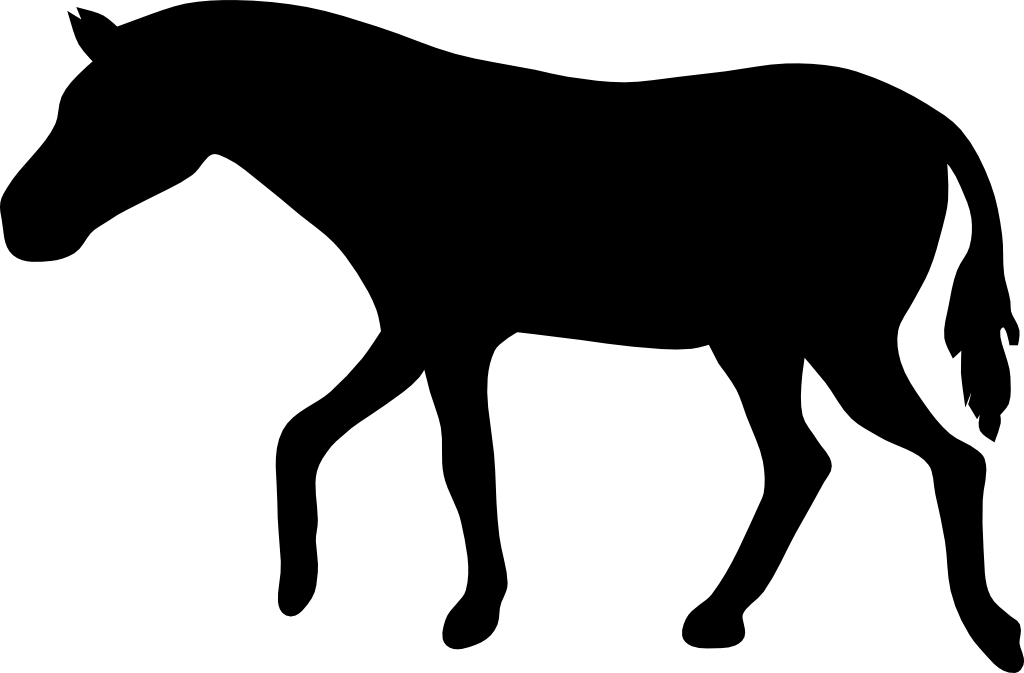

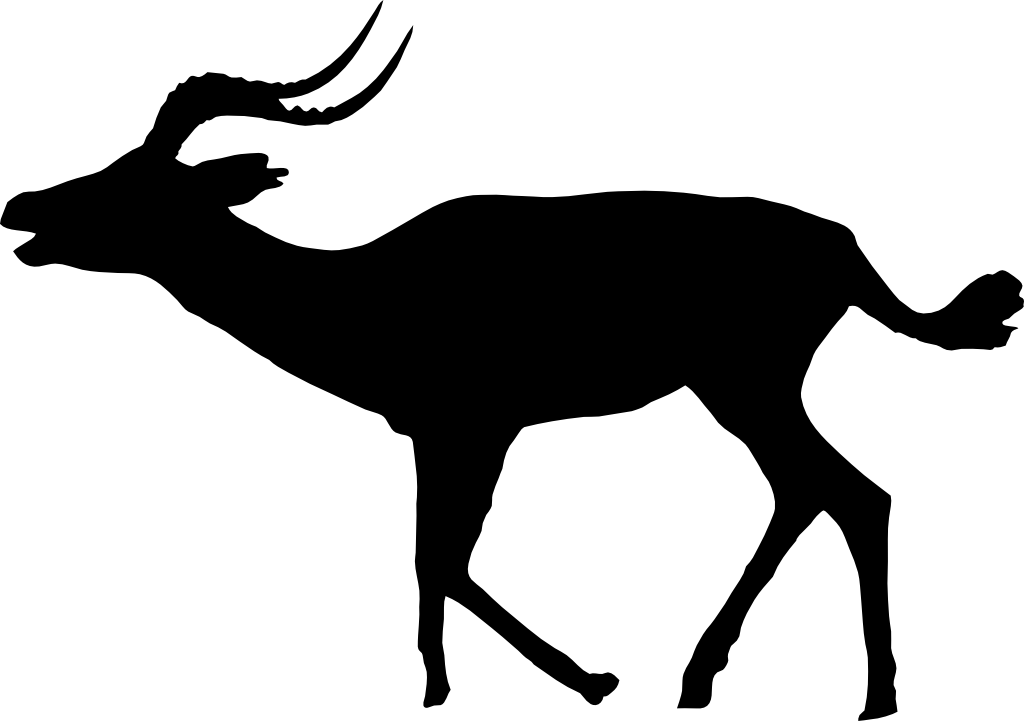

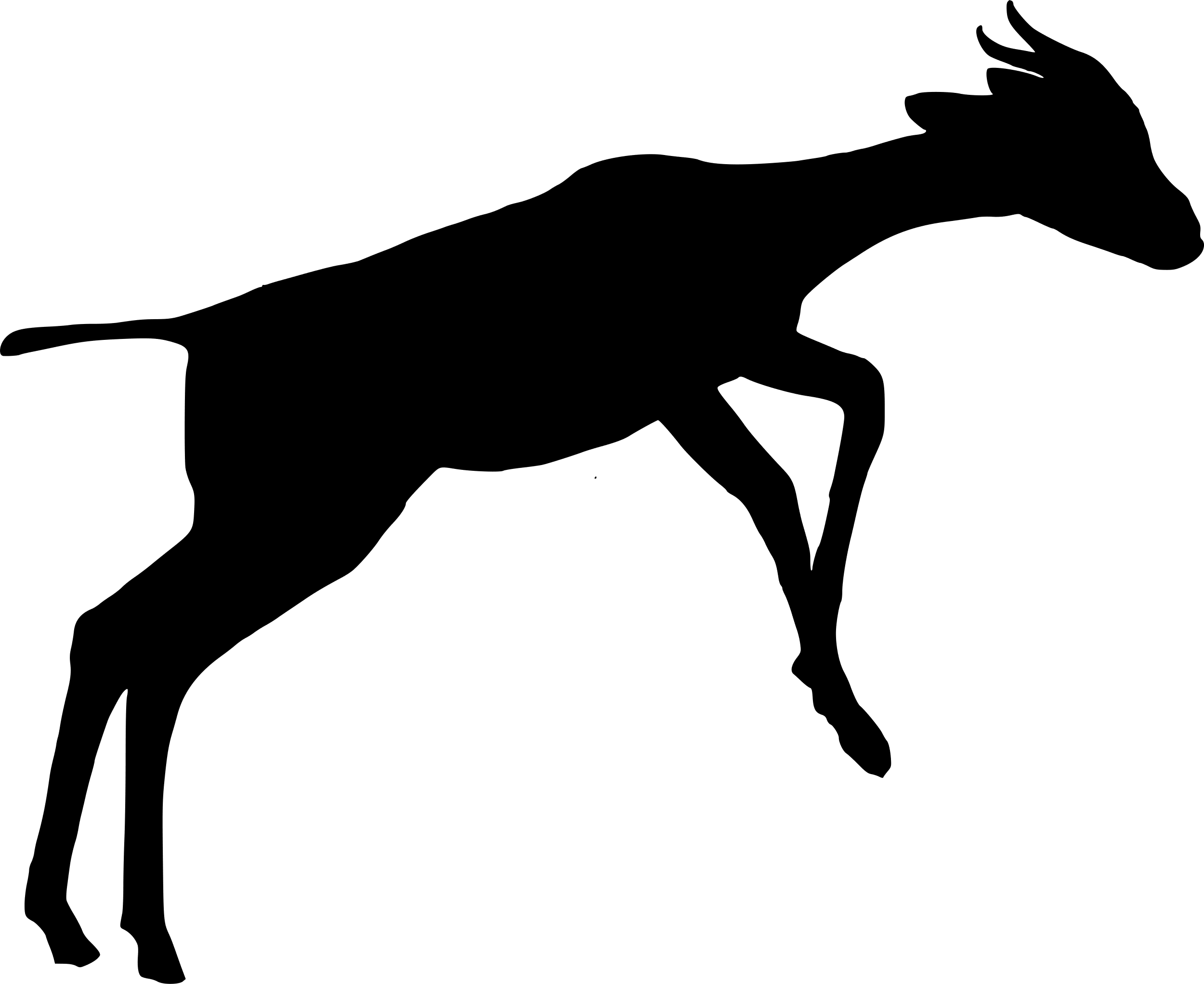

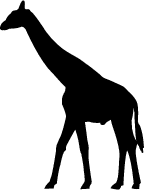

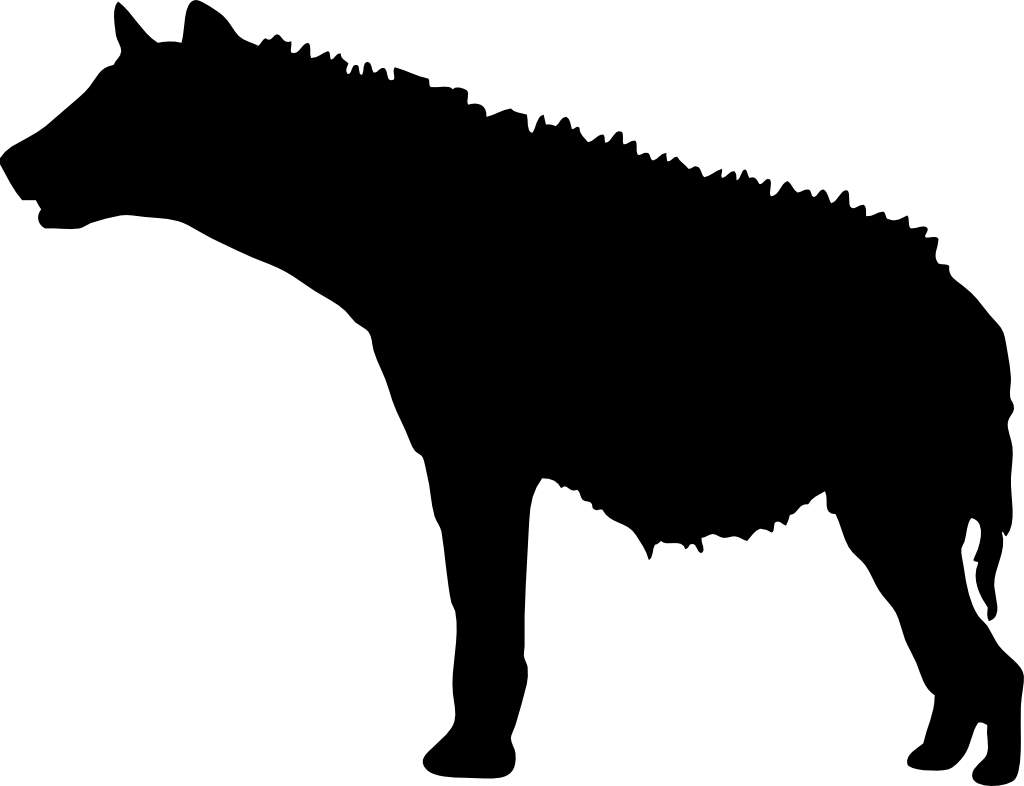

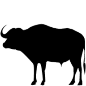

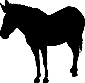

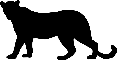

**Figure S4.** Marginal total effect of grid square-level *Opuntia* volume (standardised) on the total number of detections per day for: **A)** olive baboon, **B)** vervet monkey, **C)** elephant, **D)** buffalo, **E)** dik-dik, **F)** impala, **G)** kudu, **H)** giraffe, **I)** Grevy’s zebra, **J)** plains zebra, **K)** spotted hyena, and **L)** leopard. The models assume that *Opuntia* indirectly affects occupancy through altering the composition of the native plant community; for model structure, see Figure 2 in main text. Shaded areas represent (from outside) 95%, 89%, 80%, 70%, 60%, and 50% compatibility intervals for the January-April (light green) and October-November (purple) seasons. Black lines indicate the posterior median marginal effect for the January-April (solid) and October-November (dashed) seasons.

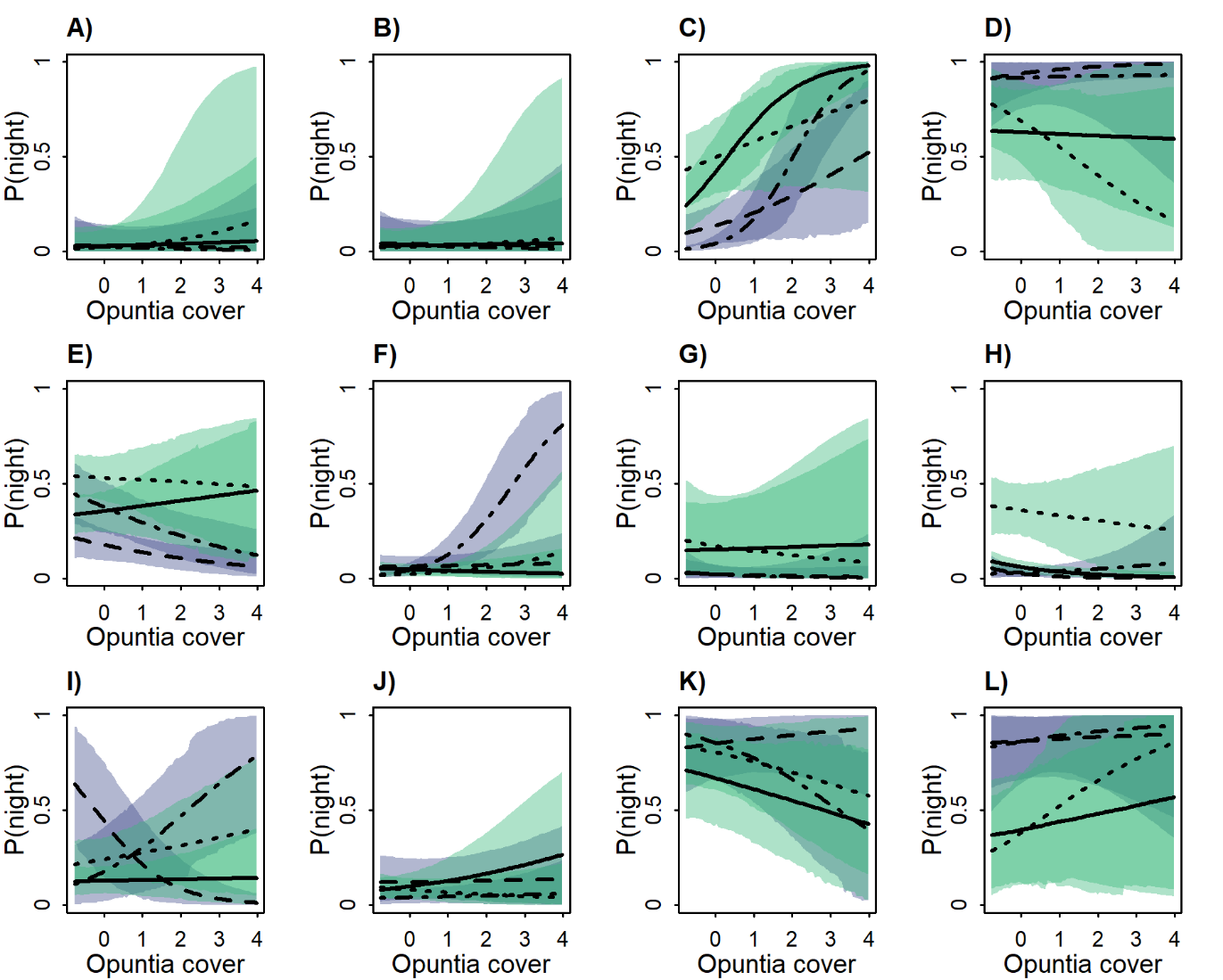

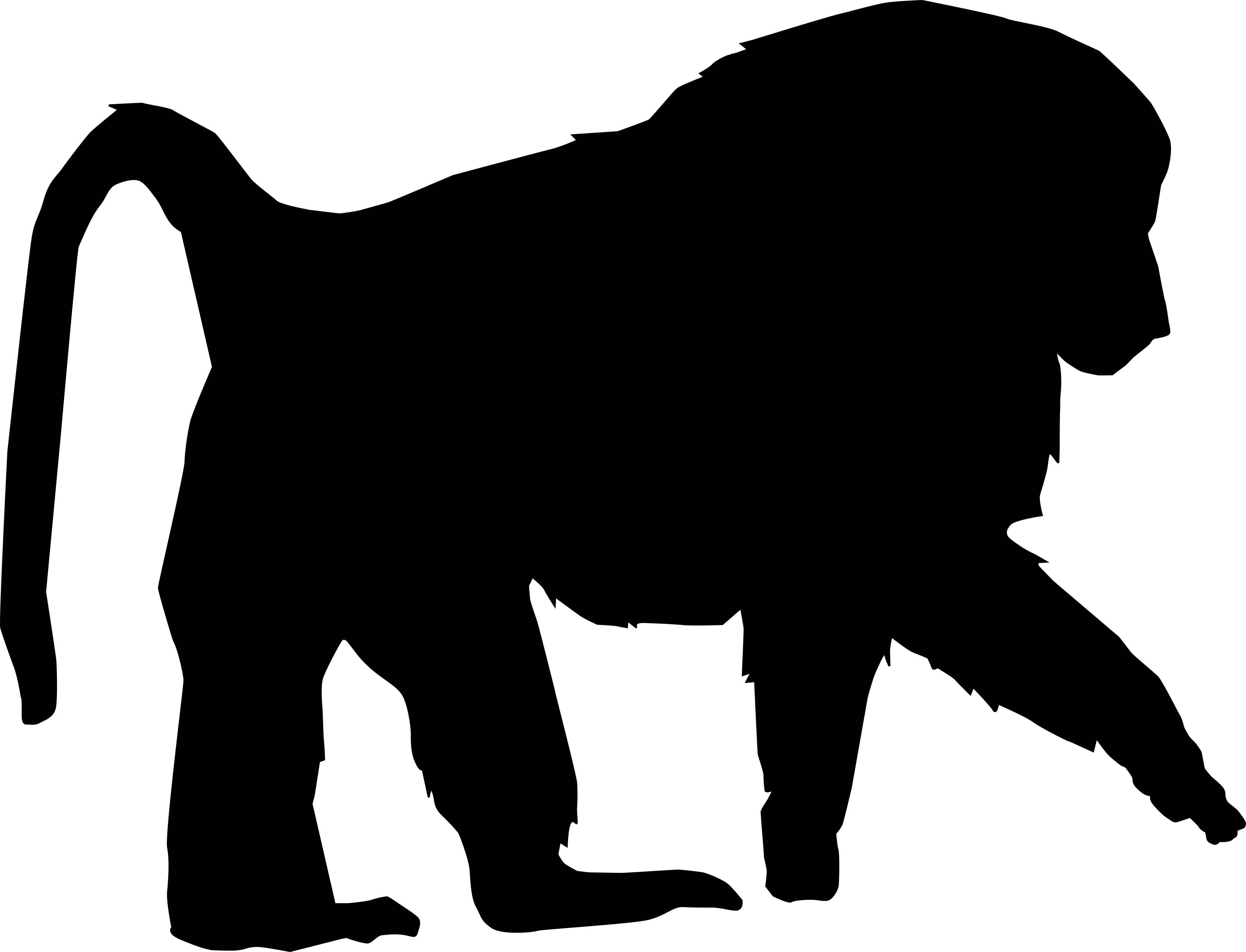

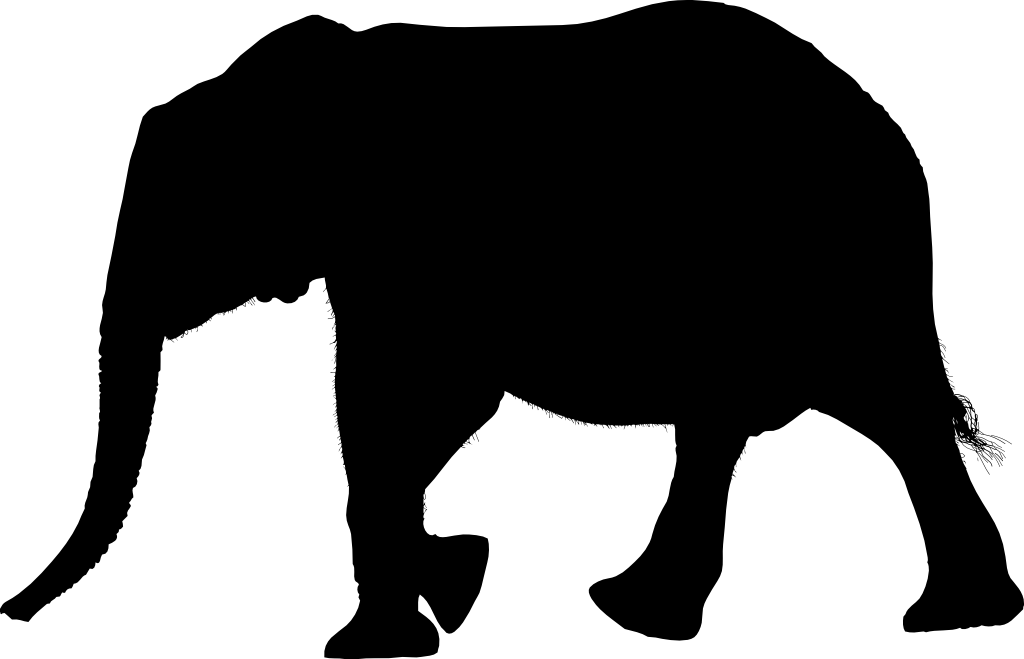

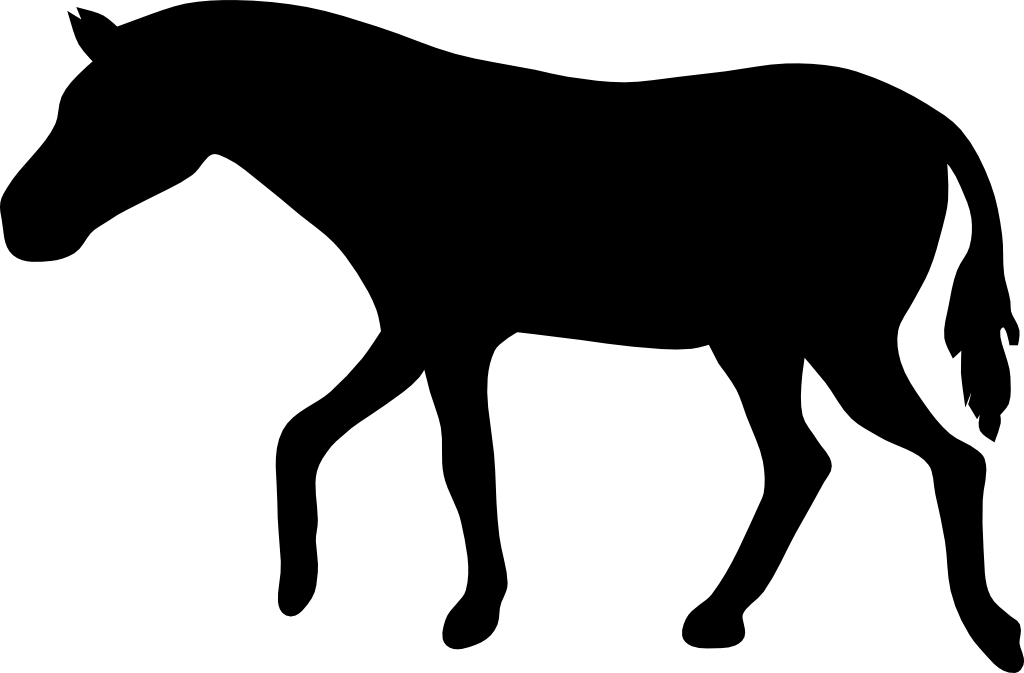

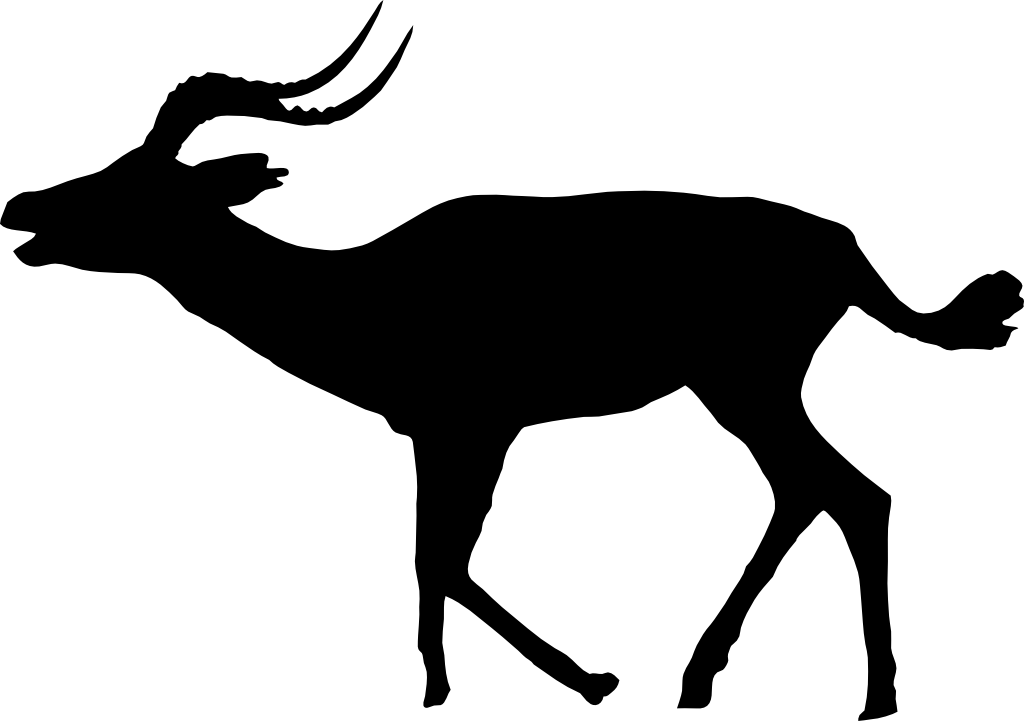

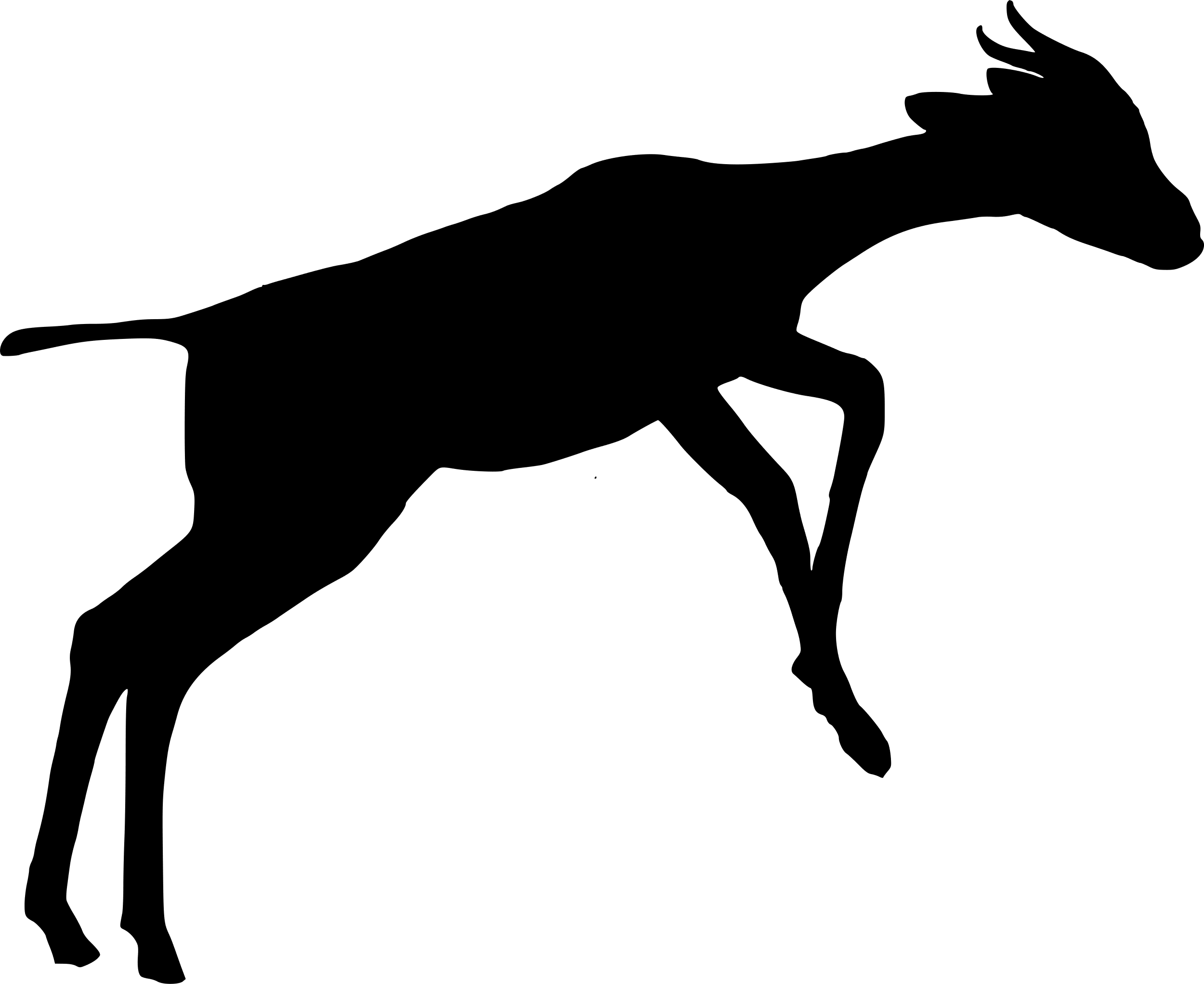

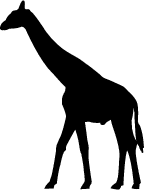

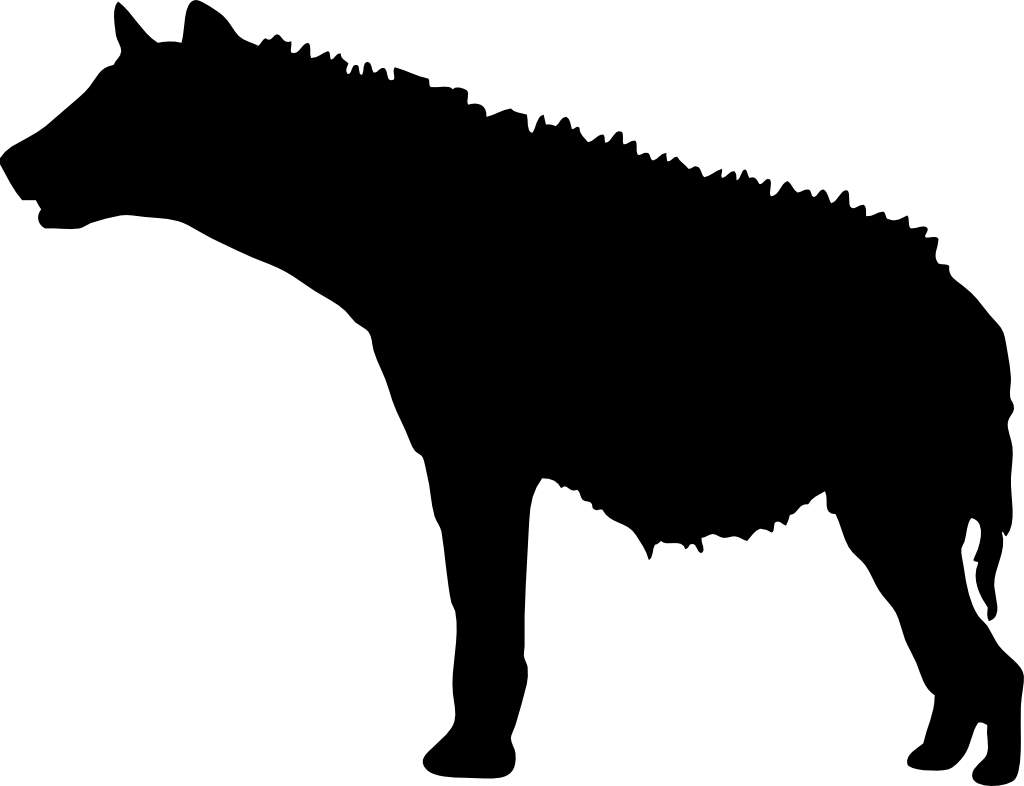

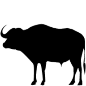

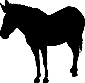

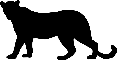

**Figure S5.** Marginal total effect of site-level *Opuntia* percentage cover (standardised) on the proportion of detections occurring at night for: **A)** olive baboon, **B)** vervet monkey, **C)** elephant, **D)** buffalo, **E)** dik-dik, **F)** impala, **G)** kudu, **H)** giraffe, **I)** Grevy’s zebra, **J)** plains zebra, **K)** spotted hyena, and **L)** leopard. The models assume that *Opuntia* does not indirectly affect occupancy through altering the composition of the native plant community; for model structure, see Figure 2 in main text. Shaded areas represent 89 compatibility intervals for the January-April (light green) and October-November (purple) seasons. Black lines indicate posterior median marginal effects for January-April under a new moon (―) and full moon (‧ ‧ ‧), and October-November under a new moon (– – –) and full moon (– ‧ –).

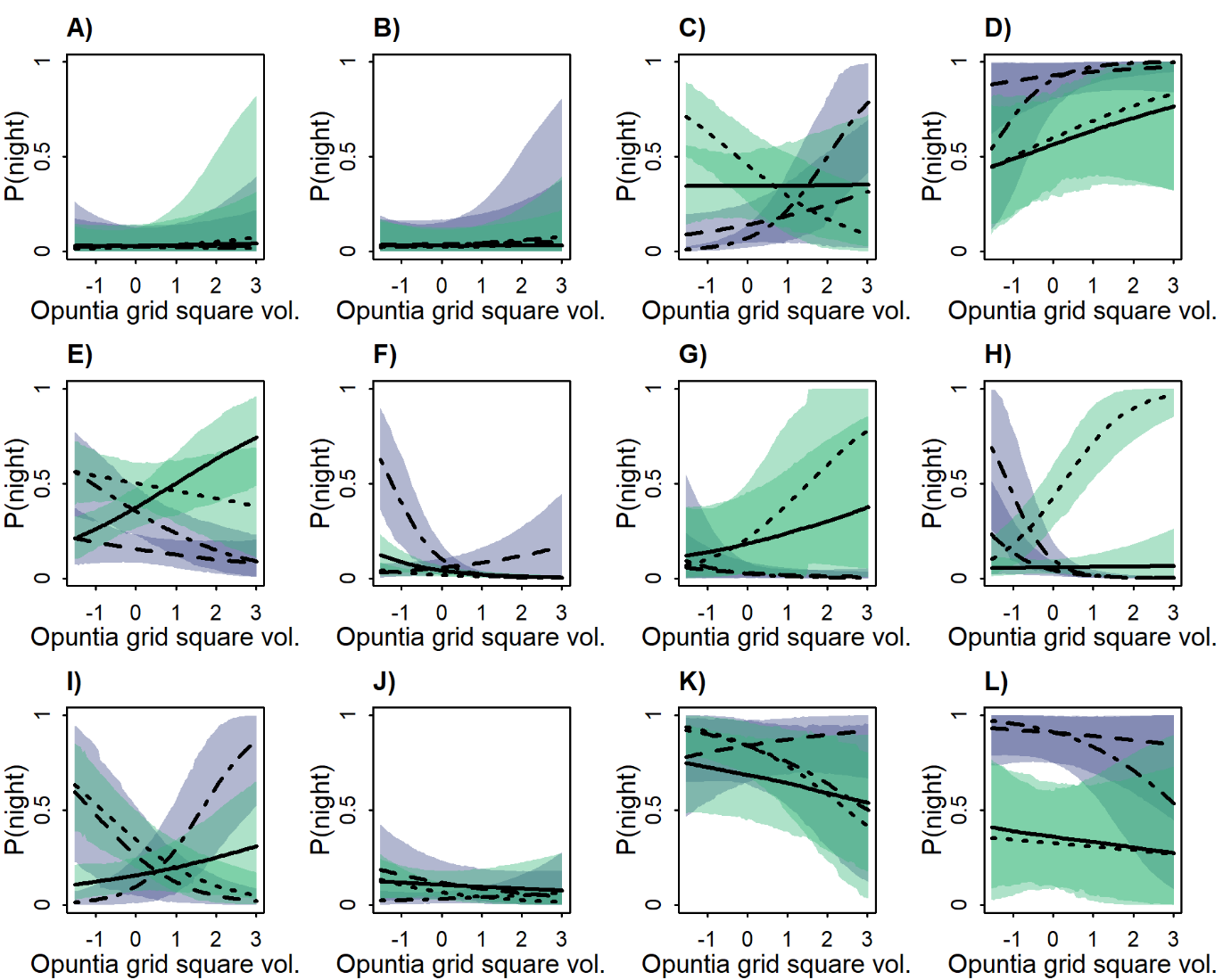

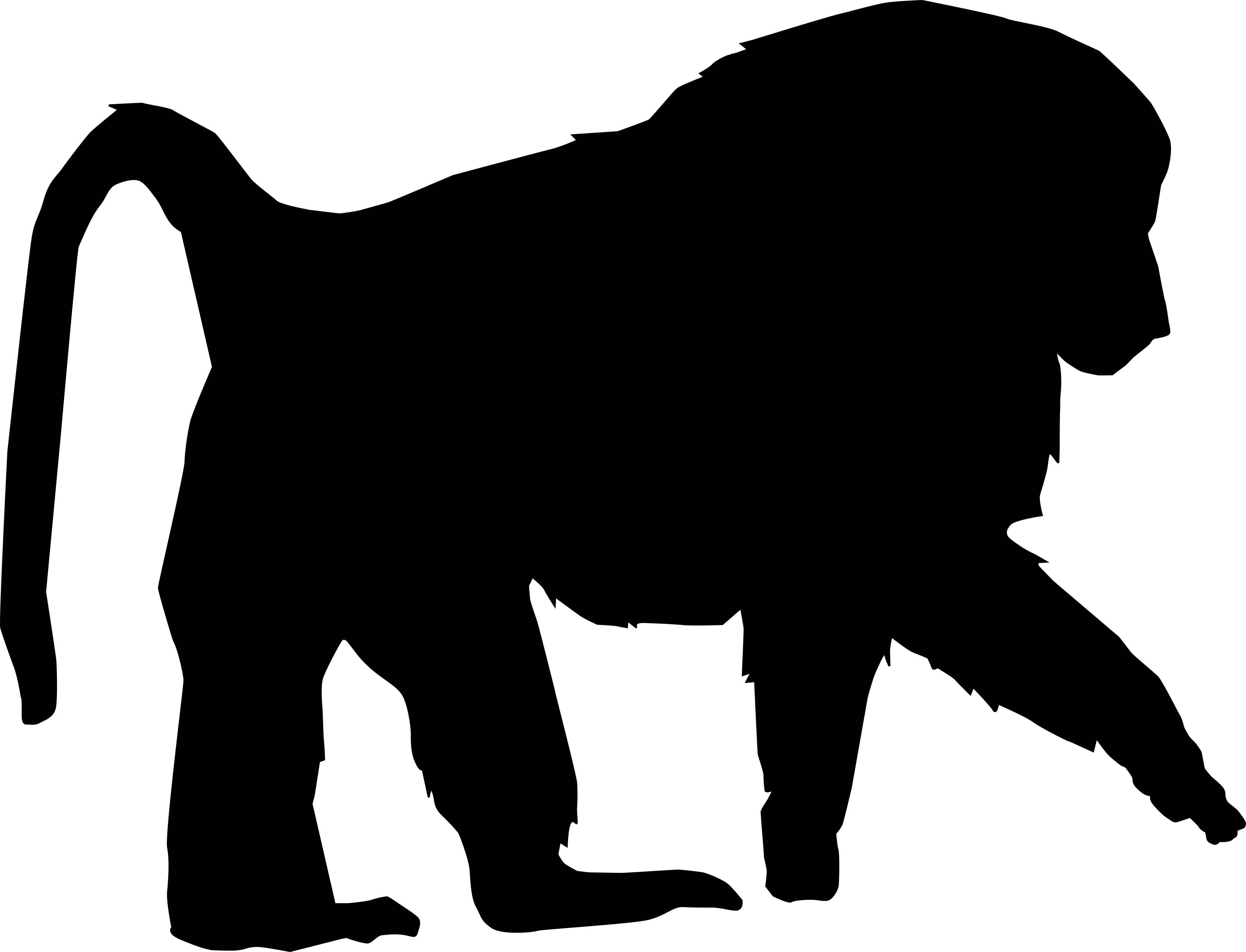

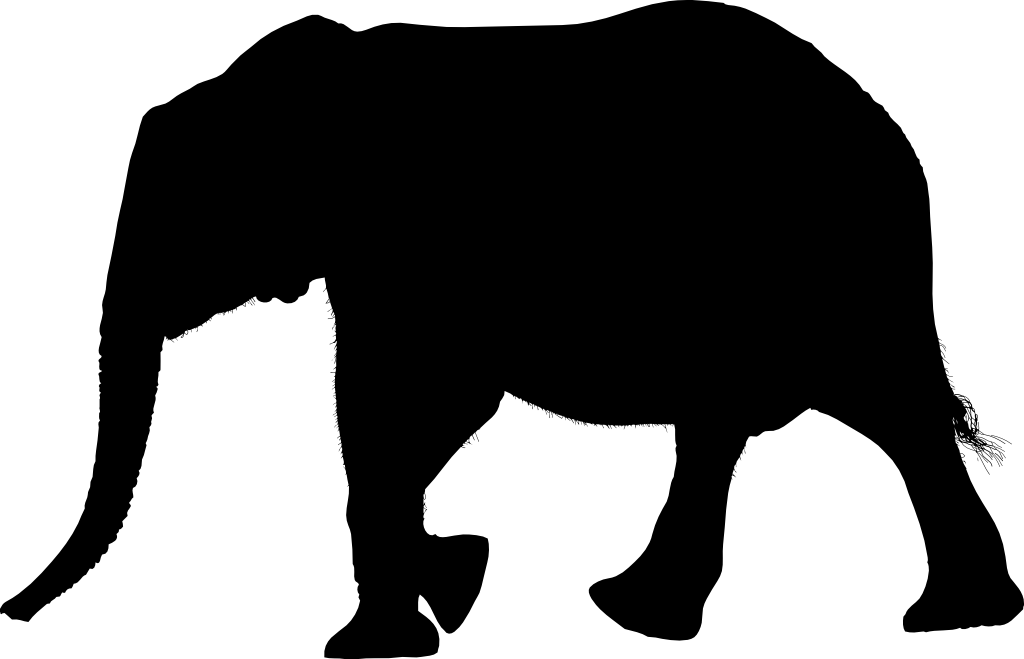

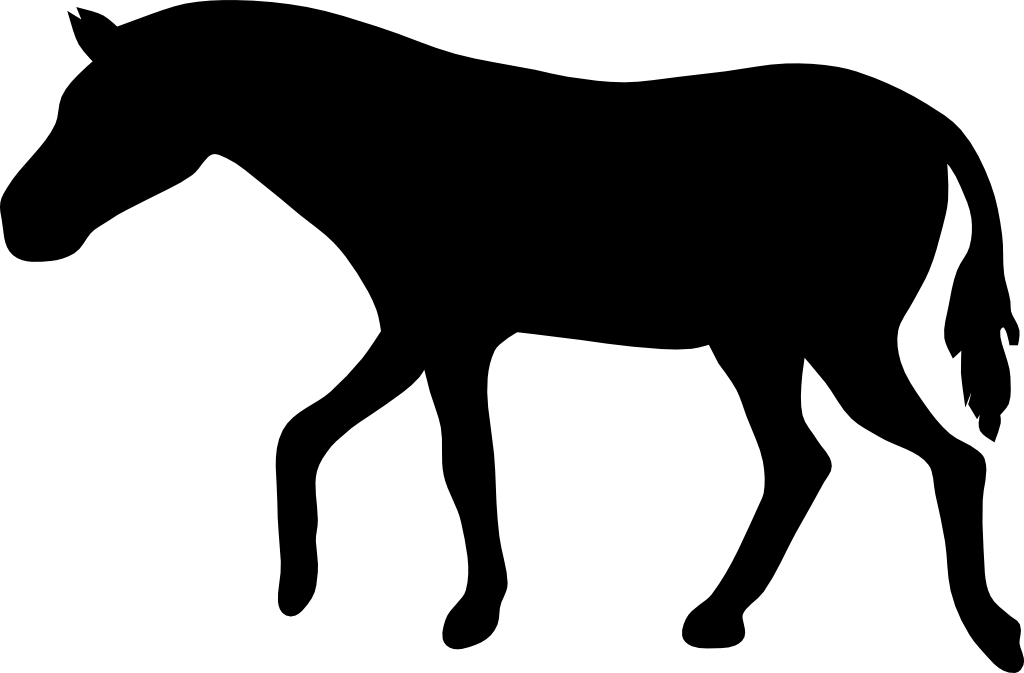

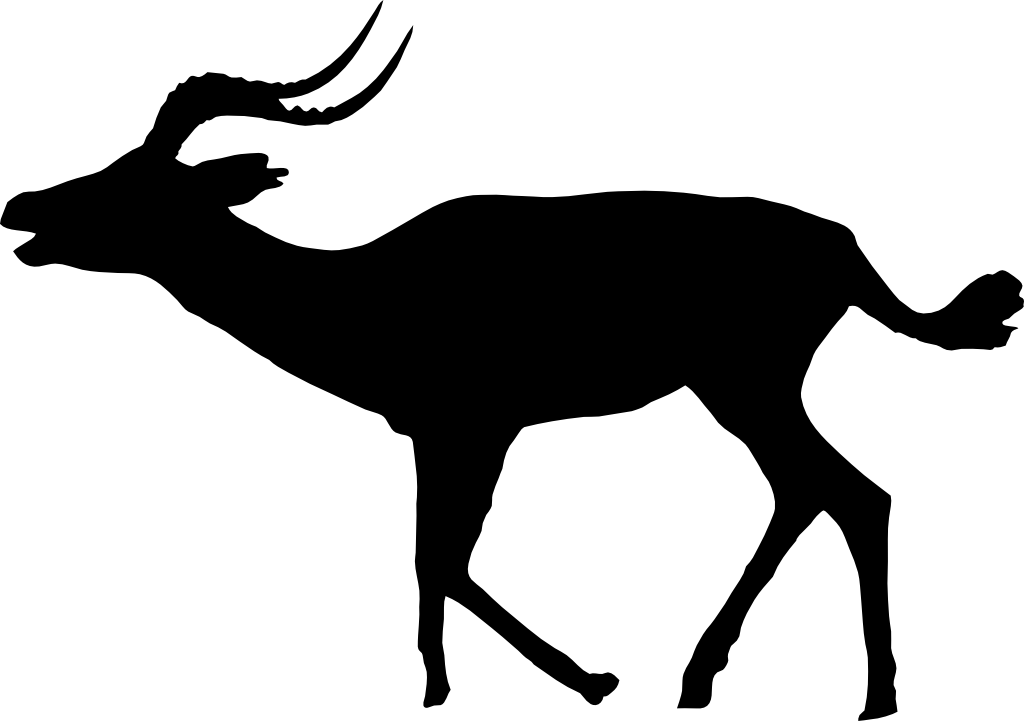

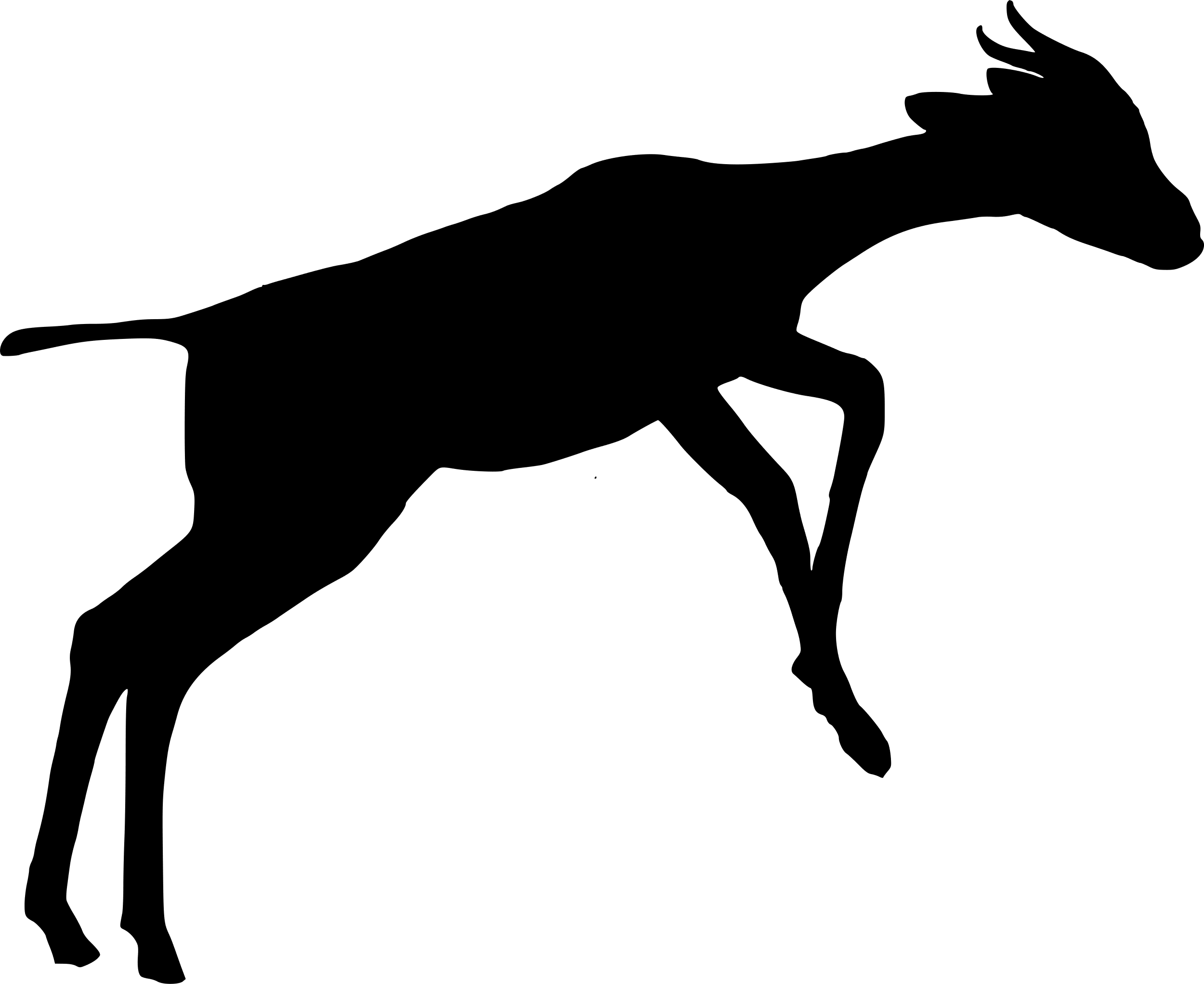

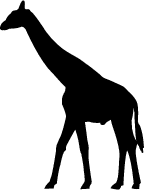

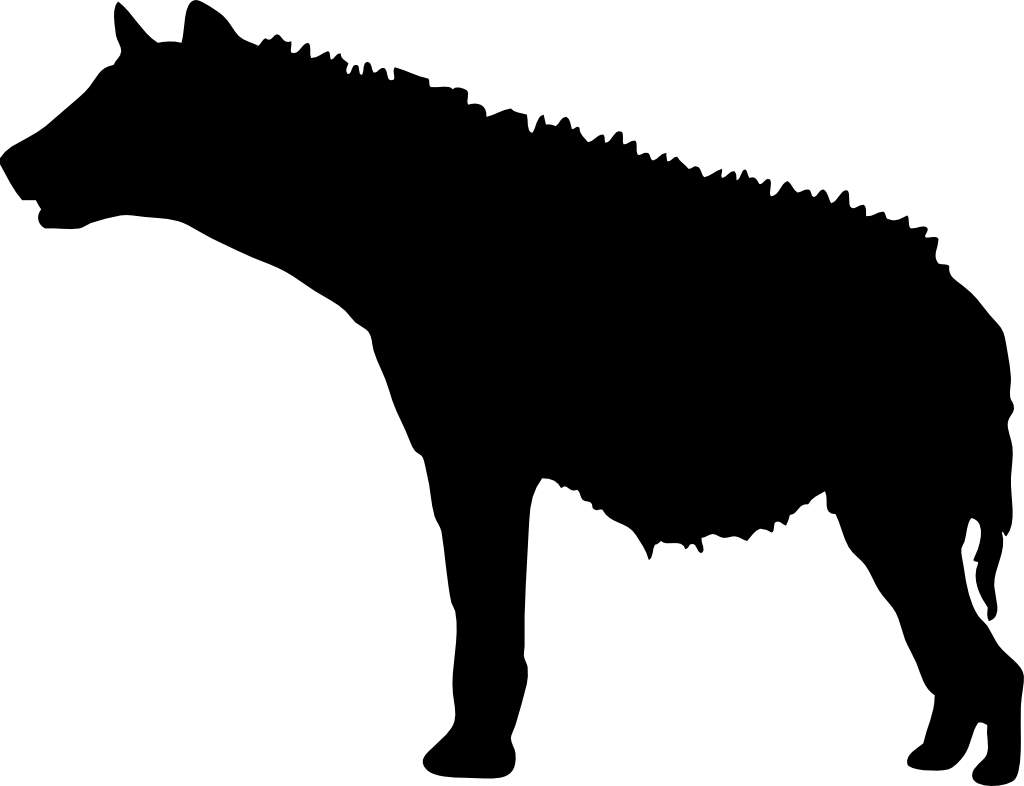

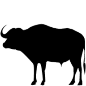

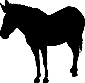

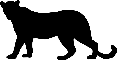

**Figure S6.** Marginal total effect of grid square-level *Opuntia* volume (standardised) on the proportion of detections occurring at night for: **A)** olive baboon, **B)** vervet monkey, **C)** elephant, **D)** buffalo, **E)** dik-dik, **F)** impala, **G)** kudu, **H)** giraffe, **I)** Grevy’s zebra, **J)** plains zebra, **K)** spotted hyena, and **L)** leopard. The models assume that *Opuntia* does not indirectly affect occupancy through altering the composition of the native plant community; for model structure, see Figure 2 in main text. Shaded areas represent 89 compatibility intervals for the January-April (light green) and October-November (purple) seasons. Black lines indicate posterior median marginal effects for January-April under a new moon (―) and full moon (‧ ‧ ‧), and October-November under a new moon (– – –) and full moon	(– ‧ –).

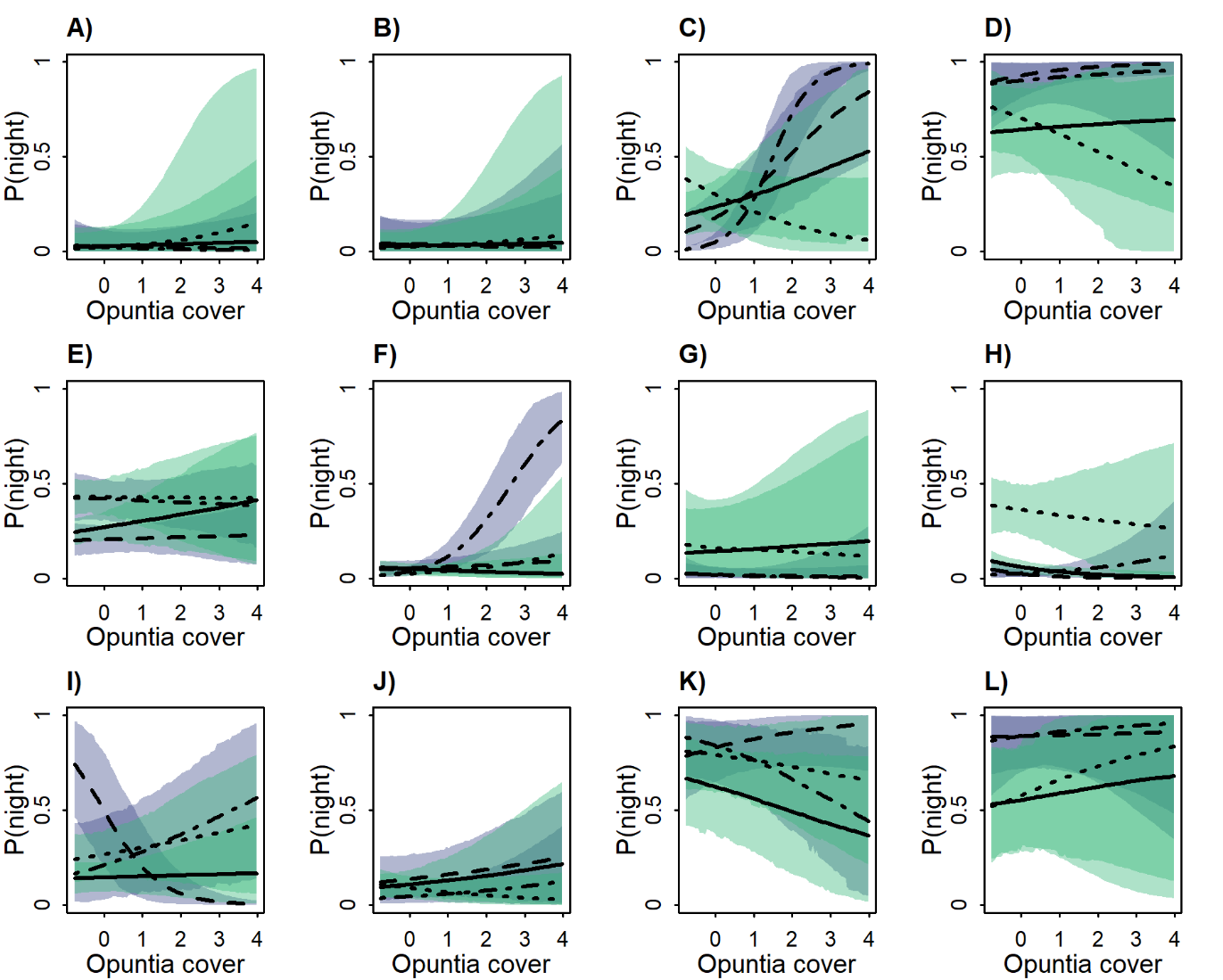

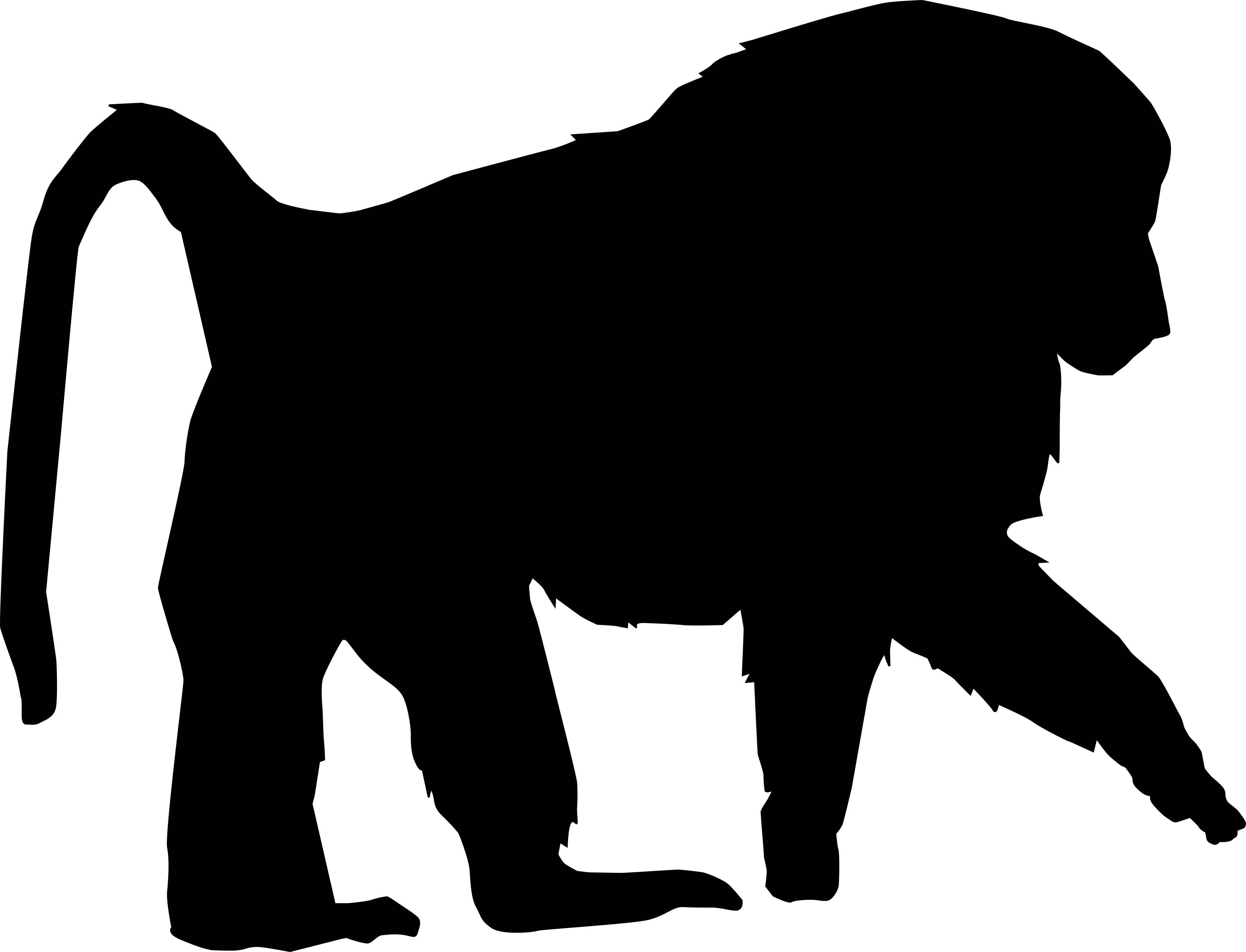

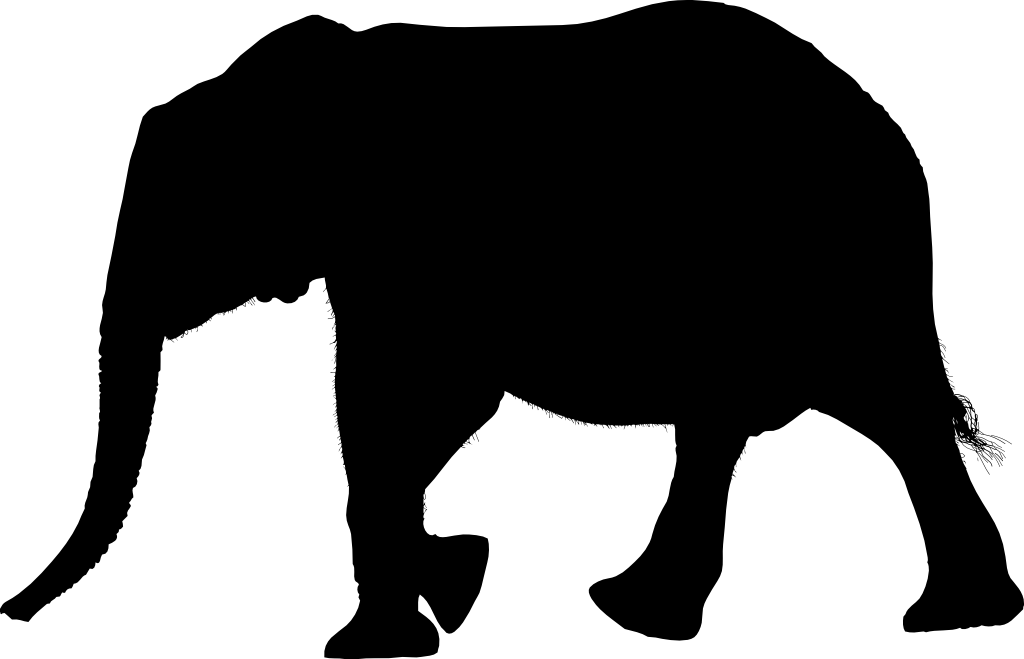

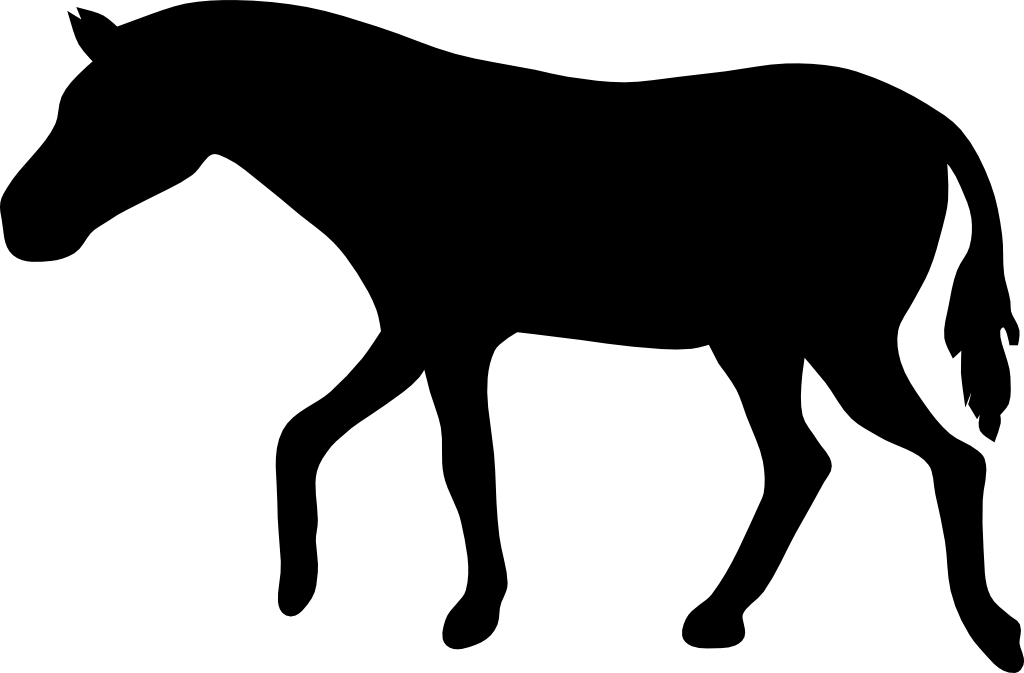

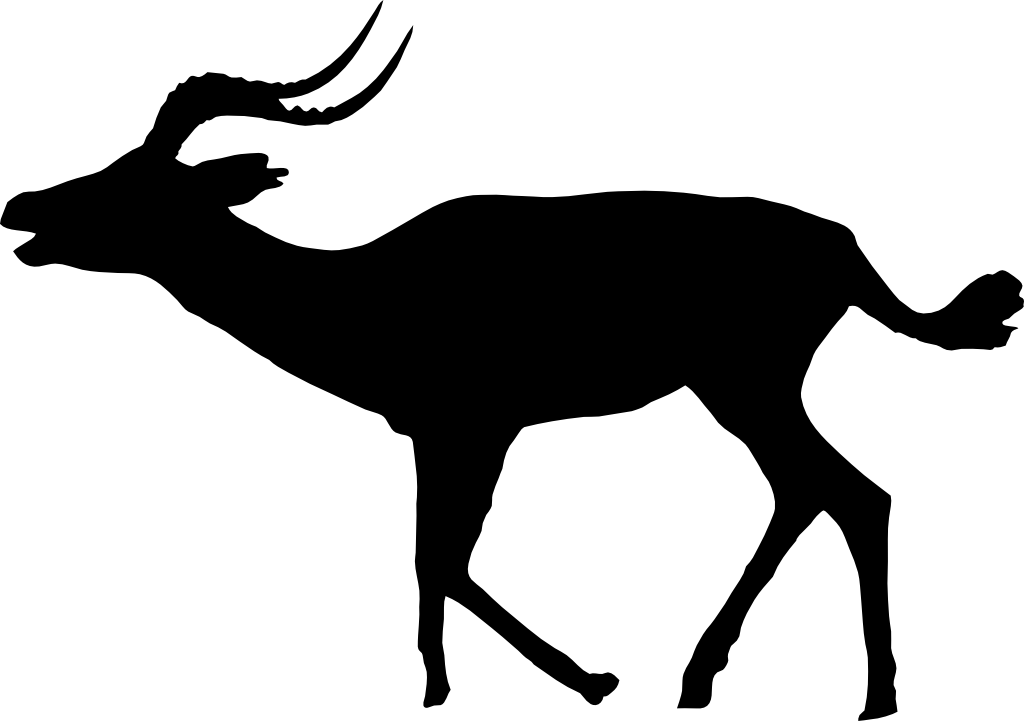

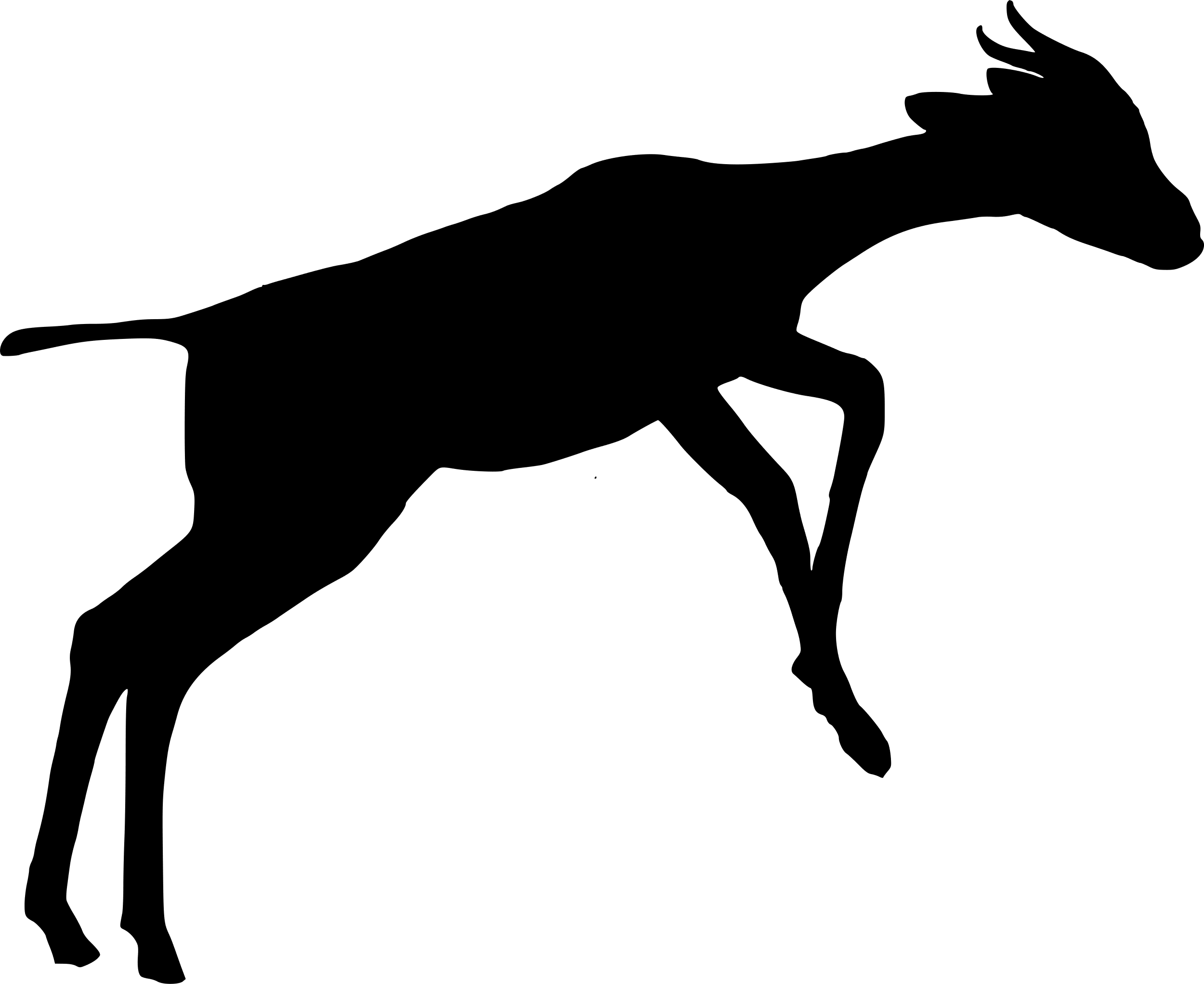

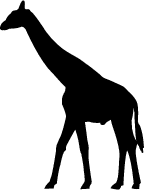

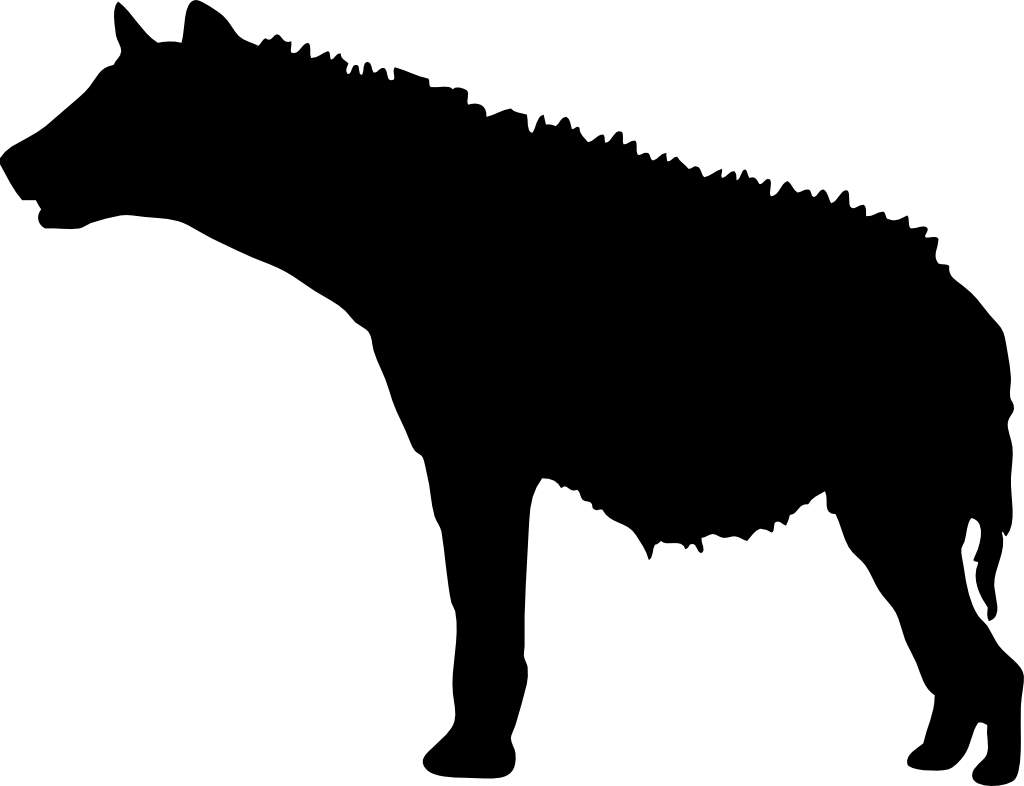

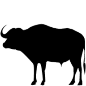

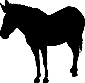

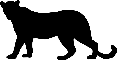

**Figure S7.** Marginal total effect of site-level *Opuntia* percentage cover (standardised) on the proportion of detections occurring at night for: **A)** olive baboon, **B)** vervet monkey, **C)** elephant, **D)** buffalo, **E)** dik-dik, **F)** impala, **G)** kudu, **H)** giraffe, **I)** Grevy’s zebra, **J)** plains zebra, **K)** spotted hyena, and **L)** leopard. The models assume that *Opuntia* indirectly affects occupancy through altering the composition of the native plant community; for model structure, see Figure 2 in main text. Shaded areas represent 89 compatibility intervals for the January-April (light green) and October-November (purple) seasons. Black lines indicate posterior median marginal effects for January-April under a new moon (―) and full moon (‧ ‧ ‧), and October-November under a new moon (– – –) and full moon	(– ‧ –).

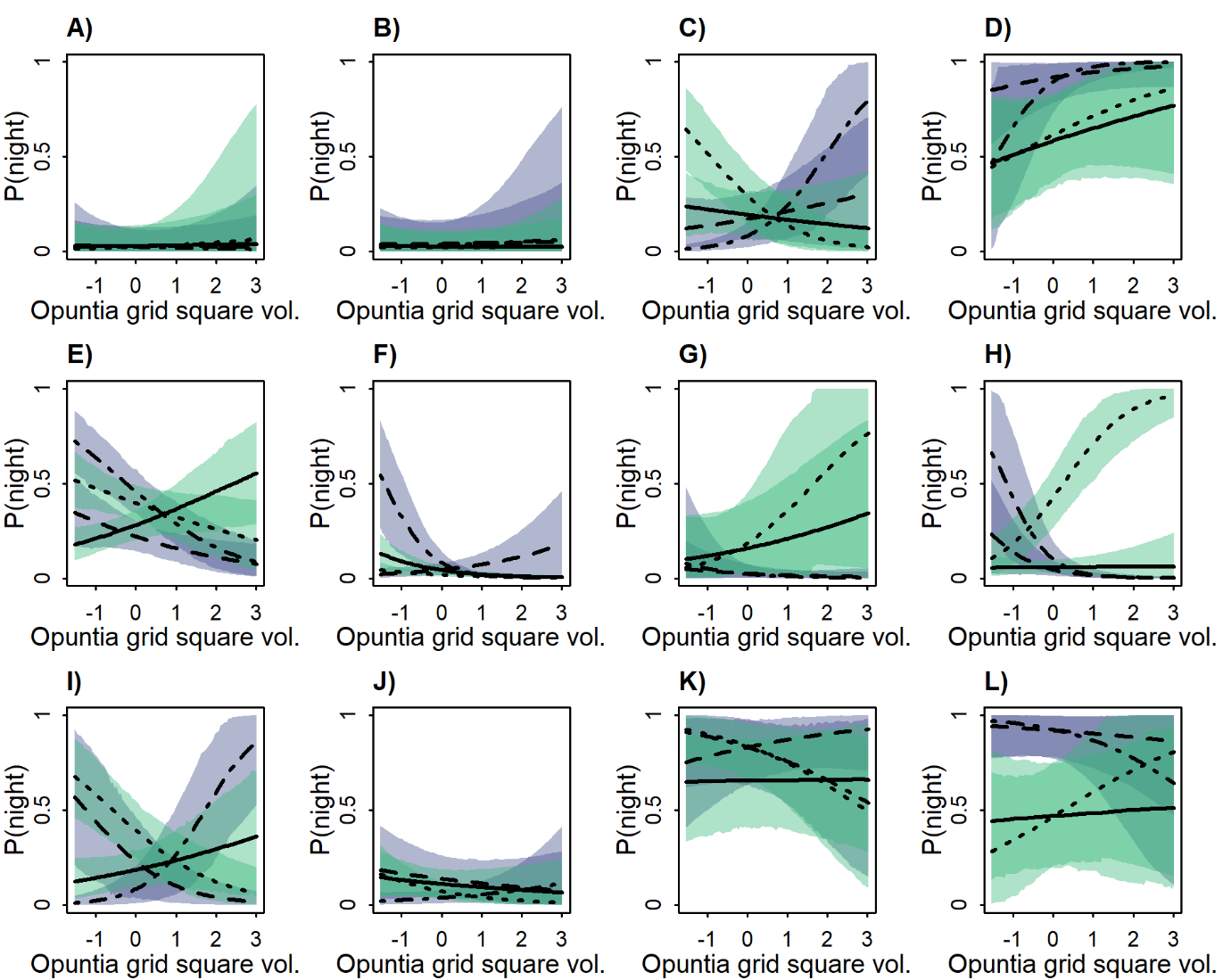

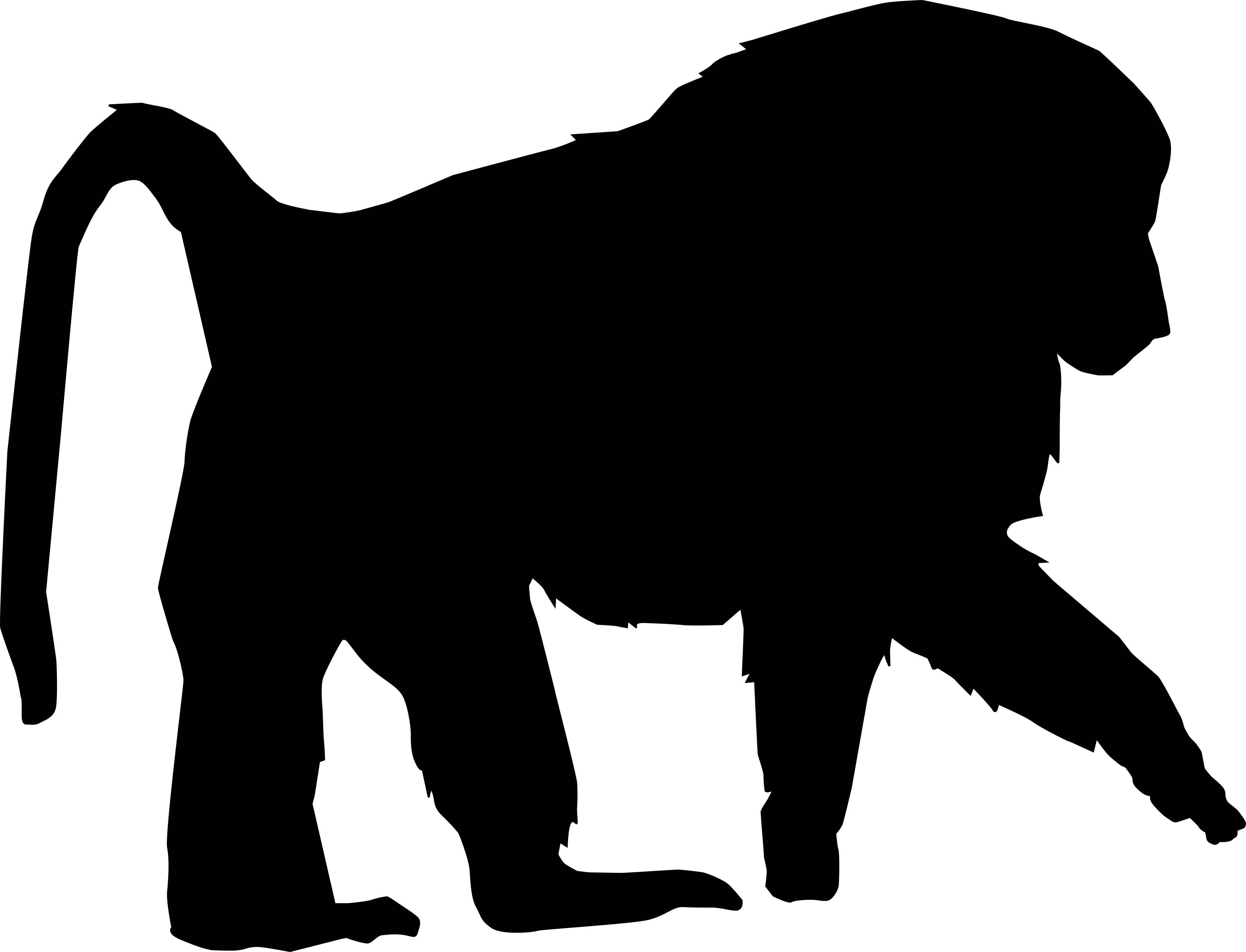

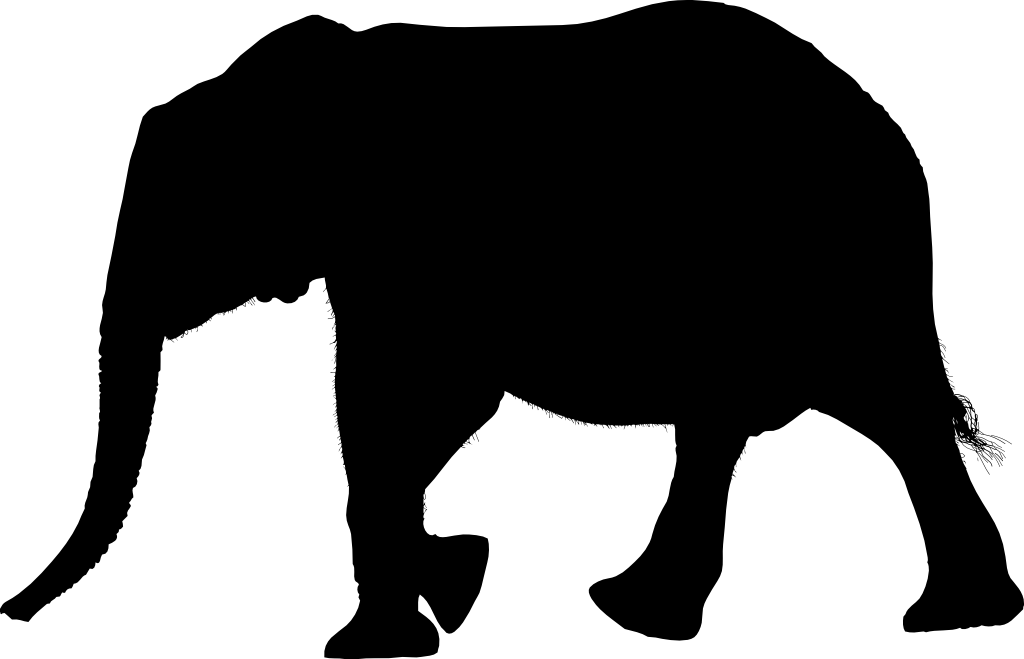

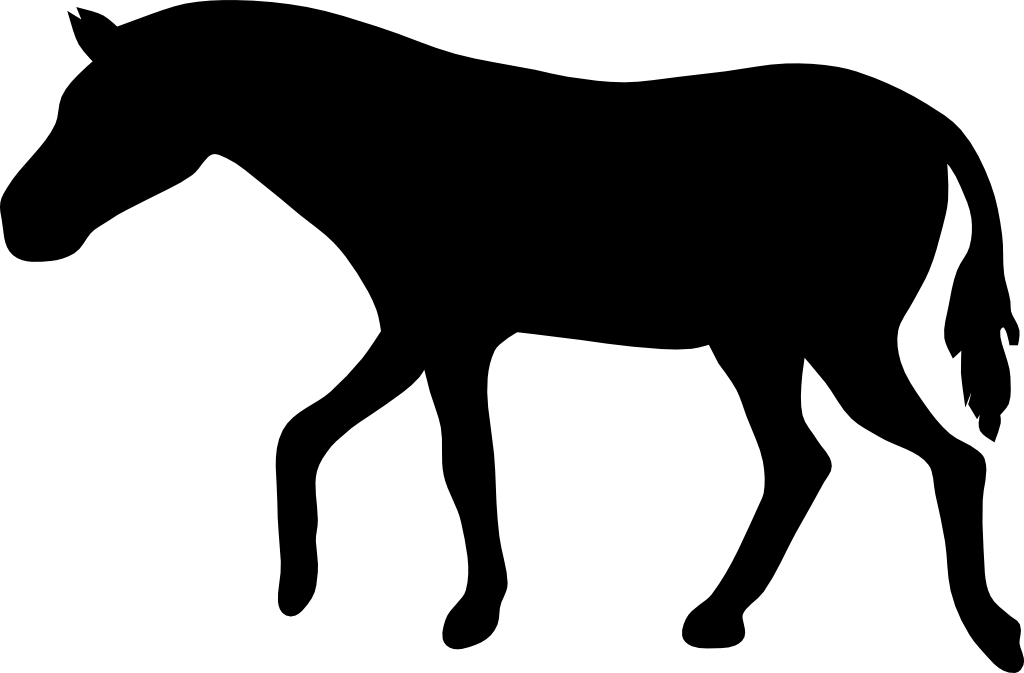

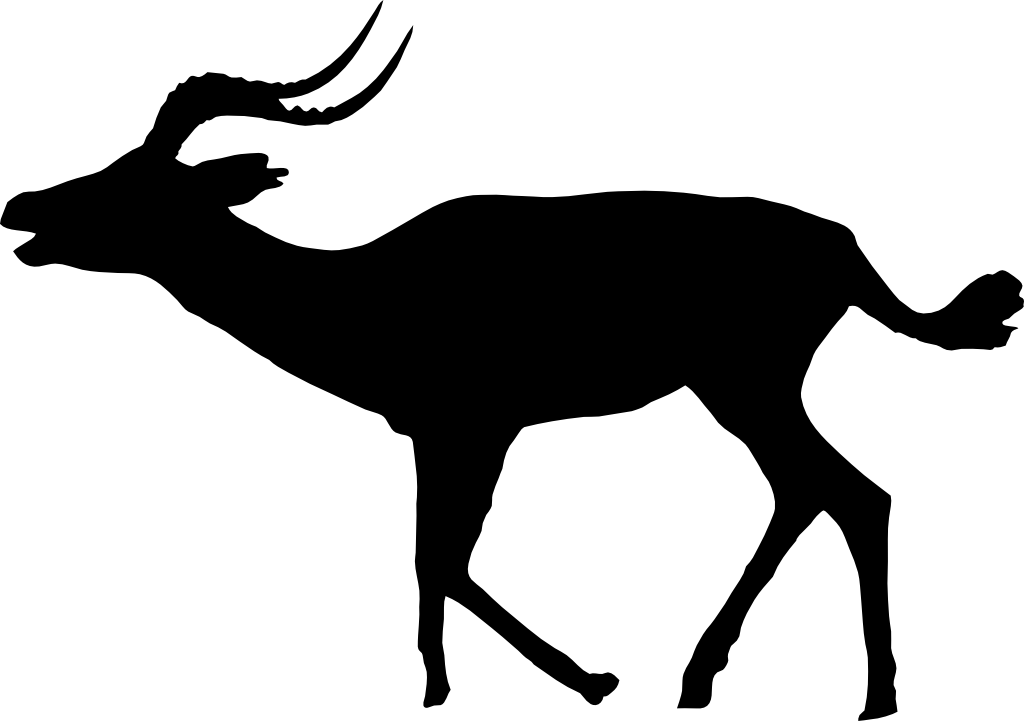

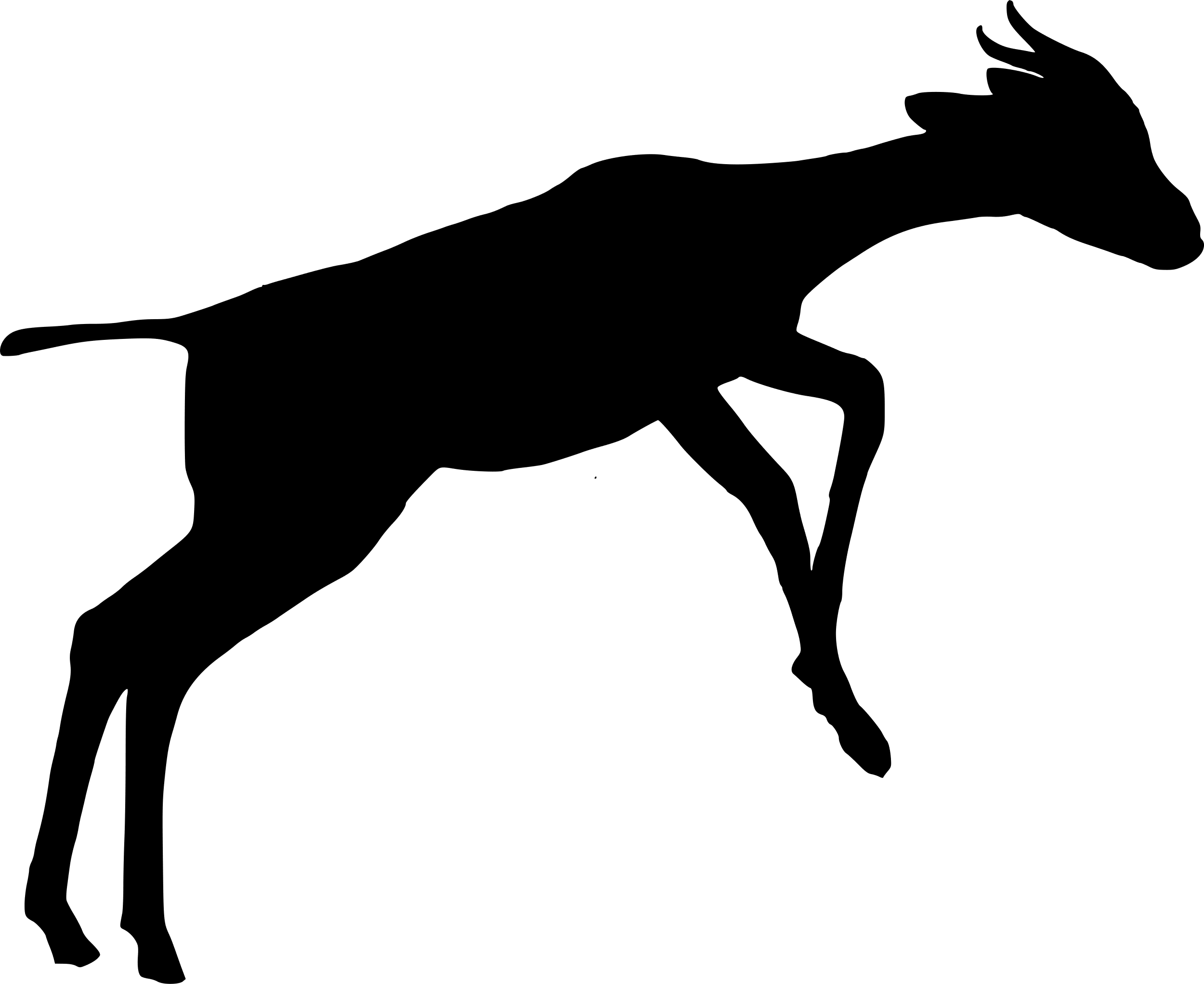

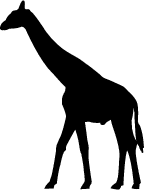

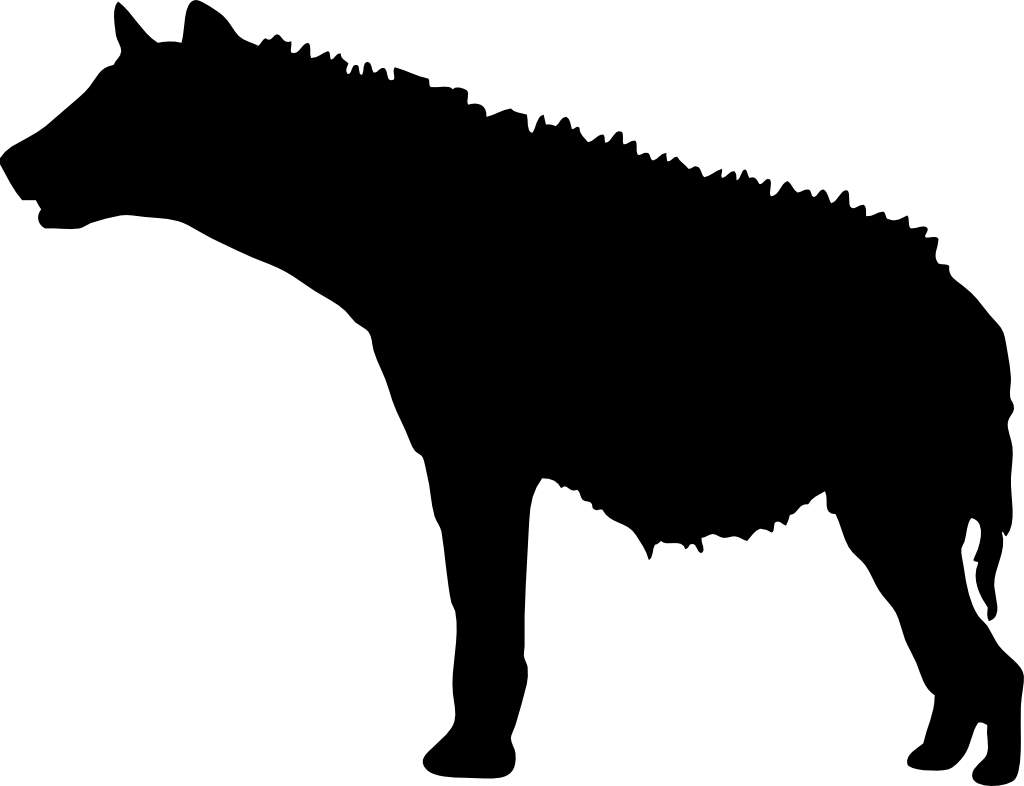

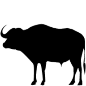

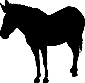

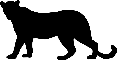

**Figure S8.** Marginal total effect of grid square-level *Opuntia* volume (standardised) on the proportion of detections occurring at night for: **A)** olive baboon, **B)** vervet monkey, **C)** elephant, **D)** buffalo, **E)** dik-dik, **F)** impala, **G)** kudu, **H)** giraffe, **I)** Grevy’s zebra, **J)** plains zebra, **K)** spotted hyena, and **L)** leopard. The models assume that *Opuntia* indirectly affects occupancy through altering the composition of the native plant community; for model structure, see Figure 2 in main text. Shaded areas represent 89 compatibility intervals for the January-April (light green) and October-November (purple) seasons. Black lines indicate posterior median marginal effects for January-April under a new moon (―) and full moon (‧ ‧ ‧), and October-November under a new moon (– – –) and full moon	(– ‧ –).
